# The shifting dynamics of ancestry and culture at a post-Roman crossroads

**DOI:** 10.64898/2025.12.12.693924

**Authors:** Deven N. Vyas, István Koncz, Tina Milavec, Tamara Leskovar, Norbert Faragó, Yijie Tian, Maja Bausovac, Balázs Gusztáv Mende, Paolo Francalacci, Uroš Bavec, Helena Bešter, Angela Borzacconi, Barbara Brezigar, Ronny Friedrich, Andrej Gaspari, Caterina Giostra, Zdravka Hincak, Špela Karo, Ana Kruh, Phil Mason, Alessandra Modi, Matic Perko, Rita Radzevičiūtė, Rok Ratej, Paola Saccheri, Anna Szécsényi-Nagy, Luciana Travan, Katarina Udovč, Stefania Vai, Bernarda Županek, David Caramelli, Johannes Krause, Walter Pohl, Tivadar Vida, Patrick J. Geary, Krishna R. Veeramah

**Affiliations:** Department of Ecology and Evolution, Stony Brook University, Stony Brook, NY 11794, USA; Institute of Archaeological Sciences, Department of Humanities, Eötvös Loránd University, Budapest 1088, Hungary; Department of Archaeology, Faculty of Arts, University of Ljubljana, Aškerčeva cesta 2, 1000 Ljubljana, Slovenia; Regional Museum of Celje, Trg Celjskih knezov 8, 3000 Celje, Slovenia; Institute of Archaeogenomics, HUN-REN Research Centre for the Humanities, Budapest 1097, Hungary; Dipartimento di Scienze della Vita e dell’Ambiente, Università di Cagliari, Cagliari 09126, Italy; Zavod za varstvo kulturne dediščine Slovenije, Poljanska cesta 40, 1000 Ljubljana, Slovenia; PJP d.o.o., Trg Alfonza Šarha 1, 2310 Slovenska Bistrica, Slovenia; Museo Archeologico Nazionale di Cividale del Friuli, Piazza Duomo 13, 33043 Cividale del Friuli (UD), Italy; Institute for the Protection of Cultural Heritage of Slovenia, Regional unit Nova Gorica, Cesta 5. maja 5a, 5000 Nova Gorica, Slovenia; Curt-Engelhorn-Center Archaeometry D6 3, 68159 Mannheim, Germany; Dipartimento di Storia, Archeologia e Storia dell’Arte, Università Cattolica del Sacro Cuore, Largo A. Gemelli, 1, Milano, 20123, Italy; Faculty of Humanities and Social Sciences, University of Zagreb, Ivana Lučića 3, 10000 Zagreb, Croatia; Institute for the Protection of Cultural Heritage of Slovenia, Centre for Preventive Archaeology, Poljanska cesta 40, 1000 Ljubljana, Slovenia; Goriški muzej, Grajska cesta 1, 5000 Nova Gorica, Slovenia; Dipartimento di Biologia, Università degli Studi di Firenze, Via del Proconsolo 12, Firenze, 50122, Italy; Skupina STIK, Cesta Andreja Bitenca 68, 1000 Ljubljana, Slovenia; Department of Archaeogenetics, Max Planck Institute for Evolutionary Anthropology, Leipzig 04103, Germany; Institute for the Protection of Cultural Heritage of Slovenia, Regional unit Maribor, Slomškov trg 6, 2000 Maribor, Slovenia; Department of Medicine, Section of Anatomy and History of Medicine, University of Udine, P.le Kolbe 3, 33100 Udine, Italy; MTA-BTK Lendület ‘Momentum’ Bioarchaeology Research Group, Budapest, Hungary; Museum and Galleries of Ljubljana, Gosposka ulica 15, 1000 Ljubljana, Slovenia; Institute for Medieval Research, Austrian Academy of Sciences, Vienna 1020, Austria; Institute for Austrian Historical Research, University of Vienna, Vienna 1020, Austria; School of Historical Studies, Institute for Advanced Study, Princeton, NJ 08540, USA

**Keywords:** paleogenomics, migration, kinship, burial archaeology, community formation, Early Medieval Europe

## Abstract

The collapse of the western Roman Empire in the 5th century created a period of geopolitical upheaval, driven by Barbarians dispersing into the former Empire and reshaping post-Roman communities. While there is now evidence for a major genetic impact of these migrations into specific regions of Europe, it is unknown whether these changes in ancestry were uniform across the continent. We investigate present-day Slovenia, a crucial crossroad connecting the Roman East and West and the gate to Italy during the Langobard invasion. We conducted paleogenomic and isotopic analyses of 410 individuals from 21 sites across Slovenia and Cividale (Italy), establishing a longitudinal transect, spanning eight centuries. During Late Antiquity, despite changes in burial artifacts, kinship practices, and settlement structures reflecting a shift in culture, we find high levels of genetic continuity with the local Late Roman population and reduced mobility. However, demographic turnover began during the 8th century, when communities with northeastern European ancestry and distinct cultural practices entered the region, gradually advancing westward over the span of three centuries, replacing the local populations. This shows that cultural change in post-Roman Europe could be decoupled from genetic change in transit zones, demonstrating a dynamic spatiotemporal process across the continent.

## Introduction

The decline and eventual collapse of the Western Roman Empire beginning in the 5th century CE resulted in a political transformation across Europe that opened the way for so-called “Barbarian” groups to migrate into its former territories and establish new political entities^1–5^. These groups came to characterize the political landscape of Western and Central Europe in the following centuries. Historical and archaeological research still debates the extent to which these changes resulted from, or only coincided with, mass movements, gradual diffusion, cultural transfer or outright invasion, as many elements in this process are unclear from the available textual or material sources^6–17^. Recent paleogenomic studies have found evidence of various types of population movements both inside and outside of the borders of the former empire, supporting historical and archaeological narratives of migrations into the territory of the former Western Roman Empire^1,18,19–20^.

The territory of present-day Slovenia played an important role connecting the eastern and western halves of the Roman Empire, and as such was crucial during the civil wars and various invasions of the 4th-6th c. CE^21–27^. With its well-developed military and civic infrastructure, this territory represents the easiest crossing of the Alps and access to the Italian Peninsula, connecting it to the Pannonian and Balkan Roman provinces. After the region’s abandonment by Roman civil and military administration in the 5th c. CE, these same routes were used by Goths and Langobards moving into Italy, and later probably by Slavic groups settling in the area^28^. These events caused profound cultural and social transformations observable in repeated relocations of settlements and changes in material culture and burial customs. The nature of these transformations remains the focus of contrasting theories in historical and archaeological narratives^29–33^.

Through the comprehensive paleogenomic and isotopic analyses of sites ranging from Late Roman Period (LR, 3rd to early 5th c. CE, n=163 genomes) to Late Antiquity (LA, late 5th to 7th c. CE, n=141) and through to the early Medieval period (EM, 8th to 11th c. CE, n=66 genomes) we investigated the following questions: Is there continuity between the LR towns and the much smaller dislocated LA communities? How did the various population movements described by the written sources - the migration of the Goths and Langobards in the 5-6th c. and the expansion of the Slavs from the 7th century onwards - affect the population structure in the region? In general, do changes in settlement patterns and material culture coincide with observable changes in the genetic structure of the population?

We find that during the 5th-6th century CE this region exhibits remarkable continuity with the core of the preceding LR population despite the historical sources reporting several movements of peoples in and out of the region^21–27^, while the only observable influx of newcomers is restricted to certain smaller communities dated to the time of the Ostrogothic Kingdom. Based on the investigated sites, the migration of the Langobards did not have a clear impact on the genetic background of the population: the increase in Northern European genetic background observed in Pannonia (the point of departure) and in Italy (the destination) in previous paleogenomic studies is not observable in Slovenia^1,18,34,35^. However, we do see a significant increase in Northeastern European ancestry after the 8th-9th century, suggesting a considerable influx of new groups coinciding with the emergence of small communities described as Slavic across the region. Our results also identify differences between four subregions, with evidence of population continuity into the 9th c. CE in the westernmost part of the territory, closest to Italy. We highlight social differences between communities with similar genetic backgrounds, proposing a complex and nuanced picture of the impact of population movements in the region rather than a simple direct correlation between genetic changes and traditional historical narratives.

### Sites and sampling

We constructed a spatial and temporal paleogenomic and isotopic sampling transect along the traditional routes connecting Italy to the east that have been historically linked to several movements of various barbarian groups (Ostrogoths, Langobards, Slavs) between 300-1100 CE. A number of sites (n=21) and samples (n=410) cover the area between northern Italy, Pannonia and the Balkans and follow the changing settlement pattern between Roman urban centres to LA central sites and then to EM rural settlements (**Figure 1; Table S1**). The 21 sites include cemeteries of late Roman civil and possibly military centers (multiple sites from Celeia, Gosposvetska cesta from Emona, Ajdovščina Castra, and Črnomelj) from the 4th-5th centuries; smaller lowland sites such as Miren and Dravlje and cemeteries connected to so-called hilltop sites (Rifnik, Vrtovin_LA, Solkan, Vrajk, Gradec, Zidani gaber) dated generally to the time of the Ostrogothic and Langobard kingdoms during the 5-7th centuries; and 8th-9th century rural cemeteries and grave groups (Hrušica, Camberk, Ledine, Gojače, Vrtovin_EM) along with an 11th c. cemetery from Ajdovščina-Šturje (**Section S1**). Outside of Slovenia we also investigated two Langobard-period cemeteries in Cividale (Ferrovia n=24 and San Mauro n=16). Celeia, Emona, Vipavska dolina, and Cividale represent a geographical transect following the main Roman road from Italy to the Pannonian provinces, while the Dolenjska and Bela krajina region to the southeast is located next to the Roman road that connected Italy with the Balkans and Constantinople. The excavated LR sites only represent the latest parts of the original cemeteries as some have over 300 excavated graves, and as such comprehensive sampling was not possible. However, in the case of LA and EM sites, we sampled all available individuals. All samples underwent 1240k single nucleotide polymorphisms (SNPs) capture and sequencing^36–38^. Altogether, the investigated sites yielded 410 samples for aDNA analysis (n=389 with 1240k SNPs coverage ≥ 0.1× and overall mean 1240k SNPs coverage of 3.54×) and 329 samples for ^87^Sr/^86^Sr analysis. These individuals were analyzed alongside a large set of Eurasian and African individuals from similar time periods (**Table S2; Section S2**). To complement the results of genetic and isotopic analyses and to test and improve archaeological chronology of the investigated sites, we also radiocarbon-dated 73 samples (**Table S3**).

**Figure 1.**
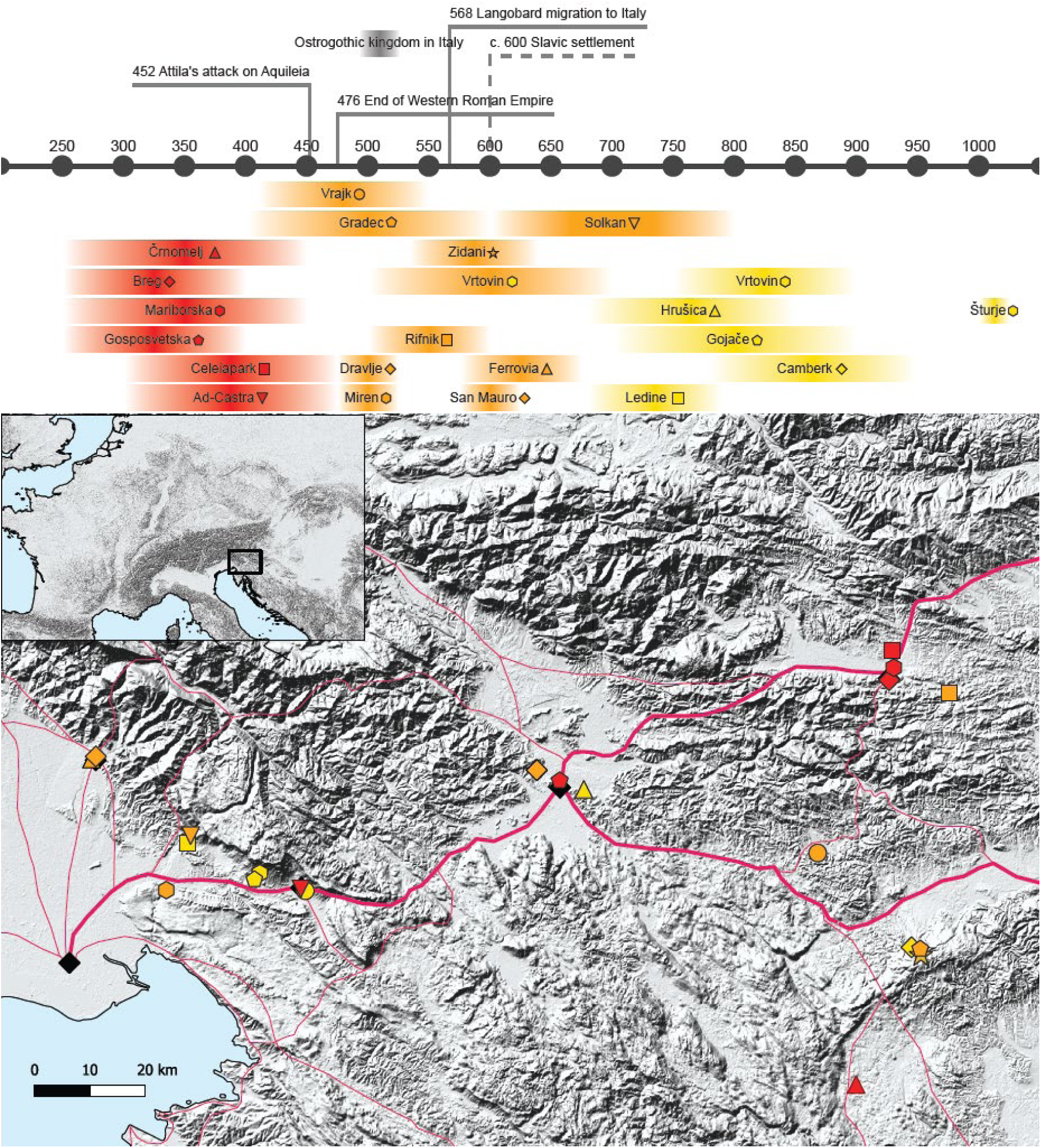
Map of the sites, color-coded based on time period, alongside the timeline.

### Population continuity and cultural transformation between Late Roman period and Late Antiquity

LR centers (Castra, Celeia, Emona) were gradually abandoned, or at least lost their administrative and logistical roles, by the mid-5th century CE^39,40^. In the late 5th or early 6th c. CE so-called late antique hilltop settlements appeared in geographically protected locations (Gradec, Rifnik, Solkan, Vrajk, and Vrtovin_LA) as the result of political instability, supposedly built by the people from the abandoned towns and countryside^41^. They also provide the main archaeological evidence of the passing and possible settlement of the Langobards in this region^42^. Only a few sites can be dated to the period in-between, these being attributed to small communities in strategic locations and interpreted as being founded by new arrivals to maintain control^39,39,43^. Cemeteries connected to the LA sites show significant changes in burial customs and material culture^44^ compared to the LR period. Through the comprehensive paleogenomic and isotopic sampling of the above-mentioned sites, we investigated how shifts in political power and settlement location impacted the population in the region. In particular we aimed to determine whether evidence of changes in settlement structure and material culture in the region were accompanied by population change and/or replacement, or were the result of cultural transformation.

We first used principal component analysis (PCA) (**Figure 2**) and model-based clustering analysis via fastNGSadmix^45^ (**Extended Data Fig. 1**) to compare patterns of genomic ancestry between communities from the LR-period centers and from LA lowland and hilltop sites ^45–48^. For PCA we compared ancient samples to a modern European reference set^49,50^, while for fastNGSadmix we decomposed them into ancestry components represented by a set of ancient individuals that form eight reference panels encompassing penecontemporaneous (c. 4th–8th century CE) Eurasian genetic diversity^18,51^ (CASIA: Central Asia, EASIA: East Asian, MEDEU: Mediterranean Europe, NAFRICA: North African, NGBI: Northern Germany/British, SASIA: South Asian, SCAND: Scandinavia, and SUBSAHARAN: sub-Saharan Africa). For fastNGSadmix, we additionally used data from the 1000 Genomes Project to form a modern reference panel^52^. Additionally, we used qpWave^36,53^ (**Figure 3**) to formally test whether individuals could be genetically modeled as descending from earlier communities (i.e. to test for the degree of population continuity). PCAs were also conducted using modern pan-Eurasian AADR populations as the modern reference set^53–62^, which indicates that the communities we studied almost exclusively overlap modern Europe genetic diversity (**Extended Data Fig. 2; Table S4**). We additionally conducted fastNGSadmix analyses using 1000 Genomes Project populations as modern references, and found that these results are largely consistent with those generated from the ancient panels (**Extended Data Fig. 1**).

**Figure 2.**
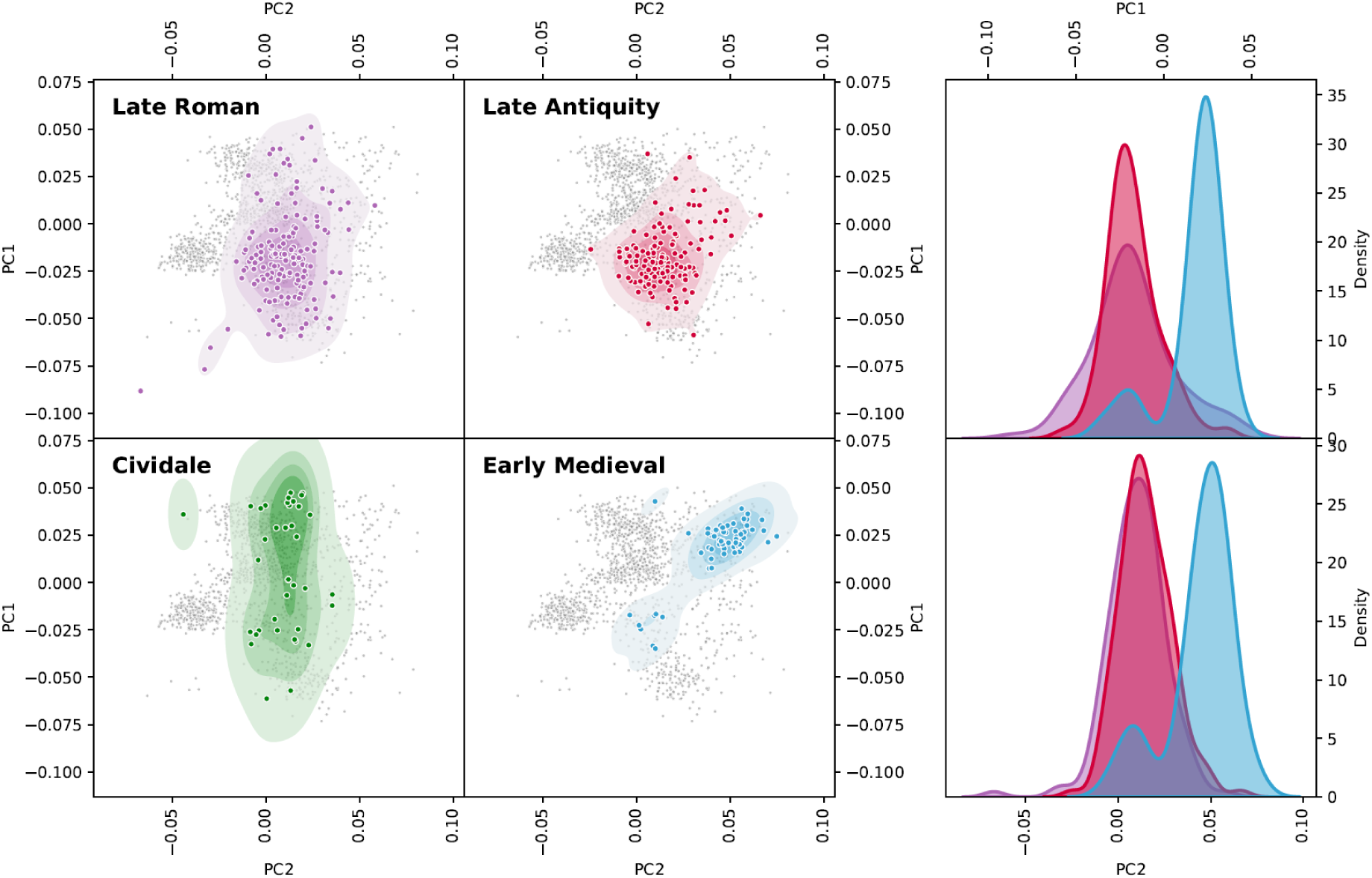
PCA of the LR, LA, and EM populations from modern Slovenia, alongside the 6th-7th century CE sites of Ferrovia and San Mauro from Cividale, Italy. Individuals from the POPRES^49,50^ are used as the reference panel and are plotted in greyl. Additionally, Kernel Density Estimation plots of PC1 and PC2 are depicted for the LR, LA, and EM plots.

**Figure 3.**
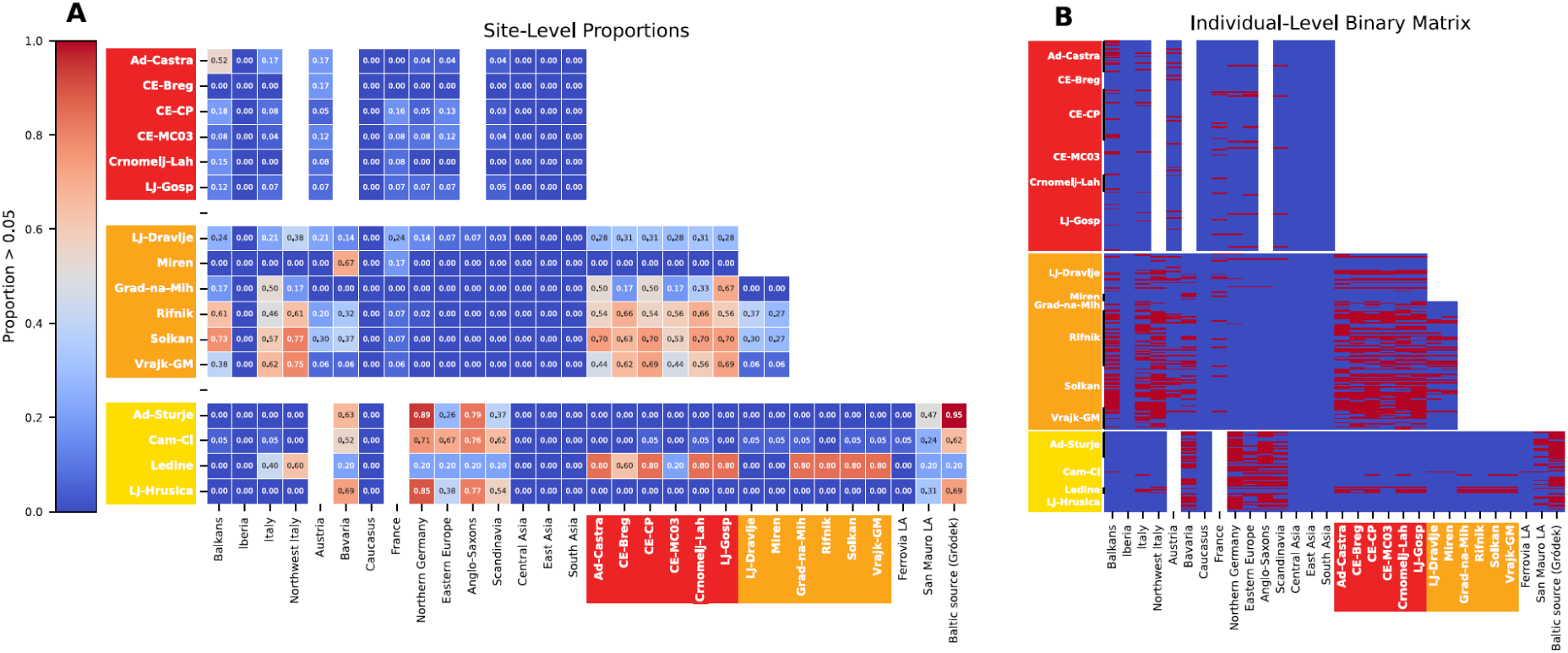
qpWave analysis. **A** presents qpWave results testing whether communities could be modeled by penecontemporary Europeans, 3rd to 10th century Asian populations, and Slovenian communities from the previous time period. Redder numbers indicate more of the test community (y-axis) can be modeled by the reference population (x-axis). **B** presents the same information at the individual level. Slovenia communities are color-coded on both x- and y-axes based on time period based on Figure 1. Reference populations are adjusted for each time period, to only reflect individuals from that time period and prior. Gapped columns reflect cases where reference populations that did not have data covering a certain time period (e.g., we lack Bavarian data from the LR period).

PCA and model-based clustering analyses showed that in the LR period, the genetic diversity in the investigated area is centered primarily around individuals from southern and southeastern Europe (as represented by both modern and penecontemporaneous European reference samples). In model-based clustering analyses, we find that the three European components—MEDEU, NGBI and SCAND—make up most of the ancestry in these communities. The southern European MEDEU component accounts for 68% of the total ancestry (when summing ancestry components from all analyzed individuals), while the northern European NGBI and SCAND components together represent 30% of total ancestry, with the Asian and African components making up the small remainder (**Extended Data Fig. 1**). However, there are a number of outliers with non-European ancestries from Africa and Asia in multiple communities, probably as the result of the long-distance networks of the Roman Empire. This genetic diversity is paralleled by the diversity in strontium isotope results, pointing to the consumption of food sourced from geologically diverse regions (**Figure 4, Supplemental Figure S4)**. The ^87^Sr/^86^Sr results show a wide range of values within single areas such as Emona, Celeia, Castra, and Črnomelj. This isotopic variability suggests mobility at regional or supra-regional scales, likely reflecting the movement of individuals in and out of the Roman towns and the circulation of resources across the regions (**Figure 4**, **Section S2**).

**Fig 4.**
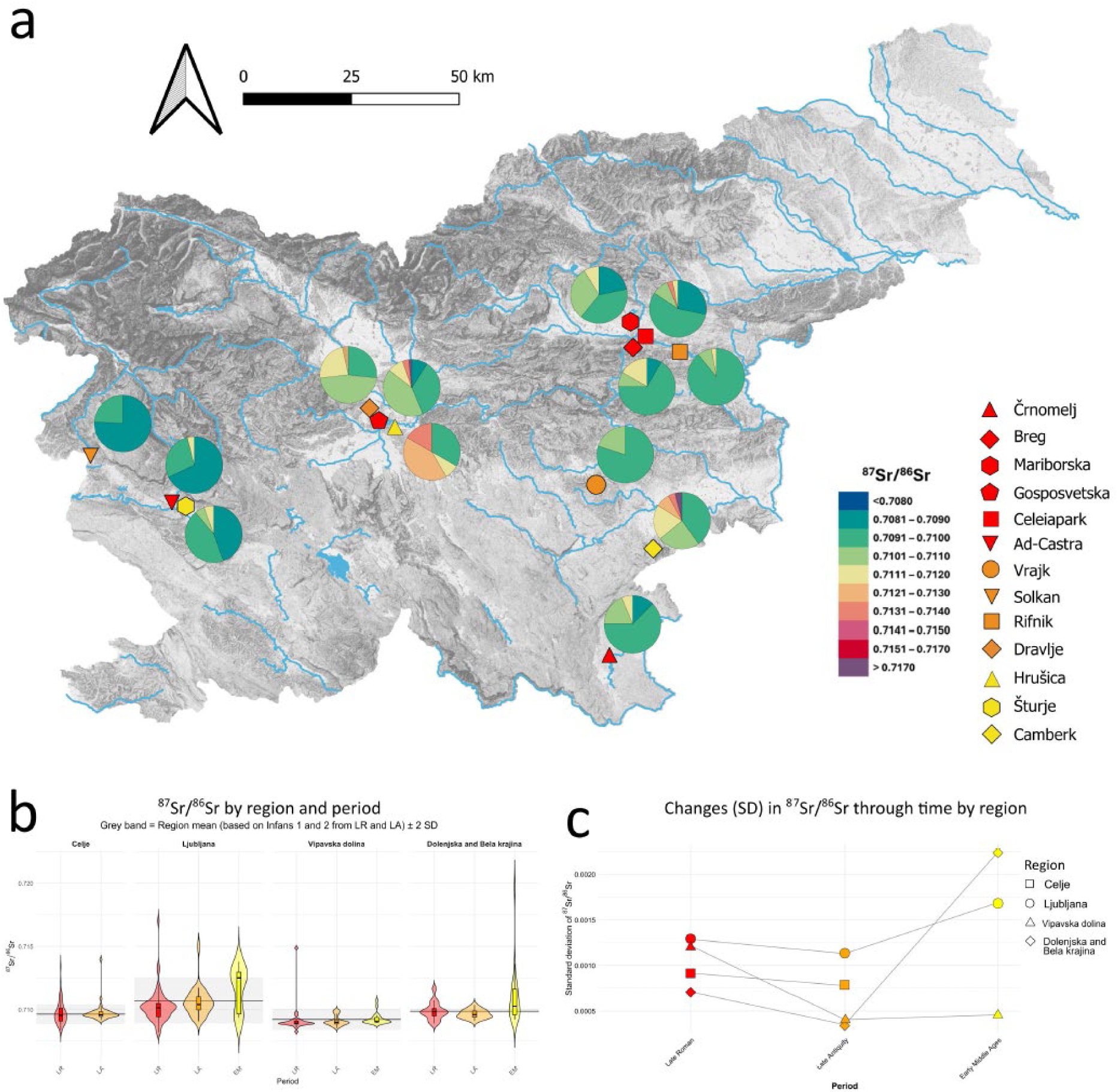
Results of ^87^Sr/^86^Sr analysis: **A** Map showing the distribution of absolute ^87^Sr/^86^Sr values among the investigated sites with more than 10 samples. Colors indicate the three main periods (LR, LA, EM); **B** Violin and boxplots show the ranges of ^87^Sr/^86^Sr values grouped by region and period. Grey bands indicate a presumed ‘local’ range based on *infans* values from LR and LA sites; **C** Changes in standard deviation (SD) of the ^87^Sr/^86^Sr values are shown across time in the investigated regions.

During Late Antiquity, despite the migrations described in written sources and the changes in material culture and settlement locations observed in the archaeological record, we do not see a substantial change in the primary composition of genetic ancestry, though the population becomes significantly more homogenous overall. Genetic diversity during the LA period as measured by variance along the first principal component is less than half that observed during the LR period (5.83×10^-4^ v. 2.42×10^-4^) while maintaining the same PC1 mode, and the mode and variances for PC2 are very similar (LR: mode: 1.13×10^-2^; variance: 2.38×10^-4^; LA: mode: 1.12×10^-4^; variance: 1.95×10^-4^) (**Table S5**). A factor analysis^63,64^ based on imputed genotypes for a subset of unrelated Slovenia individuals with coverage >=1x— which unlike PCA does not rely on any external reference genomes and controls for sample age-related genetic drift—shows a very similar Factor 1 profile for the LR and LA samples (**Extended Data Fig. 3**), suggesting no major change in the profile of genomic ancestry between the two periods. To further test for similarity or difference between the communities, we conducted pairwise F*_ST_* analyses for the investigated communities, as well as for each region per time period and the whole time periods (**Extended Data Fig. 4**). The F*_ST_* values for Slovenian communities within and between the LR/LA periods were generally low with the exception of Crnomelj and Vrajk who presented a distinct profile from the other. Comparisons without Vrajk and Crnomelj ranged from 0.000-0.0020 and 21/28 comparisons were non-significant after Bonferroni correction (*Z:* -0.56-5.50; *p*: 3.8 × 10^-8^-0.97); comparisons including these sites ranged from 0.0000 to 0.0035 and only 3/17 comparisons were non-significant (*Z:* 1.70-10.80; *p*: 0-0.090) (**Table S6**).

The qpWave analyses generally support the model that the LA communities descend from the LR communities that abandoned the Roman towns (**Figure 3; Table S7**). Most individuals from the LA communities can be modeled as descending from one or more LR communities (75%, 96/128 tested individuals). The two exceptions to this trend are Dravlje and Miren. Dravlje can only be partially (27-33% of individuals) modeled via a single stream of genetic ancestry from the earlier local LR communities, while Miren cannot be modeled at all (0%) (**Figure 3; Table S7**). In total, 23 of the 32 LA individuals that could not be modeled by any LR community came from Dravlje or Miren. Overall, there is an increase in the percentage of the population clustering with southern/southeastern Europe with less northern European and non-European outliers, although slightly different patterns emerge between lowland and hilltop sites.

The lowland communities of Miren and Dravlje, generally dated to the time of the Ostrogothic Kingdom (late 5th-first half of 6th c. CE)^65,66^ (**Figure 1**), contain individuals with a notable amount of Central Asian genetic ancestry. In Miren five out of six individuals carry small amounts of Central Asian ancestry ranging from 7-12% (**Extended Data Fig. 1**). In Dravlje, an adult male (Dravlje_25) has 76% Central Asian ancestry and his son (LJ-Dravlje_20) has 44% with 33% Northern European ancestry, suggesting a recent admixture event. Five other individuals from Dravlje have Central Asian ancestry ranging from 12% to 35%. For those individuals from Miren and Dravlje who could not be modeled via the qpWave analysis described above, we subsequently performed a qpAdm analysis which allowed for two streams of genetic ancestry, one from LR communities and the other from Asian populations (**Section S3; Table S8**). In about half of these cases (11/23, 48%) we could model them as a mixture of an LA community and Central Asians. A smaller subset of these 11 could also be modeled with East Asians (3) or South Asians (6).

The individuals with substantial Central Asian ancestry from Dravlje were buried at the eastern edge of the site, forming their own spatial cluster (**Figure 5**). The strontium isotope values for these individuals with Central Asian ancestry fall within the local range, suggesting that they consumed food grown in the region and likely grew up locally. Similarly, their associated grave goods do not differ significantly from the rest of the community and with the exception of Dravlje_25, their burials are generally poorly furnished. Dravlje_25 was buried with an elaborate copper-alloy buckle decorated with garnet inlays, a type that has its known parallels from Western Europe, including contexts dated to the Ostrogothic Kingdom in Italy^67^. An object with similar parallels, a buckle with chip-carved rectangular plate, was also found in the burial of a female with mostly Northern European ancestry and local strontium values (Dravlje_01), demonstrating that there is no clear correlation between artefact types generally interpreted as ‘Ostrogothic’ and Central Asian genetic ancestry.

**Figure 5.**
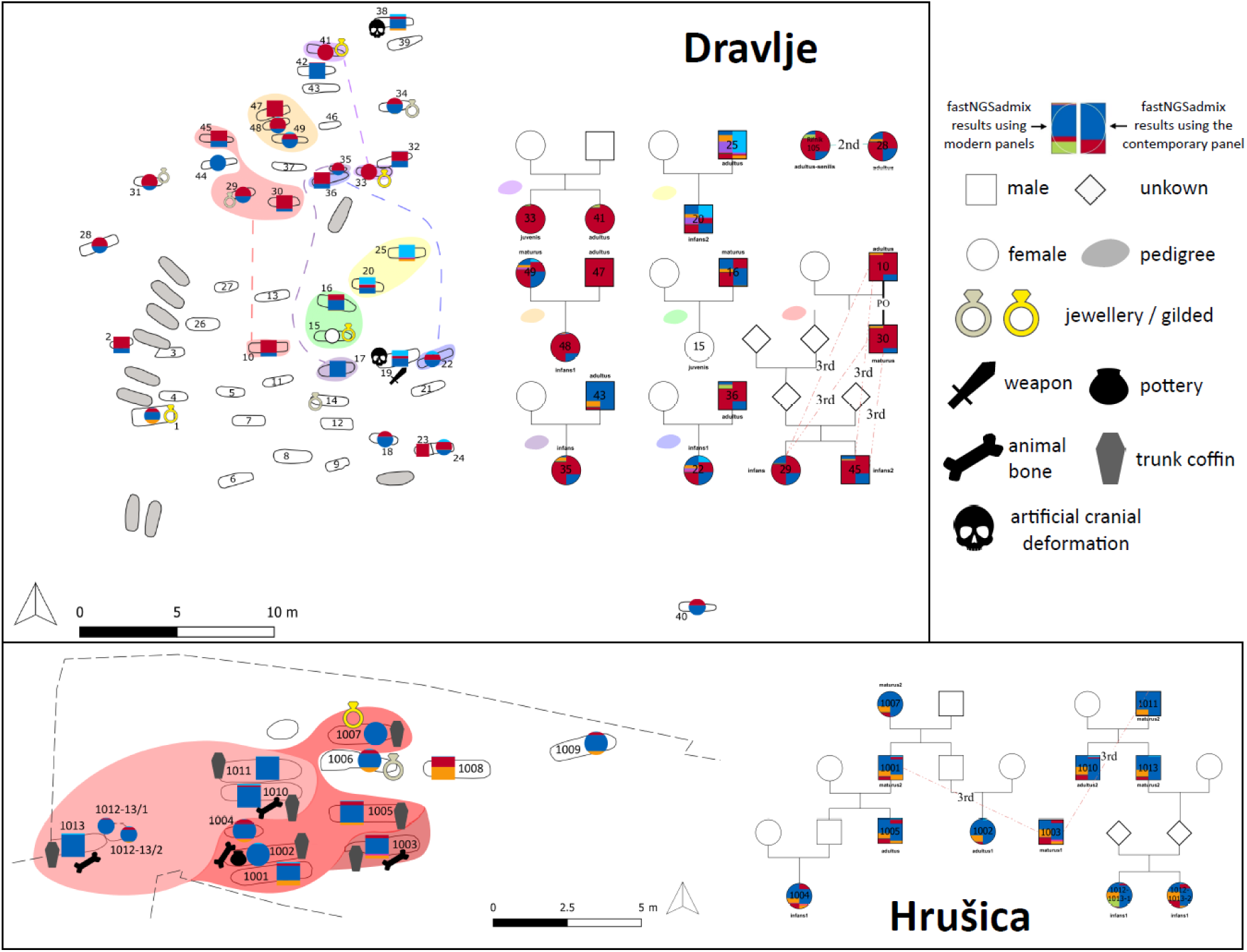
Cemetery maps of Dravlje and Hrušica. Burials with sufficient genetic data were marked male (square) or female (circle), which were colored based on their estimated genetic ancestry using fastNGSadmix (see Extended Data Figure 1). Pedigrees are included and color overlayed over the cemetery maps. The pedigree from Hrušica has two lines that are joined by a single individual and thus is in three shades. Archaeological artifacts as well as individuals with artificial cranial deformation are also labeled. For fastNGSadmix, using the modern 1000 Genomes Project panel, CEU+GBR is depicted in blue, FIN in orange, EAS in purple, IBS in light green, SAS in pink, YRI in dark green, and TSI in red. For the contemporary panel, CASIA is depicted in light blue, EASIA in purple, MEDEU in red, NAFRICA in teal, NGBI in blue, SASIA in pink, SCAND in orange, and SUBSAHARAN in green.

Artificial cranial deformation (henceforth ACD)—a foreign, most probably non-Roman tradition in the region that spread rapidly during the Hunnic period among both males and females, but also disappeared fairly quickly and was no longer practiced by the end of the 6th century^68^—appears at both sites (Dravlje_19, Dravlje_38, Miren_01, Miren_07), but only among individuals with Central Asian genetic ancestry, consistent with findings from penecontemporaneous populations from Bavaria and Pannonia^50,51^. All four individuals present local strontium isotope values, indicating a local upbringing (**Table S1**).

In comparison to the lowland communities, there is no substantial non-European genetic ancestry observed at the hilltop sites, with the exceptions of five individuals who retain 4%-9% Central Asian ancestry at Rifnik (**Extended Data Fig. 1**). Interestingly, about a third of individuals from Rifnik or Solkan could be modelled by the the slightly older LA lowland sites of Dravlje and Miren (**Figure 3**) and we even observed 2nd degree relatedness between Dravlje_28 and Rifnik_105, demonstrating the close links existed between the lowland and hilltop communities (**Table S9**). On the other hand, at each of the LA sites, between 40% and 70% of individuals can be modeled by Late Roman sites, suggesting the possibility of continuity among them. Considering all tests, 96/128 LA individuals could be modeled with at least one Slovenian LR site.

In contrast to the LR period sites, the strontium isotope values across all LA sites exhibit a very narrow range, suggesting that most individuals were likely raised locally (**Figure 4**). The site of Miren shows a slightly broader overall range; however, this variation is primarily driven by two individuals, while the majority cluster closely around the local baseline. Only three hilltop individuals stand out from the local Sr range and seem to have access to food from different regions (Rifnik_34, Rifnik_91 and Solkan_53), all of whom were adults and not part of large pedigrees. This implies that they likely spent their childhoods elsewhere, but overall mobility appears significantly lower than in the LR period.

Genetic analyses also shed light on the role of biological relatedness in these communities using lcMLkin (**Table S9**). LR cemeteries contained fewer individuals with biological relatives (10-46%) than lowland and hilltop LA sites (50-66%). In the Roman period sites, pedigrees (groups of biologically closely related individuals) are generally smaller (usually containing 2-4 individuals) and primarily restricted to two generations. However, in LA sites (particularly Rifnik and Solkan) we find larger pedigrees that extend to up to four generations, while also retaining the small pedigrees found in the prior time period. For example, the two largest pedigrees at Rifnik and Solkan contain 8 and 17 individuals, respectively (**Table S9**). These differences could reflect that Roman society and concepts of family may have been less structured around biological relatedness or have not focused biological relationships in burial placement decisions, while biological kinship played a more significant role in the life and formation of later post-Roman communities. While our sampling is substantially less representative of the overall population from the much larger LR centers than it is to the LA communities, we sampled clusters of graves as opposed to random samples, and thus might expect to find relatives buried together if this was considered important in Roman burial customs. Correlation between burial customs and biological relatedness is observable in most LA sites: In Dravlje two sisters (Dravlje_33 and Dravlje_41) were buried with rich furnishings, in Rifnik members of a large pedigree of at least four generations had the most grave goods at the site, while in Solkan the three most richly furnished males in the largest pedigree at the site were cousins (Solkan_12, Solkan_31, Solkan_51), suggesting that these connections were not just biological, but represented meaningful social ties (**Section S1**).

In order to assess more distant relationships between the communities we also studied, we used ancIBD^69^ to identify shared identity-by-descent (IBD) tracts, both within our dataset and with previous published data (**Figure 6; Extended Data Fig. 5; Tables S2, S10-12**). The disconnect between Miren and Dravlje and the other sites is also supported by the IBD results, as these communities have very few external IBD connections within Slovenia: Miren has a single link to Solkan, while Dravlje has one to Celeiapark/Mariborska. Overall, we find that 16 intra-Slovenian LR-LA pairs share IBD connections (**Figure 6**).

**Figure 6.**
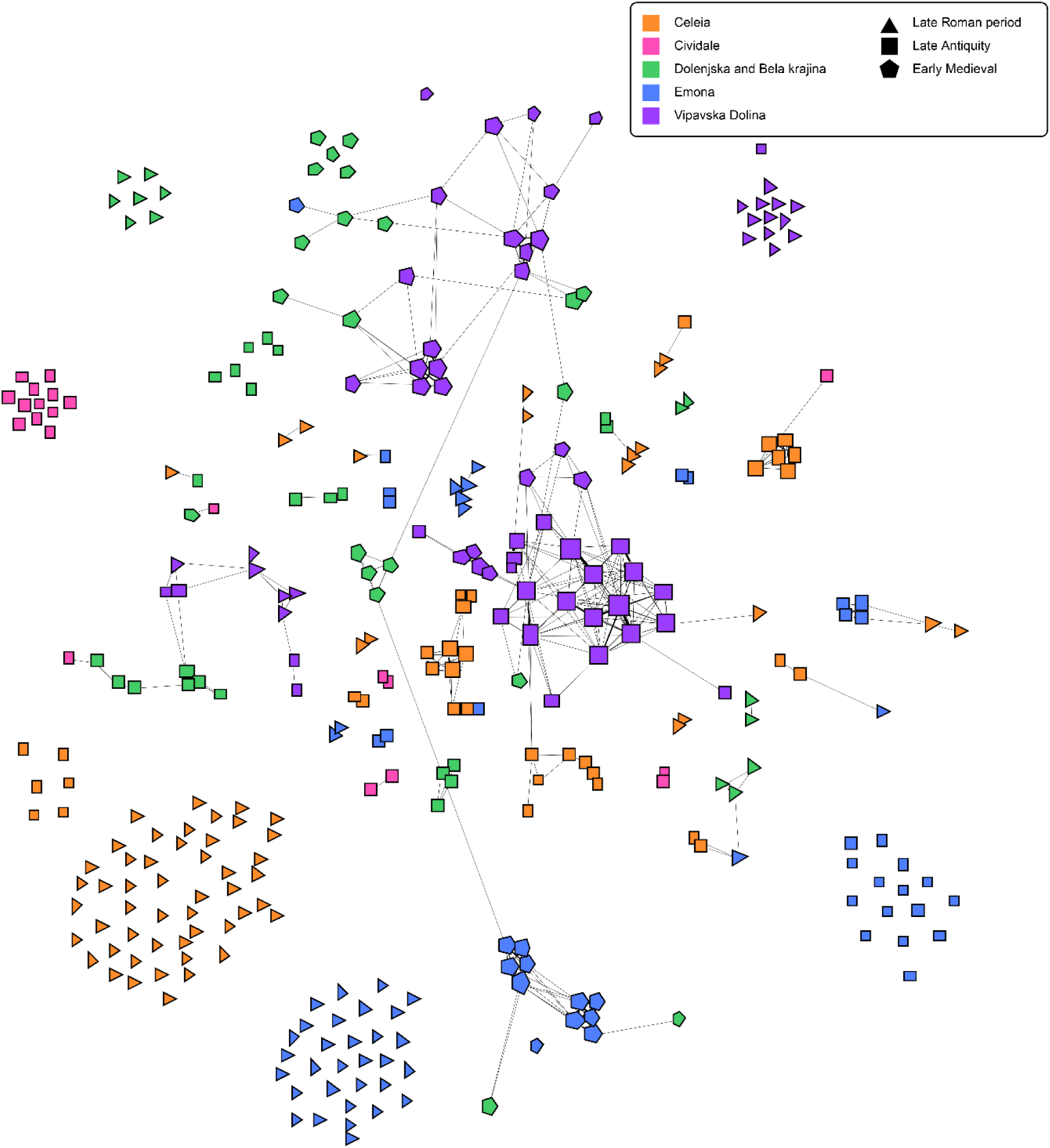
IBD connections between sites. Each node represents an individual. Nodes are colored by region and shapes are assigned by time period. The size of the node reflects the number of connections, and the width of a connection represents its strength. This network only reflects shared tracts greater than 20cM in length, as calculated using ancIBD^69^.

Our results show that despite various population movements mentioned by the written sources and the significant changes in political power, settlement, social structure and material culture, the genetic composition of the population remained fairly stable between the 4th to 7th/8th centuries CE, with the genetic heterogeneity decreasing over time. This trend is most strongly seen in hilltop sites such as Solkan and Rifnik; however, an exception is found in the lowland sites of the Ostrogothic period (Miren and Dravlje), with an introduction of a substantial amount of Central Asian ancestry. The appearance of this Central Asian ancestry together with foreign cultural traditions, such as ACD, as well as new artefact types suggest an influx of people into the territory. Similar artefact types and ACD appear together earlier in the Carpathian Basin during and after the Hunnic period (mid to late 5th century) and certain areas of the Balkans, and these areas could be considered as possible sources of origin based on historical sources as well^2,51,70^. Based on archaeological chronology and radiocarbon dating, these lowland communities do not replace the LR centers, but partly coexist with their latest phase and are probably related to the consolidation of power through the occupation of strategic locations.

### A migration without a genetic impact - the Langobards?

The migration of the Langobards is one of best historically and archaeologically documented population movements in early Medieval Europe. According to the written sources^12,71–73^, in 568 the Langobards led by their king, Alboin, left Pannonia (today’s Western Hungary) and moved to Italy, where their kingdom existed until the 8^th^ century, when it was finally absorbed by the Carolingian Empire. Recent paleogenomic studies^1,18,34,35,51^ have identified a change in genetic background consistent with the proposed timing of these events, with a significant increase of Northern European genetic ancestry in communities from both Pannonia and Northwestern Italy. These results suggest that the historically documented movements of the Langobards – first an expansion to Pannonia from today’s Southern Czechia and Lower Austria then followed by their subsequent migration to Italy – impacted the population demographics of these regions substantially. In modern-day Slovenia, archaeological research has linked certain sites to the Langobard migration based on artefact types known from both Pannonia and Italy – such as brooches, jewellery, belt accessories and weapons, but also every-day objects like combs – generally dated to the middle/second half of the 6^th^ century^42,74,75^. We investigated sites along the presumed route of the Langobard’s migration through the region and complemented it with previously published data (**Table S2**) to test the various interpretations of historical sources, as it is still debated^42,71,73,74,76^ whether they simply passed through the region or established control over it and settled there for an extended period.

Four sites are linked to the Langobards by archaeological research in our dataset. Rifnik (n=46) located to the east, Solkan (n=42) in Vipavska dolina, at the foothills of the Alps along the traditional route crossing the Alps and Cividale San Mauro and Ferrovia, two cemeteries connected to Forum Iuli (Cividale) (n=40), the first city conquered by the Langobards in Italy as described by Paul the Deacon^72^, dated right after its occupation. Both the PCA and the model-based clustering analysis showed significant differences between the sites located on the two sides of the Alps (**Figure 2, Extended Data Fig. 1**). Sites east from the Alps, in modern-day Slovenia exhibit genetic backgrounds similar to that of the LR sites, as explained above, with a high ratio of predominantly Southern European genetic ancestry. The two Cividale sites on the other hand show a substantial increase of Northern European ancestry, similarly to the previously published 6^th^-century sites in Transdanubia (Western Hungary)^35^ and the 6-7^th^ century site of Collegno^18,34^ in Northern Italy. This is corroborated in the F*_ST_* analysis, where elevated values are found between the LR/LA Slovenian communities and the LA Cividale communities (ranging from 0.0021-0.0042) and 19/20 are significant after Bonferroni correction (*Z:* 3.27-9.34; *p:* < 0.001-0.0011), consistent with expectations given the increase in northern ancestry (**Fig. 2; Extended Data Figs. 1, 4; Table S6**). The genetic ancestry observed in individuals from Cividale is also not only different from those of the Slovenian sites, but also from the observed results of Torino-Lavazza and Bardonecchia^51^, two sites in Northern Italy predating the Langobard conquest.

These results suggest that the population movement that led to this significant increase of Northern European ancestry in the Cividale sites as well as Collegno in Northern Italy (presumably the Langobard migration) did not have the same local demographic impact east from the Alps. To investigate the temporal relationship between these sites, we conducted 22 radiocarbon measurements on Rifnik and Solkan (**Table S3**), two sites archaeologically dated to the second half of the 6^th^, beginning of the 7^th^ centuries based on certain jewellery types and belt accessories. The results confirmed the 6^th^ century dating of Rifnik, placing it around the historical dating of the Langobard migration to Italy. Radiocarbon dates, however, place Solkan in the mid-7^th^ to 8^th^ centuries compared to the earlier late 6^th^-early 7^th^ century dating, making it the youngest site in the LA dataset.

Despite the apparent lack of genetic impact, the fact that both Rifnik and Solkan show clear archaeological similarities to Pannonian and Italian Langobard period cemeteries^75,77^—such as the brooches from Rifnik_57 and Rifnik_83, the pottery with stamped-in decoration from Rifnik_86 and the weapon belt from Solkan_18—suggests that the region still belonged to the cultural, and possibly political, sphere of the Langobard influence during their migration, and that territories close to the Italian border of their kingdom continued to be affected even a century later. The founding of the Rifnik community most probably predates the arrival of the Langobards as the settlement/fortification has Roman period roots. At Solkan - despite its later dating - IBD results show strong connections between communities dated from the 5th to the 7th-8th centuries within Vipavska dolina, but far fewer links outside the region, suggesting strong continuity and restricted external genetic influences (**Figure 6; Extended Data Fig. 5; Tables S11 and S12**). Due to its strategically important location overlooking the presumed route of the Langobard migration, the Rifnik community may have played a significant role during their initial efforts to occupy Italy. The Solkan community, however, probably represents the cultural and political influence of the already established Langobard kingdom in Italy in its bordering areas. Nevertheless, this cultural influence does not appear to be associated with any genetic impact, suggesting present-day Slovenia may have been merely a transit zone, with no long-term settlement taking place in the region.

Despite similarities in genetic background across most LA sites, differences in relatedness are observable. Rifnik and Solkan include multigenerational extended pedigrees and the ratio of individuals with 1st to 3rd degree relatedness within the site is proportionally higher compared to the rest of the sites (**Section S1; Table S9**). The structure and size of these pedigrees are similar to what has been observed in the Langobard period Little Hungarian Plain^35^ or Northern Italy^18,34,51^, suggesting that cultural similarities across the multiple regions involved in the Langobard migration might not only be present in the material culture, but also in the social organization and development in these communities.

### Population change detected after the 8th century CE

According to traditional archaeological research, most late antique hilltop sites in present-day Slovenia were abandoned by the end of 7th century CE and the emergence of new sites is not attested until after 750 CE when small rural cemeteries started to appear^41,78^. These newly-established cemeteries—Camberk from Dolenjska, Hrušica near Ljubljana and Ledine, Gojače, Vrtovin_EM, Šturje from Vipavska dolina in our dataset—exhibit little to no spatial continuity with earlier periods and are characterized by markedly different burial practices and material culture (**Section S1**).

Paleogenomic analysis of these sites show patterns of between-site genomic heterogeneity that point to differences in the migration and gene flow across the region during the EM period. The Hrušica (n=14) and Camberk (n=22) sites, dated to the 8th-9th centuries, show a major change in the genetic composition of these communities, with a shift towards predominantly Northeastern Europe on the PCA. The Hrušica and Camberk samples do not share any IBD connections with LA communities except via two samples, Cam-Cl_15 and Cam-Cl_22 (**Table S9**). In addition, only two individuals from Camberk (Cam-Cl_17 and Cam-Cl_22) and none from Hrušica can be modeled by LA populations using qpWave. The remainder of the communities can only be modeled by northern European groups (primarily Anglo-Saxons and Northern Germany) strongly suggesting an influx of new population groups into these regions, (**Figure 3, Table S9**). The distinction of these EM sites (Hrušica and Camberk as well as Šturje) from the older LR/LA sites is further corroborated by the results of our F*_ST_* analysis (**Extended Data Fig. 4**). Site-level F*_ST_* comparisons within the set of LR/LA communities range from 0.000 to 0.0035; however, when these communities are compared to EM communities (e.g., the differentiation between an LR/LA community and one of the EM communities), we found substantially more differentiation, with values ranging 0.0028 to 0.0070, where all comparisons were significant after Bonferroni correction (20/20) (*Z*: 6.35-23.26; *p*: 0.000-2.1×10^-10^); similar results were obtained when we used regional scale comparisons, with all 16 comparisons remaining significant (*Z*: 6.35-23.26; *p*: 0.000-2.1×10^-10^), as well as comparisons between the pooled time periods (LR-EM, F*_ST_* = 0.0035, *Z* = 23.459, *p* = 0; LA-EM, F*_ST_* = 0.0030, *Z* = 19.431, *p* = 0), firmly indicating that these communities are distinct (**Table S6**). The factor analysis also showed a very different Factor 1 profile between the LR and LA sites compared to the EM period, even after controlling for drift due to age of samples, with this primarily driven by Hrušica, Camberk, and Šturje, suggestive of a major change in genomic ancestry between these periods (**Extended Data Fig. 3**). Furthermore, this discontinuity between the earlier sites is even supported by ^87^Sr/^86^Sr values, which occupy much wider ranges when compared with the LR and LA sites, indicating these communities either relocated from a different geographical region or had different agricultural land-use practices (**Figure 4**).

We observe a more complex picture in Vipavska dolina: Ledine, Gojače and Vrtovin_EM show a similar genetic composition to that of the LA sites based on results from PCA and model-based clustering (**Figure 2, Extended Data Fig. 1**). These are small grave groups spatially linked to previous settlements (Solkan, Vrtovin_LA), so both archaeological and genetic evidence indicates small scale population and settlement continuity in the region. Ledine, Gojače, and Vrtovin_EM maintain many IBD connections to LA communities (16.58 connections per possible 1000 pairs) (**Figure 6, Extended Data Fig. 5, Table S12**). With the exception of outlier Ledine_02 (who has substantial northern ancestry unlike the rest of Ledine), the remaining 7 of these individuals can be modeled by at least 3 of the LA communities in the qpWave analysis (**Figure 3**). The later Šturje site, however, showed the same increase in Northeastern European ancestry in the PCA as detected in the other two EM regions—a pattern not observable in LR or LA sites. Notably, despite their very similar genetic backgrounds, Hrušica, Camberk, and Šturje are not necessarily more closely linked together by IBD relationships. When analyzing pairwise IBD connections between these three sites, there are no direct connections between Hrušica and Šturje (**Table S12**). On the other hand, Camberk does maintain connections with Šturje and Hrušica individually (30.81 and 19.84 connections per possible 1000 pairs, respectively). These results indicate that while there are strong IBD connections within sites, these three sites are more loosely connected than in the LA/LR communities, which have connections both within and between sites (**Table S12**).

As the archaeological chronology of EM sites in the southeast Alps is still a matter of debate^79–81^, we radiocarbon-dated 23 graves to establish a more reliable timeline to understand the temporality of the appearance of the Northeastern European ancestry (**Tables S1 and S3**). Based on the refined chronology, Hrušica, Camberk, Ledine, Gojače, and Vrtovin_EM are all dated between ca. 750 and 950 CE and can be considered generally contemporaneous. Šturje, however, is now dated to the 11th century. This indicates that the observed change in genetic composition in the region occurred at different times across the various locales. The arrival of individuals with Northeastern European ancestry likely occurred before or around the 8th century in Emona and Dolenjska, while in Vipavska dolina—the region closest to Italy—population continuity is observable for longer, and communities with Northeastern European genetic ancestry might have appeared only much later, after the 9th century. Since ^87^Sr/^86^Sr values observed at Šturje do not show any difference compared to the values observed at earlier sites in the region, nor compared to the biologically available strontium of the region itself, an alternative explanation could be that this represents the relocation of an already established community within the region that retained its genetic homogeneity, perhaps intentionally through selective reproductive strategies (**Figure 4; Section S2**).

Recent studies have described the same Northeastern European genetic ancestry in other EM populations outside Slovenia, with two recent papers^14,15^ arguing that this ancestry originates from the Baltic and spread through demic diffusion throughout Eastern/Central Europe during the 6th to 8th centuries CE, chronologically coinciding with the expansion of the Slavs attested by the written sources. Other researchers have linked the genetic ancestry itself directly to the Slavs^82^. However, these studies lack samples from the 7th-8th centuries that are crucial to the accurate dating of these processes. When comparing the published communities associated with the Northeastern European ancestry and a similar chronology of 8th-11th c. CE (Velim from Croatia, Gródek from Poland, and Niederwünsch, Obermöllern, Steuden, and Steuden-Melmesdorf from Germany) (**Table S2**) from Gretzinger et al. (2025), we found that these communities cluster closely with Camberk, Hrušica, and Šturje on PCAs (**Extended Data Fig. 7**) and also share 17 IBD links in contrast to only 3 with the LA Slovenian communities (1.82 and 0.16 connections per 1000 pairs, respectively) (**Extended Data Figs. 5-6; Tables S10-12**). We found 11 of these were specifically connecting Camberk and Hrušica with Velim in Croatia, the site closest both chronologically and geographically, yielding 6.49 and 9.47 connections per 1000 pairs, respectively, with connections ranging from 21 to 33cM. In contrast, only 6 connections were detected with the sites from Germany/Poland. Furthermore, we ran qpWave analyses testing if the EM individuals could derive from the Baltic Expansion populations by using the seven penecontemporary source individuals (from Gródek) that were used by Gretzinger et al.^14^. Ultimately we found that for Hrušica, Camberk, and Šturje most individuals could derive their ancestry from these seven Gródek individuals (Hrušica 69%, Camberk 62%, and Šturje 95%) (**Figure 3**).

According to the written sources, Slavic groups arrived in the territory of present-day Slovenia around 600 CE^28,40^, yet this distinct genetic ancestry does not show up at sites dated to that time (Rifnik, Solkan, Zidani gaber, Vrtovin), but rather it only appears from the 8th-9th centuries onwards. It is important to note that archaeological research places the earliest appearance of Slavic settlement in the northeastern part of present-day Slovenia, a region not covered by the present study.^30,35,78^ Furthermore, the three sites with NE ancestry do not represent a homogeneous group. Hrušica and Šturje, while 200 years apart, are both organized around extended pedigrees, i.e. groups of biologically related individuals, that included the majority of the buried individuals (78.6% and 72.7% respectively), while at Camberk only one parent-offspring pair (Camberk_33 and Camberk_36) was observed (22.7%) (**Tables S1 and S9**). The clear differences in chronology, material culture and social structure and lack of IBD connections between these sites indicate that the change in genetic composition in the communities in these three regions was probably not the result of a single event despite the homogeneous Northeastern European ancestry. Some of this genetic change may have even occurred after the region became part of the Ottonian empire in the late 10th c. CE, with people gradually began burying their dead in newly established churchyards^83^.

## Conclusions

The area of modern-day Slovenia served as a crossroads between West and East Europe during the time of the Roman Empire(s), and retained its strategic importance for centuries as the gate to Italy even after the collapse of the western empire. Written sources describe various types of population movements (relocation, emigration, immigration) in this area between the 4th to 11th centuries CE, but their exact nature, social and demographic impact is debated in both historical and archaeological research. Our study reveals a nuanced and regionally diverse population history between the end of Roman rule and the early Middle Ages, where most regions show the appearance of new population groups during EM, but one region also showed evidence of population continuity throughout the investigated centuries.

Despite profound political and social changes, including the decline of Roman urban centers, the rise of hilltop settlements, the documented movement of the Langobards, and the later appearance of Slavic groups in the area, we observed striking genetic continuity across several centuries, with evidence of influx of new groups limited to two sites. The investigated LR and LA communities show clear similarities with a dominant southern European genetic background with the exceptions of late antique lowland sites (Dravlje and Miren), where individuals carrying Central Asian genetic background appear. The overall stable genetic composition of communities dated between the 4th to 7th centuries suggest a level of population continuity between LR centers (Celea, Emona, Castra, etc.) and LA hilltop sites (Rifnik, Solkan, etc.), with an influx of new groups, during the time of the Ostrogothic Kingdom around 500 CE, leaving no lasting genetic impact. Interestingly, the both historically and archaeologically well-evidenced migration of the Langobards from Pannonia to Italy in 568 seems to have left no genetic trace in the area, despite earlier studies showing its significant impact in both Pannonia and Italy. Modern-day Slovenia might have only served as a transit zone for their migration, but changes of material culture and of the role of biological relatedness in the formation of communities attest to their cultural and social impact. These results highlight that cultural transformation and changes in settlement patterns did not necessarily correspond with immediate or large-scale population replacements.

The transition to the EM period however reveals a significant change in genetic ancestry and also differences in the population history of Vipavska dolina, the region bordering Italy and the more eastern regions. From the 8th century onwards, in the Emona and the Dolenjska regions, genetic results show a significant increase of NE European ancestry —a genetic component described as Baltic and linked to the expansion of the Slavic groups in Eastern and Central Europe by recent studies. However, despite the similarity in genetic composition between these newly emerging communities, we observed clear differences in their chronology and social structure. Our results show the diverse genetic and social impact of various types of population movements in the region and emphasise the complexity of these processes rather than a simple and direct correlation between genetic changes and historical narratives.

## Methods

### Genetic data generation and processing

We sampled 370 individuals from Slovenia for the biomolecular analyses. The overwhelming majority of individuals were sampled for DNA from the petrous bone, but a small minority were sampled from teeth and in one case from humerus (LJ-Hrusica_1012-1013-2). Bone samples were cleaned and drilled or powdered at the ELTE Research Centre for the Humanities, Institute of Archaeogenomics. Samples were further processed (DNA extraction and library preparation) in the clean room of the Max Planck Institute for Evolutionary Anthropology (MPI-EVA) in Leipzig, Germany. Libraries. Single-stranded libraries were generated and underwent 1240k SNPs capture and enrichment twice^36–38^. Enriched libraries were sequenced at MPI-EVA using the Illumina HiSeq 4000 platform, sequencing 75-cycle single-end reads, with around 20 million reads per library. Sequence data were bioinformatically processed on site. All fastq files were processed and aligned using BWA^84^ to Hs37d5 (based on GRCh37) in Leipzig, generating BAM files.

Individuals from Cividale (San Mauro and Ferrovia, n=40) were processed separately from the Slovenian dataset using bone samples from the petrous bones alongside the individuals sequenced by Tian, Koncz, Defant et al^18^. Petrous bones for 24 individuals from Cividale Ferrovia and 16 individuals from Cividale San Mauro were processed at the Laboratory of Molecular Anthropology and Paleogenomics of the University of Florence, Italy. The external surface of the petrous bones was brushed with disposable tools and irradiated with ultraviolet light (254 nm) for 30 minutes to eliminate surface contaminants. In order to maximize the recovery of well-preserved endogenous DNA, bone powder was collected from the densest part of the petrous bone (inner ear) as described in Pinhasi et al., using a low-speed micro drill equipped with a disposable disc saw and dental burs. DNA extraction and purification were performed using a silica-based protocol (Dabney et al. 2013). Double-stranded libraries with partial uracil-DNA glycosylase (UDG) treatment were prepared (Rohland et al). To monitor for potential reagent contamination, blank controls were included during DNA extraction and library preparation. The Cividale libraries underwent the same 1240k SNPs capture as the Slovenian DNA libraries at the Max Planck Institute for the Science of Human History (MPI-SHH) in Jena, Germany. Sequences from Cividale were processed analogously to and alongside the data published by Tian, Koncz, Defant et al^18^.

VCF files were called from the BAM files using an in-house indent caller (https://github.com/kveeramah). The indent caller disregarded the first and last five bases of any given read and produced diploid VCFs that included diploid genotypes and genotype likelihoods.

We calculated autosomal 1240k SNPs sequencing coverage using gatk v4.2 DepthOfCoverage^85^. For most downstream analyses, individuals with less than 0.10× coverage were excluded. We used the Sex.DetERRmine pipeline^86^ (https://github.com/TCLamnidis/Sex.DetERRmine), which used relative X and Y chromosome coverage, to test for genetic sex. Schmutzi was used to calculate mitochondrial contamination in all individuals, while Angsd was used to calculate nuclear contamination in genetic males.

In order to assess mitochondrial genome haplogroups, endogenous mitochondrial assemblies from Schmutzi^87^ were analyzed using MitoTool 1.1.2^88^. We also identified the Y chromosome haplogroups for genetic males from all sites based on the 1240k Y chromosome SNPs. Phylogenetic analysis was performed using the published ISOGG database, allowing for identification of NRY haplogroups.

Information on all newly sequenced individuals are presented in Table S1.

### Principal Component Analysis

PCAs were generated using the POPRES database as a modern reference background consistent with Vyas, Koncz, et al.^51^ A PCA was conducted for each individual using smartPCA^46,47^ and a Procrustes transformation^34,89^ was used to merge all individuals into a single PCA. Analyses conducted using 328,670 SNPs from the imputed POPRES dataset from Veeramah et al.^50^ Prior to principal component analysis, all diploid genotypes were converted into pseudohaploid using an in-house script. Heterozygous genotype calls were made hemizygous for the allele with greater allele depth; in cases of ties, one was chosen randomly. PCAs for each individual in the Procrustes transformation were conducted using an automated script (modified from https://github.com/ShyamieG/ to conduct individual PCAs in parallel). Individuals with less than 10,000 SNPs overlapping with the POPRES dataset were excluded. PCAs were plotted using matplotlib^90^ and kernel density estimation plots were generated using the seaborn package’s kdeplot(). PCAs were also generated using the AADR populations as the modern reference set^53–62^, using a subset of pan-Eurasian populations genotyped on the Affymetrix Human Origins array. Pseudohaploidized genotypes were processed analogously to the POPRES analyses and used an overlap of 536,325 SNPs.

### Model-based clustering analysis

Model-based clustering analyses were conducted using fastNGSadmix^45^, using both a modern 1000 Genomes Project^52^ reference panel and a panel of penecontemporaneous individuals. Both panels are based on Vyas, Koncz, et al.^51^ However, the penecontemporaneous panel also includes the Central Asian (CASIA) reference panel, as used in Tian, Koncz, et al. Beagle PL files were generated for all individuals with coverage ≥0.1× using vcftools.^91^ For each individual, each fastNGSadmix analysis was conducted 50 times and a custom python script was used to select the run with greatest (least negative) likelihood.

### Model testing using qpWave and qpAdm

In order to test whether individuals were consistent with deriving from populations from previous time periods or different regions of the world, we conducted qpWave analyses using Admixtools^53^. Due to the sensitivity of qpWave analyses to the type of pseudohaploidization used in the our PCAs (described above), we use an in-house random read caller on all analyzed individuals as well as comparative and reference populations. Following the merger of all the random-read VCFs, we retained 703,069 autosomal 1240k SNPs. For our qpWave analyses, we used Morocco_IM, Iran_Neolithic, WHG, Steppe_Eneolithic, and Anatolia_Neolithic as the right populations, and a rotating set of left populations. Each comparative left population for each time period was constructed of ten randomly selected individuals (Table S2)^36,37,54,55,92–99^. All left populations (comparative or Slovenian) were constructed such that there were no biological relatives within the same panel.

For left populations we divided Europe into geographic regions and then divided them based on time period (only including individuals from the same time period as well as the immediately previous period), such that the LR panels only contained individuals from the LR period, the LA panels contained individuals from the LA/LR periods, and the EM panels contained individuals from the LA/EM periods. Due to limited samples, the Asian panels remained static across all time periods. Each individual was tested one by one for each corresponding left population. Custom python scripts were used to calculate the number of individuals per Slovenian community consistent with deriving from each respective population (p > 0.05). Further details are presented in Section S3.

### Calculating F*_ST_*

We calculated F*_ST_* using smartPCA^46,47^ We used the exact same random read pseudohaploid dataset to calculate F*_ST_* between the communities in our dataset that was used for qpWave and qpAdm. As the dataset is pseudohaploid, we enabled the inbreeding setting in the analysis. Not all individuals were used in this analysis, as we had to account for biological relationships, and thus we selectively omitted based on pedigrees, to avoid having relatives within any group. We calculated values on three scales: the community level (e.g., Solkan vs. Rifnik), the regional level per time period (e.g., Celeia LR vs. Emona LA), and then by whole time period (e.g., LA vs EM). We included the sites from Cividale in these analyses, but we bifurcated the LA time period to reflect them separately. We did this, because prior analyses suggested that many individuals from Cividale have northern ancestry not found in the other LA communities (**Fig. 2; Extended Data Fig. 1**). All analyzed groups required 10 usable individuals to be considered for calculation (e.g., Hrušica was not used as a group due low sample size (n=4) after accounting for biological relatives). Furthermore, we did not use the individuals from Gojače, Ledine, or Vrtovin in any of these analyses due to small sample sizes. To account for multiple testing corrections, we used a Bonferroni correction (α = 0.05) for each scale of analysis (i.e., site level, regional level, time period level).

### Factor Analysis

Factor analysis was performed on the genome-wide imputed genotypes from Slovenia using the tfa package in R4.5.1^63,64^. First, we identified a set of putatively unrelated individuals, as assessed by having no pairwise pi_hat value from lcMLkin > 0.05 from the full set of n=410 samples (**Table S9**). We used a greedy algorithm that iteratively removed the sample with the most connections to other samples, resulting in a final set of 251 individuals. We then used Plink v1.9^100,101^ to filter out SNPs with a minor allele frequency <0.1 (--maf 0.1) and hardy-weinberg deviation p-value <1 × 10^-8^ (--hwe 1e-8) and linkage disequilibrium (--indep-pairwise 50 5 0.2), resulting in a final SNP amount of n=78,236. Samples were assigned ages in years before present based on the median value estimated from the C14 dating (**Tables S1 and S3**). Samples from sites with no C14 estimates were excluded from the analysis, resulting in a final set of 217 individuals (LR=117, LA=66, EM=34).

PCA on the genotype matrix was performed using prcomp, with the variance explained being notably increased for PCs 1, 2 and to some extent 3 (Supplemental Fig S27a), suggesting the use of 2-3 ancestral populations in the subsequent factor analysis. The coverage_adjust function of tfa was applied to the genotype matrix to perform a correction for coverage across samples using a latent factor regression model with four factors (*k*=4). A drift parameter value, *λ*, of 10 was chosen based on running a grid search from log_10_(-3 to 3) with 10 gridpoints using the choose_lambda function for *k*=2 factors (Supplemental Fig S27b). Factor analysis was then performed for *k*=2 and *λ*=10 using the tfa function. We also ran this procedure for *k*=3 and obtained equivalent results for factors 1 and 2 as when using *k*=2, while factor 3 showed no noticeable structure. When plotting Factor 1 against age we fit a loess-smoothed trend line with 95% confidence intervals to each period using geom_smooth (method-loess, se=True).

### Biological relatedness analysis and Identity-by-descent

We tested for relatedness within our dataset using the program lcMLkin^102^. For this analysis, we included all individuals regardless of coverage level, but we only considered the lcMLkin results for pairs of individuals sharing coverage at 10,000 SNPs or more. Analyses were conducted consistent with Vyas, Koncz, et al.^51^. We used genotype likelihood data from 1,079,996 autosomal 1240K SNP sites as input and three reference panels based on the 1000 Genomes CEU and TSI populations (CEU, TSI, and CEU+TSI). We used the kinship coefficients from lcMLkin alongside mtDNA and NRY haplogroups and osteological information to inform pedigree constructions.

In order to assess more distant relationships between sites and regions, we also conducted identity by descent (IBD) analyses using ancIBD^69^. Prior to using ancIBD, all individuals were imputed using GLIMPSE v1.1^103^. We imputed all the sites within the 1000 Genomes Project Phase 3 v5a dataset^52^ using genotype likelihoods from whole genome VCFs. Following imputation, we filtered down the dataset back to the autosomal 1240k SNPs. We conducted ancIBD using a standard pipeline, using default parameters. For this analysis, all individuals with coverage less than 1× were excluded, resulting in a dataset where all individuals had around ∼600,000 SNPs or higher natively covered pre-imputation. Additionally, individuals where less than 65% of sites have a max genotype probability rating of 0.99. In order to be conservative in our IBD analysis, we restricted our focus to segments 20cM or larger.

We normalized the rate of observed IBD connection (edges) based on the number of possible edges (which are the number of possible pairs between the two involved populations). We used this statistic to calculate the rate observed. per 1000 possible pairs, which we used to estimate the strength of IBD connections.

For ancIBD analyses, we focused on three separate scales. Initially, we focused on connections between the individuals in our dataset (Figure 6). This was to understand connections at a local level to see how the five regions we studied were connected to each other through time and space. For the next scale, we included comparative populations from the Piedmont of Italy^18,34,51^, the Lake Balaton region^34,51^, the Little Hungarian Plain^35^, as well as four communities associated with the Baltic Expansion^14^ (**Extended Data Fig. 5**). For the final scale, we include all the communities that we included in the comparative populations used in our dataset.

## Data Availability

Our newly generated sequence data from 410individuals are available from the NCBI Sequence Read Archive (SRA) database under accession PRJNA1378448. We accessed the POPRES (Population Reference Sample) dataset collected and published by Nelson et al.^49^ from dbGaP (accession dbGaP: phs000145.v4.p2).

## Acknowledgement

This project was funded by the European Research Council (ERC) under the European Union’s Horizon 2020 Research and Innovation Programme (HistoGenes, grant 856453 ERC-2019-SyG). We would like to thank the laboratory staff of the ELTE Research Centre for the Humanities, Institute of Archaeogenomics for their work in the project: Viktória Oravecz, Viktória Bódis, Koppány Kerestély, Botond Heltai and Melinda Megyes.

## Extended Data Figures

**Extended Data Figure 1.**
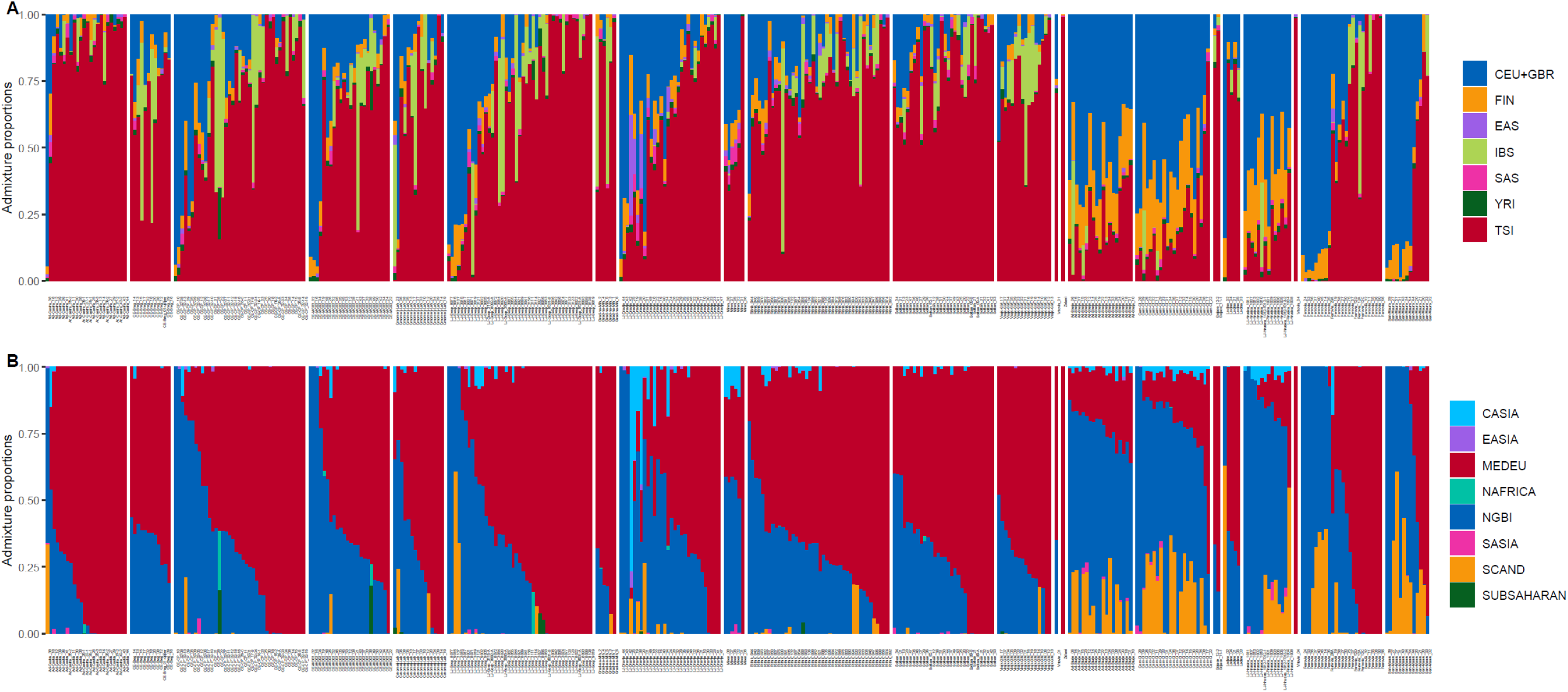
Model-based ancestry proportions for novel individuals from Slovenia and Cividale. In a. proportions were estimated using fastNGSadmix while using 1000 Genomes populations as references (CEU+GBR: “Northern Europeans from Utah” (CEU) and “British in England and Scotland”; FIN: “Finnish in Finland”; IBS: “Iberian populations in Spain”; TSI: “Tuscans from Italy”; EAS: the East Asian super-population; SAS: the South Asian super-population; and YRI: “Yoruba in Ibadan, Nigeria”). In b. proportions were estimated using penecontemporaneous individuals to form reference populations (CASIA: Central Asia, EASIA: East Asian, MEDEU: Mediterranean Europe, NAFRICA: North African, NGBI: Northern Germany/British, SASIA: South Asian, SCAND: Scandinavia, and SUBSAHARAN: sub-Saharan Africa). Individuals are sorted by site, with the LR sites first and EM sites last, followed by the sites from Cividale.

**Extended Data Figure 2.**
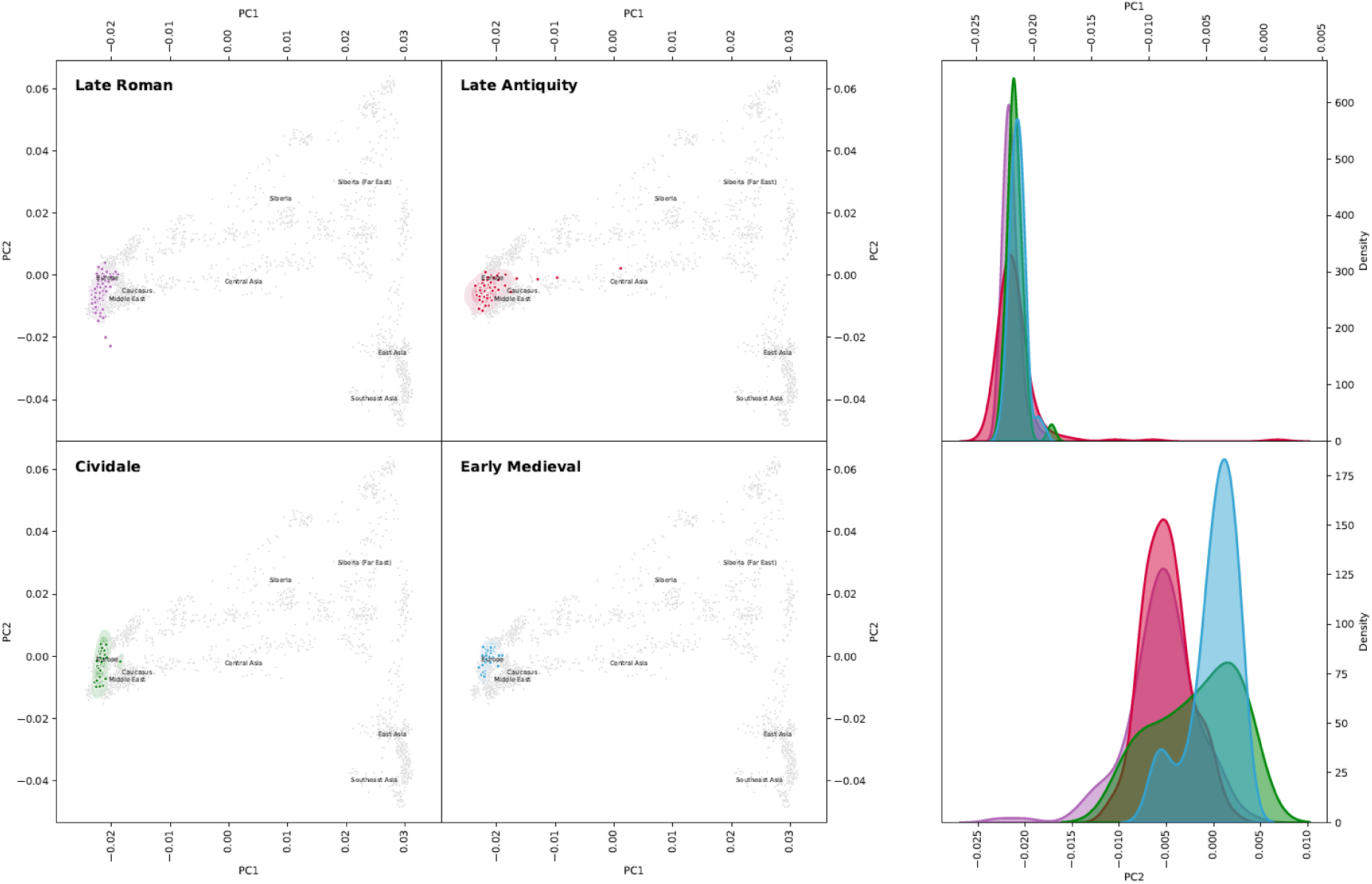
PCA of the LR, LA, and EM populations from modern Slovenia, alongside the 6th-7th century sites of Ferrovia and San Mauro from Cividale, Italy. Pan-Eurasian individuals from the AADR dataset are used as the reference panel.^53–62^. Additionally, Kernel Density Estimation plots of PC1 and PC2 are depicted for the LR, LA, and EM plots.

**Extended Data Figure 3.**
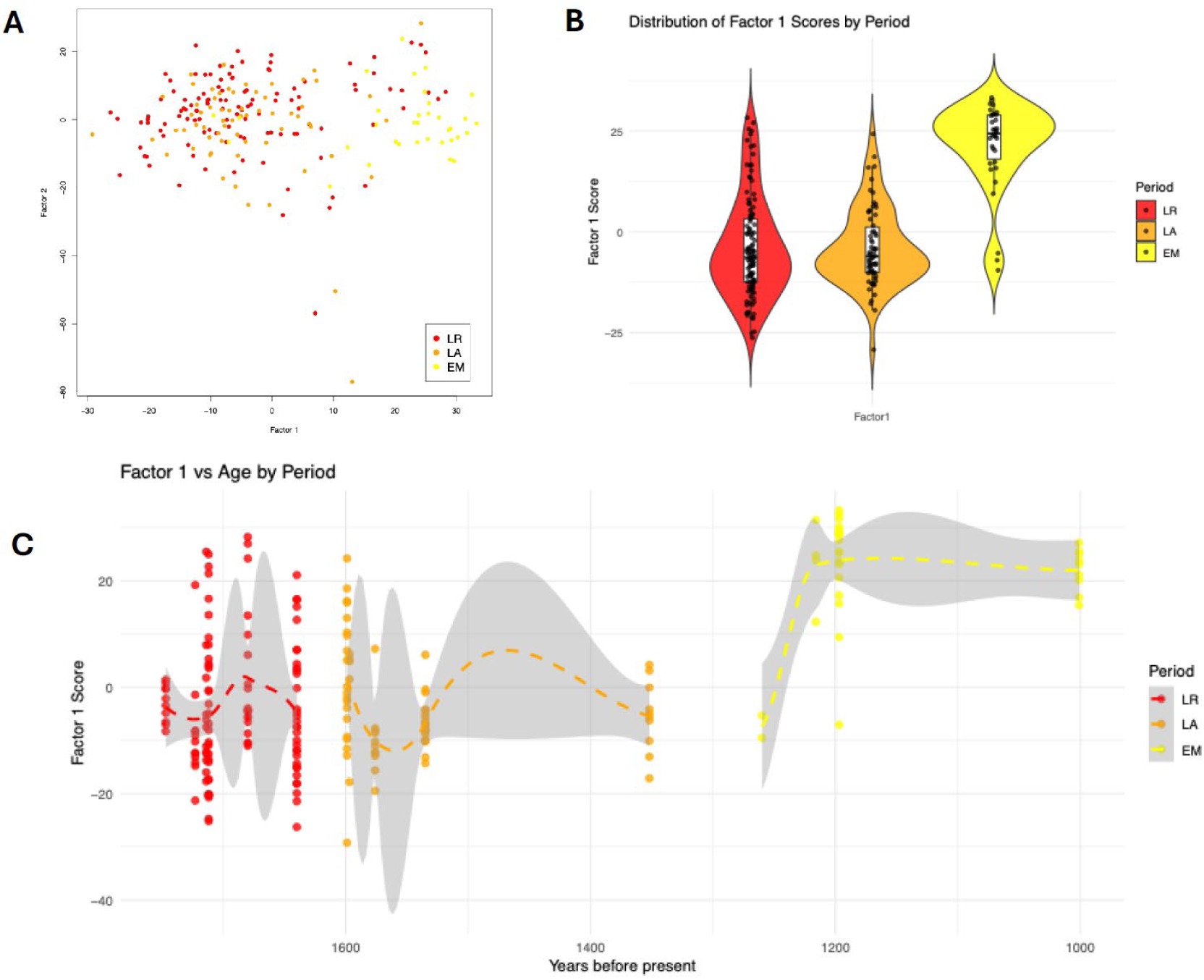
**A.** Scatter plot of Factor 1 and Factor 2 scores for a set of imputed genotypes labeled by period. Factor 1 and factor 2 explain 56% and 44% of the variance captured by the temporal factor model respectively. **B.** Violin plot showing distribution of Factor 1 scores, grouped by period. **C.** Plot showing Factor 1 scores for each sample with respect to their age (i.e. time before present). Dashed lines represents the loess-smooth trendline for each period, and grey area represents their respective 95% confidence interval.

**Extended Data Figure 4.**
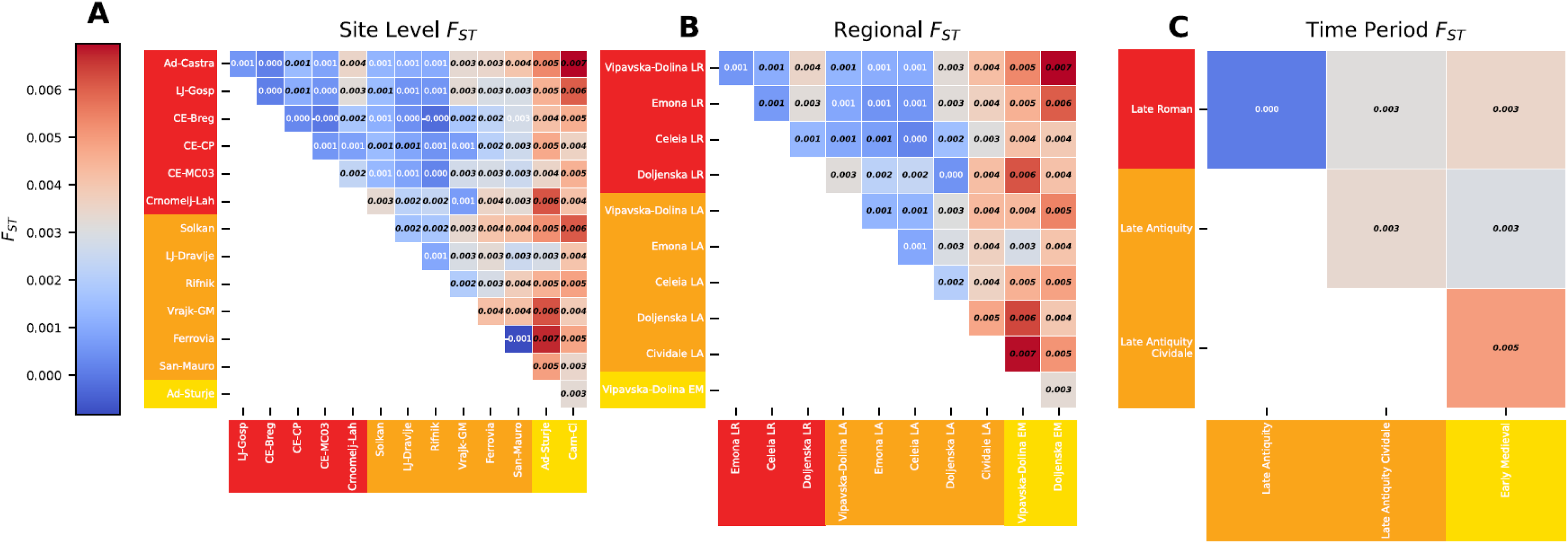
Pairwise F*ST* values. **A** presents values for comparisons between each community (with 10 or more available individuals), **B** presents values for comparisons between each region per each time period (with 10 or more available individuals), and **C** presents values for each of the time periods in whole. Redder cells indicate higher values and greater differentiation. Values that are significantly different after Bonferroni correction (α =0.05) are bold, italicized and in black font. Slovenia communities are color-coded on both x- and y-axes based on time period based on **Figure 1**.

**Extended Data Figure 5.**
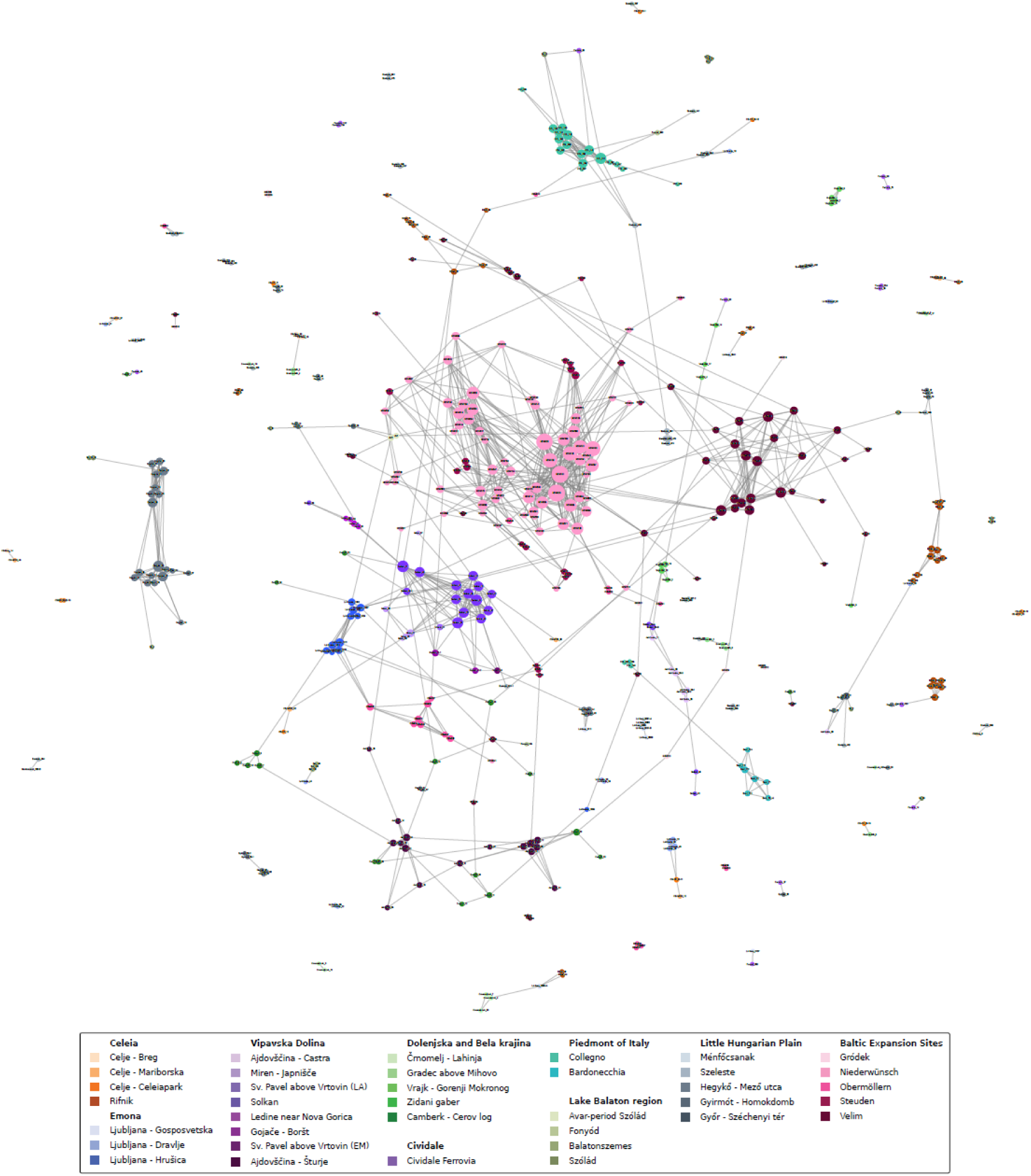
This figure presents the full IBD network generated using ancIBD^69^. Analysis include the novel data from Slovenia and Cividale generated in this study as comparative populations from the Piedmont of Italy^18,34,51^, the Lake Balaton region^34,51^, the Little Hungarian Plain^35^, as well as four communities associated with the Baltic Expansion^14^. Individuals are labeled and colored by site. The size of the node reflects the number of connections, and the width of a connection represents its strength. This network only reflects shared tracts greater than 20cM in length. Individuals with no connections have been omitted from the figure.

**Extended Data Figure 6.**
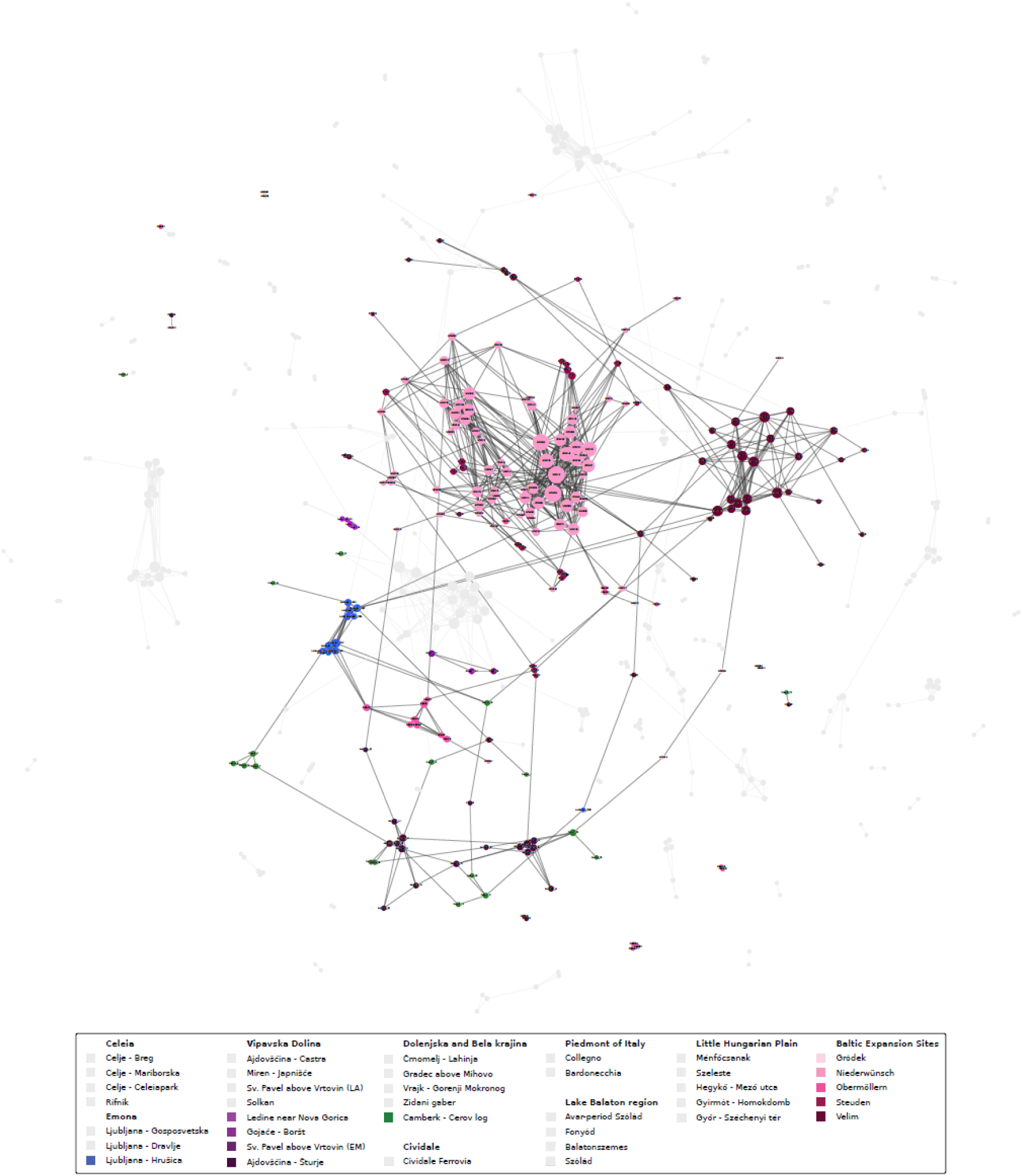
This figure presents the full IBD network generated using ancIBD^69^; however, this figure focuses on the Early Medieval individuals in the analysis. Individuals from previous time periods have been made grey and are unlabeled.The size of the node reflects the number of connections, and the width of a connection represents its strength. This network only reflects shared tracts greater than 20cM in length. Individuals with no connections have been omitted from the figure.

**Extended Data Figure 7.**
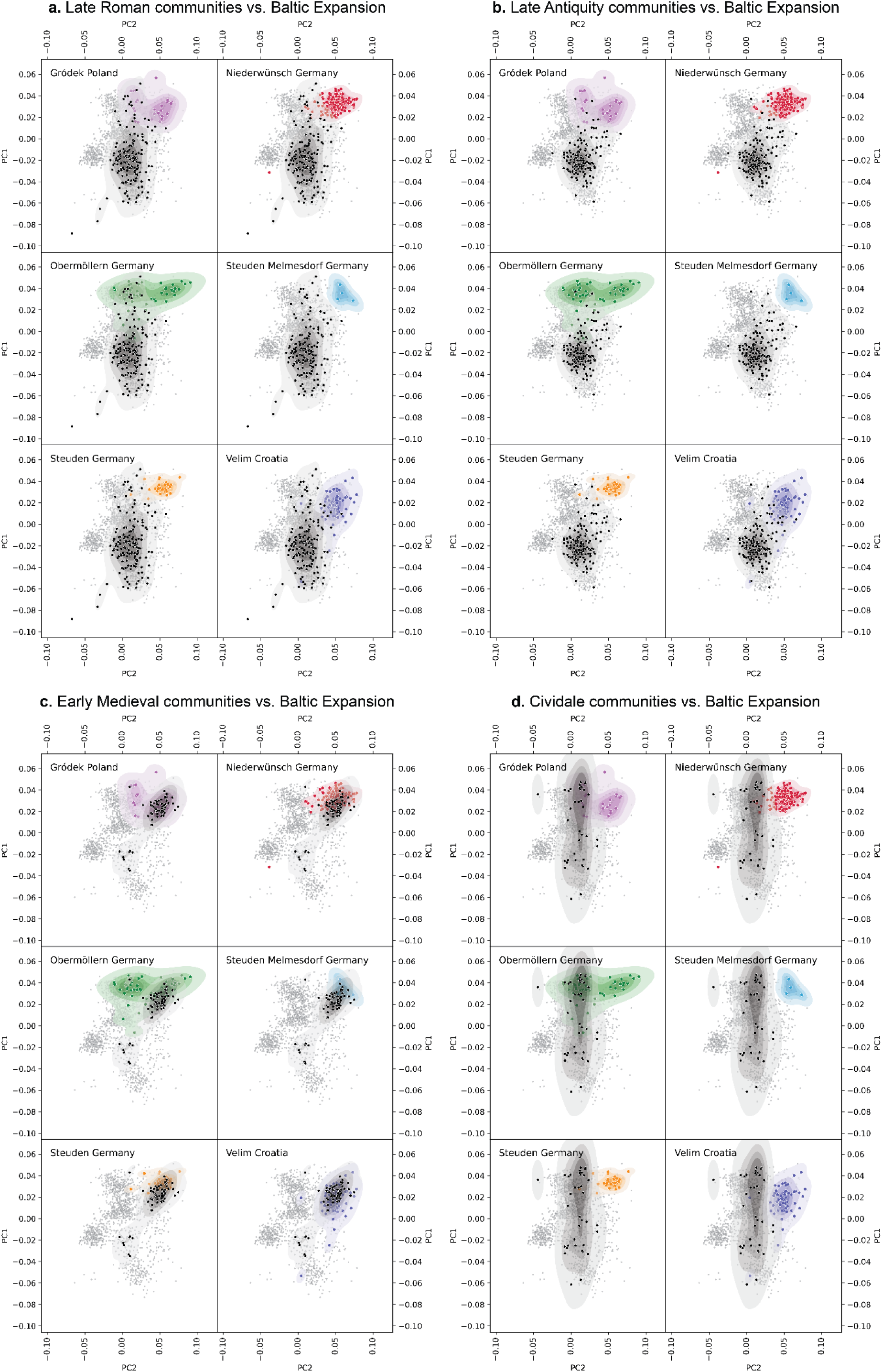
PCA of the LR, LA, and EM populations from modern Slovenia and the Cividale communities (in black) compared to the 8th-11th c. CE of Gródek from Poland, Niederwünsch, Obermöllern, Steuden, and Steuden-Melmesdorf from Germany, and Velim from Croatia^14^. Individuals from the POPRES^49,50^ are used as the reference panel.

## Author contribution

Conceived and led by: T.M., I.K., K.R.V. Formal analyses: D.N.V., I.K., T.M., T.L., N.F., Y.T., P.F. Sample preparation and laboratory work: I.K., T.L., N.F., M.B., B.G.M., P.F., U.B., H.B., A.B., B.B., R.F., A.G., C.G., Z.H., Š.K., A.K., P.M., A.M., M.P., Ri.R., Ro.R., P.S., L.T., K.U., S.V., B.Ž. Visualization: D.N.V., I.K., T.L. Writing: D.N.V., I.K., T.M., T.L., N.F. Supervision: D.C., A.Sz-N., Z.H., W.P., J.K., T.V., P.J.G., K.R.V.

