## Supplemental Texts for "The shifting dynamics of ancestry and culture at a post-Roman crossroads"

**Supplemental Sections**
**S1 Description of Archaeological sites**

**Section 1. Celeia region case study**

Municipium Claudium Celeia (Celje) was a Roman town in the province of Noricum Mediterraneum, situated on the itinerary road Poetovio-Aquileia between Pannonia and Italy. It flourished between the 1st-4th century CE and was gradually abandoned in the second half of the 5th c. CE. Mariborska cesta III (183 graves) and Celeiapark (36 graves) sites are parts of the northern and eastern necropoleis of Celeia respectively. The sites are not far from the early Christian church and baptistery, Mariborska cesta III is dated to the 4th c. and Celeiapark to the late 4th-5th c. CE. Breg (20 graves) is part of the southern necropolis of Celeia, located on the southern (right) bank of the Savinja river, dated to the second half of the 4th c. CE. (Table S1 and S3)

After the town was abandoned the wealthier inhabitants moved either to safer regions in the Adriatic coast or Italy. A part of the late antique population built settlements on hilltops in the hinterground of the Roman towns. Rifnik near Šentjur is one such hilltop site, consisting of a defence wall, several masonry dwelling houses, water cistern and two churches, and a cemetery with 109 graves outside the defence wall. It is a potential location for the translocated people from Celeia, positioned only about 10 km away.

**Sites: Mariborska cesta III, Celeiapark, Breg**

In the northern and eastern necropoleis of Celeia (archaeological sites Mariborska cesta III and Celeiapark) in 88 graves 100 individuals were buried. Most of the graves were singular, while two to four individuals were buried together in the rest of the graves. Additionally, at Celeiapark a mass grave with 21 skulls was discovered. Thus, all together, 121 individuals were documented. One third of the individuals were non-adults, while two thirds were adults with 60:40 ratio males vs. females. In the mass graves, one subadult male and four females were present, twelve were adult males and five undetermined. Pathological changes observed indicate various dental and joint diseases, possible lack of nutrients and rarely healed fractures, neoplasms or inflammations. Out of the 60 discovered individuals buried in 57 graves, 24 individuals (40%) were sampled from Mariborska cesta III. All the individuals sampled were adults, 10 were anthropologically females (42%) and 14 were males (58%). Out of 40 individuals buried in 31 graves at Celeiapark, 22 were sampled. One non-adult individual of undetermined sex and 21 adults, 10 females (48%), 8 males (38%) and three individuals (14%) of undetermined sex were sampled.

At Breg, 13 graves (15 skeletons) had been discovered in 1955 and 2 more in 2020, but the bones are not preserved. 20 graves were excavated in 2010 and out of those 21 individuals could be sampled from this site. All the graves were singular, with the exception of graves 1 (two adult females) and 6 (40 weeks in utero child (not sampled) next to an adult female). 15 individuals (71%) were adults and five were non-adults (24%), predominantly children younger than 10 years of age, and one juvenile individual (5%). Anthropologically, eight (53.3%) adult individuals were assessed as female and five (33.3%) as males, two (13.3%) were undetermined.

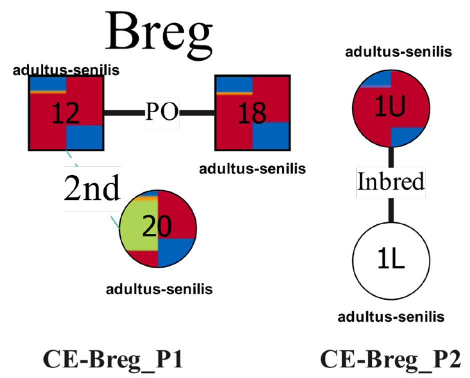


**Figure S1: Pedigrees found at Breg. Squares indicate males, while circles indicate females. Individuals are colored based on their fastNGSadmix results, with the modern panel results on the left and the ancient panel results on the right (Extended Data Fig. 1). The connection PO represents a parent offspring pair where the polarity is unknown. Unknown relationships are represented using dashed lines labeled with the degree.**

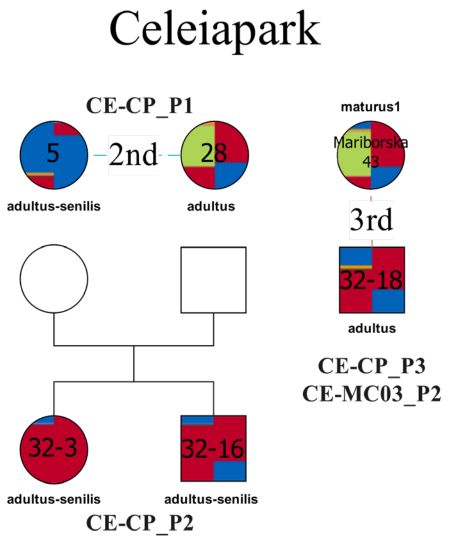


**Figure S2: Pedigrees found at Celeiapark. Squares indicate males, while circles indicate females. Individuals are colored based on their fastNGSadmix results, with the modern panel results on the left and the ancient panel results on the right (Extended Data Fig. 1). The connection PO represents a parent offspring pair where the polarity is unknown. Unknown relationships are represented using dashed lines labeled with the degree.**

### Mariborska

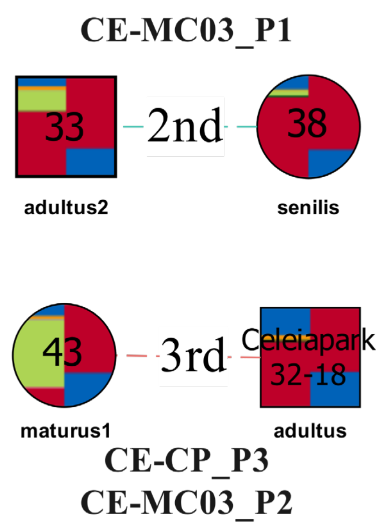


**Figure S3: Pedigrees found at Mariborska. Squares indicate males, while circles indicate females. Individuals are colored based on their fastNGSadmix results, with the modern panel results on the left and the ancient panel results on the right (Extended Data Fig. 1). The connection PO represents a parent offspring pair where the polarity is unknown. Unknown relationships are represented using dashed lines labeled with the degree.**

Relatedness:

Mariborska cesta: **P1**: adult male Mariborska_33 is 2nd degree related to adult female Mariborska_38, they are buried next to each other.

Celeiapark: **P1**: adult female Celeiapark_5 is 2nd degree related to adult female Celeiapark_28.

Mass grave: **P2**: adult female Celeiapark_32-3 and adult male Celeiapark_32-16 are siblings.

**P3**: adult male from the mass grave Celeiapark_32-18 is 3rd degree related to adult female Mariborska_43.

Breg: **P1**: adult males Breg_12 and Breg_18 are a P-O pair. Breg_12 is 2nd degree related to adult female Breg_20.

**P2**: adult females Breg_1U and Breg_1L, buried together, show signs of inbreeding.

**Site: Rifnik**

Rifnik above Šentjur is a fortified hilltop settlement at 568 m.a.s.l. ca. 15 km to the southeast from Celeia. It is partially protected by rock face and partially by a defence wall, built already in the late Roman period. Some of the buildings were already in use in the 4th c. CE as well, while in the 6th c. the defence wall was strengthened with towers, two churches, a water cistern and most of the houses were constructed. About 60 m below the settlement on the southern slope a cemetery belonging to the settlement dates to the 6th c. CE. Furnished graves and some finds from the settlement led the researchers to interpret the site as a civil and military fortification with the presence of Ostrogothic and Langobard state representatives.

109 graves are known from the cemetery while a sarcophagus with 2 skeletons was found in the larger church. Radiocarbon dating puts the majority of the buried individuals to the second half of the 6th c.

From 109 documented graves at Rifnik including a double burial; altogether the graves contained 112 individuals with an unbalanced male-to-female ratio: 42 adult females, 23 males, 25 indifferent adults and 22 subadults. Neonati were completely missing, while other subadults were underrepresented at the burial site. Osteological analysis was heavily limited by the poor preservation of the material; in many cases only differences between adults and children were identified without more accurate age estimation possible.

Altogether 46 samples were collected from 112 individuals buried at the site. Sample collection was heavily limited by poor preservation of the osteological material. As a result samples of 43 or 44 individuals (ca. 38%) were genetically analysed. The sampled individuals include 24 genetic females (19 adults and 5 subadults) and 22 genetic males (13 adults, 9 subadults) showing a much more balanced male-to-female ratio compared to the osteological sex estimation.

Out of 44 analysed individuals (two samples were identified as double sampling and were excluded) 26 (59.1%) showed biological relatedness with other members of the population, this group includes 11 females (9 adults and 2 subadults) and 13 males (9 adults and 4 subadults). Compared to the balanced sex ratio of related adults, among unrelated individuals there are twice as many adult females as males (4 compared to 8).

| | |
|---|---|
| Related | 11 females (3.3): 9 adults (3.6) and 2 subadults (1.5) |
| | 13 males (2.1): 9 adults (2.6) and 4 subadults (1.2) |
| Unrelated | 8 males (2.0): 4 adults (1.5) and 4 subadults (2.3) |
| | 10 females (1.7): 8 adults (2.1) and 2 subadults (0.0) |

Adults were buried with more artefacts than subadults (2 to 1 on average), females were buried with more artefact types on average than males (3.1 to 1.8) and individuals with relatedness were buried with more artefact types in general than unrelated individuals. The latter is most notable in case of adult females (3.6 artefact types on average compared to 2.1; the average among all adults at the site is 1.9), but also observable in case of adult males (2.6 to 1.5) and subadult females (1.5 to 0). The sole exception are subadult males with 1.2 to 2.3 in favour of unrelated individuals.

Genetic analysis identified seven pedigrees at the Rifnik site with the largest containing 8 members (initially 9 with a possible twin that is the result of double sampling one individual) in four generations and multiple smaller groups with 2 to 5 related individuals. One pedigree (Pedigree 4) also showed cross site connections to Dravlje.

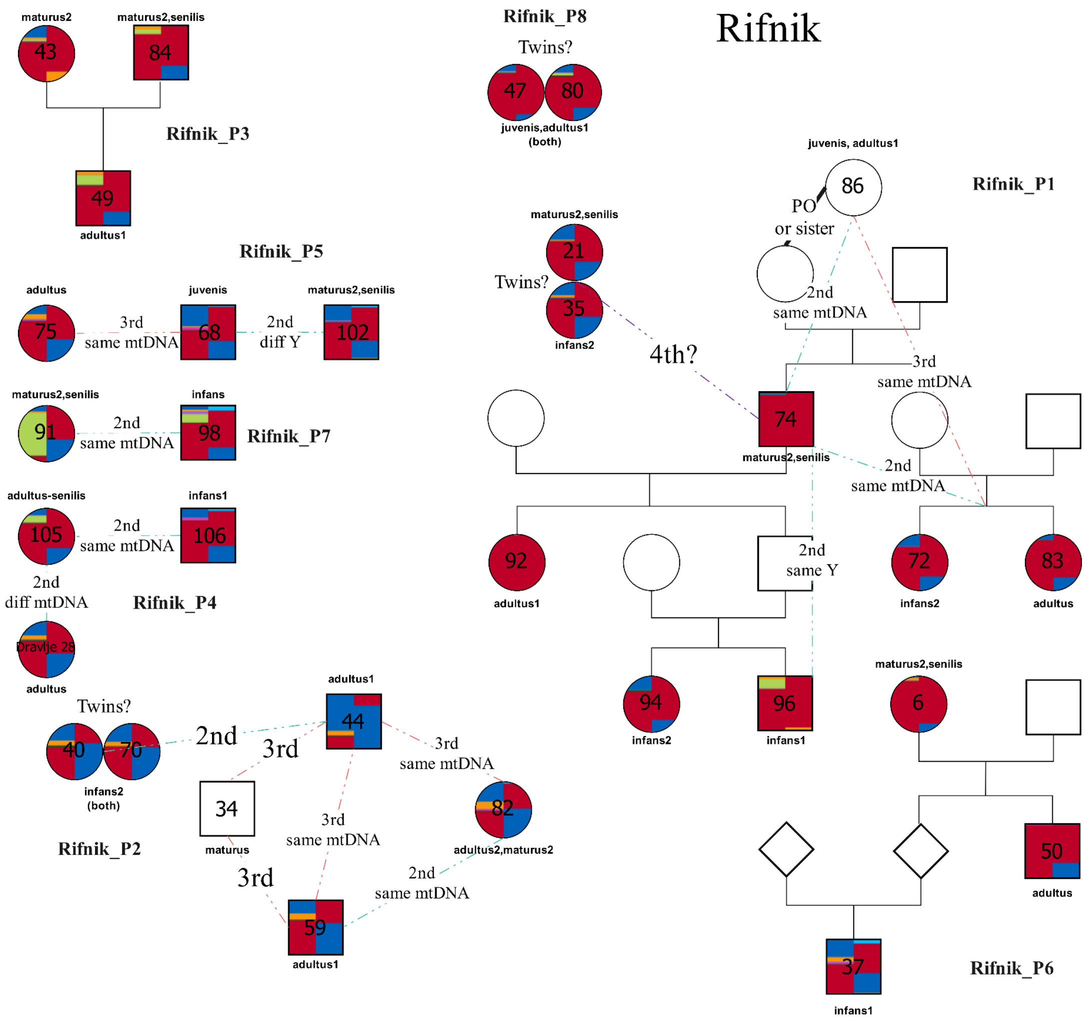


**Figure S4: Pedigrees found at Rifnik. Squares indicate males, while circles indicate females. Individuals are colored based on their fastNGSadmix results, with the modern panel results on the left and the ancient panel results on the right (Extended Data Fig. 1). The connection PO represents a parent offspring pair where the polarity is unknown. Unknown relationships are represented using dashed lines labeled with the degree.**

**Pedigree 1 (P1)**: The largest pedigree at the Rifnik site with 8 members. Rifnik_021 and Rifnik_035 were identified as twins, but a more likely explanation is the double sampling of one individual (possibly Rifnik_021). P1 spans at least 3 (possibly 4) generations at the site. Its identified members are sets of siblings (Rifnik_094 and Rifnik_096; Rifnik_072 and

Rifnik_083) and one pair of parent-offspring (Rifnik_074 and Rifnik_092; Rifnik_074 is also the grandparent of Rifnik_094 and Rifnik_096) with additional more distantly related (3rd-4th degree) individuals (Rifnik_021 and Rifnik_086). Accurate reconstruction of the pedigree was problematic due to the large number of missing individuals, probably as the result of the limited possibilities of the sampling. The pedigree included 6 females (4 adults and 2 subadults) and 2 males (1 adult and 1 subadult).

Multiple burials from the pedigree can be dated to the middle of the 6th century, such as Rifnik_083 based on the Schwechat-Pallersdorf type S-brooch and Rifnik_086 based on its pottery with stamped-in decoration. The direct parallels of these types can be found mainly in 6th century Pannonia, but in smaller numbers they are also present in late 6th-century Italy, where they are elements of a newly appearing material culture that is interpreted as the result of the Langobard migration.

Members of P1 were buried with more artefact types (4.1) on average than the rest of the community. The pedigree includes the three individuals (all adult females) with the highest number of artefact types: Rifnik_021 (8), Rifnik_083 (9) and Rifnik_086 (9). Their grave assemblages included brooches and other jewellery items, various types of combs, belt elements and tools of everyday life.

Members: Rifnik_021; Rifnik_035 (excluded); Rifnik_072; Rifnik_074; Rifnik_083; Rifnik_086, Rifnik_092; Rifnik_094; Rifnik_096

**Pedigree 2 (P2)**:

With 5 (originally 6 members, but the twins were later invalidated[1]) members (4 males and 2 or 3 females) P2 is the second largest pedigree at Rifnik. The accurate structure of the pedigree was impossible to reconstruct as it lacks 1st degree relatedness. Its members are concentrated around the middle part of the site, where most burials were unavailable for sampling due to poor osteological preservation, so it is possible that originally the pedigree might have been much larger.

The burials of P2 generally contained a low number of artefacts (1.6 on average) and none of the types are well datable and the coin of Constantine I (306-337 CE) is only useful as a terminus post quem.

members: Rifnik_34, Rifnik_44, Rifnik_59, Rifnik_82, Rifnik_40 (excluded), Rifnik_70 (excluded)

**Pedigree 3 (P3)**:

Trio of mother-father-offspring buried around the middle area of the site, but not in close proximity. The burial of the already adult male offspring contained datable artefact in the form of a shield-on-tongue belt buckle (Schilddornschnalle) decorated with cross motif generally

[1] Grave 40 was described in the original publication as a burial of an adult male (Bolta 1981, 32), so the twins are probably the result of double sampling (one pp ls and one pp ld) of a single genetically male individual, possibly an adult based on the pars petrosae, so they probably belong to the individual from grave 40.

dated to the middle of the 6th century. A 4th-century coin of Valentinian I (364-375 CE) is only useful as a terminus *post quem*.

members: Rifnik_43, Rifnik_84, Rifnik_49

**Pedigree 4 (P4)**:

2nd-degree related adult female and subadult male buried directly next to each other in the southern part of the cemetery. Stratigraphic evidence based on the superposition between Rifnik_086 and Rifnik_106 suggests that P3 predates P1. The earlier dating is also supported by the 2nd degree relatedness between Rifnik_105 and Dravlje_028 from the predating Dravlje site.

members: Rifnik_105, Rifnik_1106, Dravlje_28

**Pedigree 5 (P5)**:

Adult female Rifnik_75 is 3rd degree related to subadult male Rifnik_68 who is 2nd degree related to adult male Breg_102.

**Pedigree 6 (P6)**:

Adult female Rifnik_6 is the mother of adult male Rifnik_50.

**Pedigree 7 (P7)**:

Adult female Rifnik_91 is 2nd degree related to subadult male Rifnik_98.

There is no clear correlation between the spatial organization of the site and the pedigrees. The spatial analysis is limited by the fragmentary sampling of the site, so any observable patterns might be misleading. Members of the smaller pedigrees (3 or less members; P3, P4, P6, P7) with the exception of P5 are buried close, but never directly next to each other. Members of P2 are concentrated just north of the central area of the site, while P1 has multiple members in both the northern and southern parts.

#### Section 2. Emona region case study

Colonia Iulia Emona (Ljubljana) was a Roman town in the X. regio of Italy, strategically situated at the crossroads of Roman communications from Pannonia to Italy and to the Alps. It flourished in the 4th c. CE when an early Christian centre (possibly the seat of the bishop) was built in town; a baptistery was added in the beginning of the 5th c. CE. After mid-5th c. CE the town was gradually abandoned. Cemeteries stretch along all the major routes leading from Emona towards other major settlements in the region. Gosposvetska cesta is a recently excavated part of the large northern cemetery of Emona where a surprising number of people were buried in sarcophagi, thus representing a wealthier part of the population of Emona. After the town area of Emona was abandoned a cemetery was established at Dravlje, ca. 3 km to the northwest, along the road to Carinthia. The settlement was not discovered but is presumed nearby. Due to the characteristic gravegoods the cemetery was traditionally ascribed to a group of Ostrogothic incomers guarding the road and dated to the period of the Ostrogothic kingdom (493-526).

Between mid-6th and mid-8th c. CE there is a gap in the settlement of the Emona basin. Among the earliest early Medieval sites in the area a cemetery of 14 graves dated traditionally to the 9th-10th c. CE was excavated in 2015 in present day Ljubljana at Hrušica, Litijska cesta, 5 km to the southeast of Emona.

##### Site: Gosposvetska cesta

The site at Gosposvetska cesta (Gosposvetska street) is a part of the large northern necropolis of Roman Emona along the road to Celeia. 337 graves were excavated between 2017-2018, dated between second third of the 4th - first decades of the 5th c. CE.
The individuals were buried in simple grave pits, wooden coffins, chests made of stone plates, sarcophagi or in tombs. Most of the graves were singular (282), while two to eight individuals were buried together in the rest of the graves. Of the 391 individuals, one third were subadults, two thirds were adults, with a 38 : 47 male to female ratio. Beside various dental and joint diseases, signs of malnutrition and occasionally healed fractures were noted.

44 individuals were selected for sampling (12%), 15 subadults and 29 adults. After genetic analysis there are 16 adult females, 13 adult males, 8 subadult females and 7 subadult males among the sampled individuals. In 8 cases genetic sex differs from the anthropologically assessed one, most probably due to poor preservation of the skeletal material.

Out of 44 sampled individuals 8 form three small pedigrees, P1, P2 and P3, the rest are unrelated.

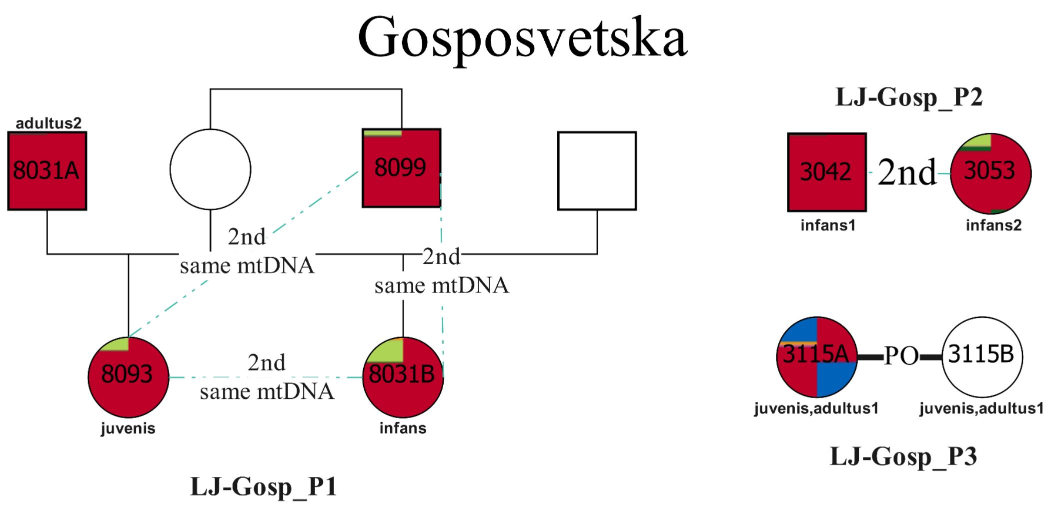


**Figure S5: Pedigrees found at Gosposvetska. Squares indicate males, while circles indicate females. Individuals are colored based on their fastNGSadmix results, with the modern panel results on the left and the ancient panel results on the right (Extended Data Fig. 1). The connection PO represents a parent offspring pair where the polarity is unknown. Unknown relationships are represented using dashed lines labeled with the degree.**

**P1**: unsampled female and her two reproductive partners, one is unsampled, the other is adult male 8031A. In the pedigree there is their subadult daughter 8093 and another subadult daughter of the unsampled female and the unsampled partner, subadult 8031B. She was apparently buried together with her stepfather. Pedigree is completed with the subadult brother of the mother, 8099.

**P2**: subadult male 3042 is 2nd degree related to subadult female 3053.

**P3**: mother and child buried together, adult female 3115A and subadult female 3115B.

##### Site: Dravlje

Three kilometres to the northwest from the Roman town of Emona, in the Ljubljana town quarter called Dravlje, a cemetery dating to the late 5th and early 6th c. CE was discovered during construction work in 1968. Several graves were destroyed and undocumented and the remaining 49 graves do not represent the whole cemetery. The graves belong to an undiscovered settlement, situated along the Emona-Carnium road most probably to supervise the area after the end of life in the town.

Burials consist of simple pits with parts of wooden coffins or tree trunks detected in some cases. Individuals were buried on their back with hands outstretched or sometimes on the pelvis. Several graves were furnished, females usually with ear rings, brooches and beads, males with belt buckles and combs. Archaeological and C14 dating points to the time around 500 CE (Table S3).

Most graves were singular, only in one grave two individuals were buried together (graves 23 and 24). Out of 49 individuals (29 adults and 20 subadults) 31 were sampled and analysed (63,27%). Out of the 31 sampled individuals 19 adults presented a balanced ratio with 11 males to 8 females. Out of 12 subadults 8 were genetic females and 4 males. Besides various dental and joint diseases, and possible malnutrition, interestingly congenital torticollis, neoplasms and possible tuberculosis were noticed. Two cases of pyramidal molars, commonly found in Asian and only rarely in European populations, and one confirmed case of ACD (grave 38), with two additional possible ACD (graves 17 and 42) were detected.

Pedigrees and relatedness

Out of 31 sampled individuals 17 were related (54,84%), 3 adult females, 7 adult males, 5 subadult females and 2 subadult males. 14 individuals were unrelated; 4 adult males, 5 adult females, 2 subadult males and 3 subadult females. One adult female (Dravlje_28) is related ($2^{nd}$ degree) to an adult female from another site, Rifnik in Celeia region (Rifnik_105).

Only one larger pedigree was detected among the people buried at Dravlje (P1).

Apart from P1 there is one nuclear family (P2), three sets of fathers with subadult children (P3, P4, P5, P6) and a pair of adult sisters (P7).

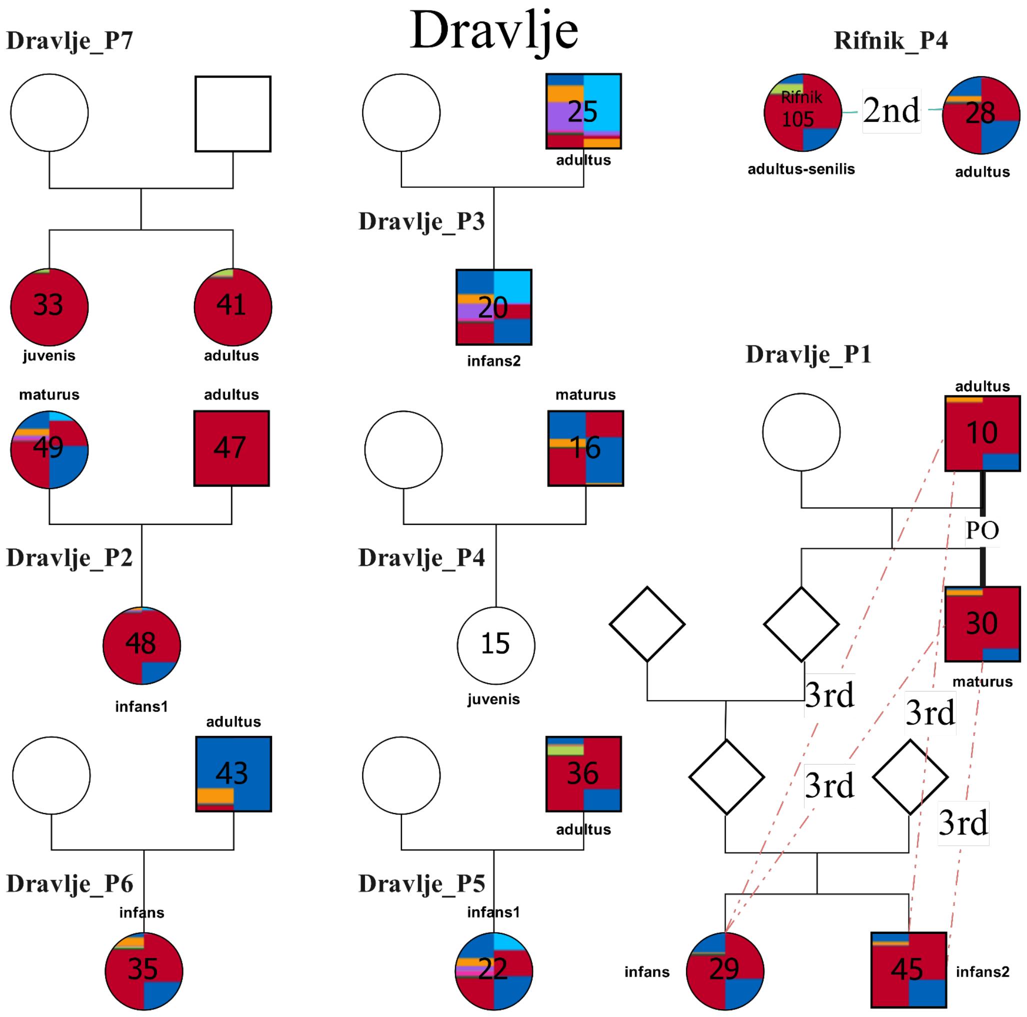


**Figure S6: Pedigrees found at Dravlje. Squares indicate males, while circles indicate females. Individuals are colored based on their fastNGSadmix results, with the modern panel results on the left and the ancient panel results on the right (Extended Data Fig. 1). The connection PO represents a parent offspring pair where the polarity is unknown. Unknown relationships are represented using dashed lines labeled with the degree.**

**P1**: consists of an adult male 10, his adult son 30 and a pair of his (10's) great-grandchildren, subadult female 29 and male 45.

**P2**: consists of adult male 47, adult female 49 and their subadult daughter 48.

**P3**: consists of adult male 25 and his subadult son 20

**P4**: consists of adult male 16 and his subadult daughter 15

**P5**: consists of adult male 36 and his subadult daughter 22

**P6**: consists of adult male 43 and his subadult daughter 35

**P7**: consists of adult sisters 33 and 41

Adult female Dravlje 28 is related (2$^{nd}$ degree) to adult female Rifnik_105.

Individuals in graves 1, 2, 18, 19, 23, 24, 28, 31, 32, 34, 38, 40, 42, 44 are unrelated.

**Site: Hrušica – Litijska cesta**

5 km to the southeast of Emona a small cemetery of 14 graves (Hrušica, Litijska cesta) was excavated in 2015. The cemetery is limited to the east and west but could continue to the north and south. Graves are oriented E-W. Some of the burials consisted of wooden coffins or tree-trunks; two graves had stone lining. Some graves were simple pits.

No settlement remains were discovered in the vicinity, the cemetery would have belonged to a small early Medieval community in a period before the first beginnings of a medieval town of Ljubljana were starting to appear below the Castle hill, on the opposite bank of Ljubljanica River to the abandoned Roman town. Radiocarbon dating of the graves shows a span between early 8th and late 9th c. CE for most graves, some are even earlier (grave Hrušica_1007, late 7th-mid 9th c. CE) (Table S3). Only one grave is richly furnished, adult female Hrušica_1007, with silver and copper alloy earrings, beads, knife and a needle-case. The adult female Hrušica_1006 was buried with two sets of copper alloy head rings. Other graves contain iron knives, a pot and chicken bones or are unfurnished (children Hrušica_1004, Hrušica_1012-1, Hrušica_1012-2 and adults Hrušica_1001, Hrušica_1011, Hrušica_1008, Hrušica_1009).

Altogether the graves contained 14 individuals with an unbalanced male-to-female ratio: 4 adult females, 7 adult males and 3 subadult females. Subadults represented one fifth of individuals. There is a discrepancy between the results of osteological sex estimation and genetic sex determination in 2 cases, one osteological male is genetic female, and one osteological female is genetic male, the ratio stays the same. Grave Hrušica_1014 only contained three parts of long bones and was not sampled. The other 13 graves contained 14 skeletons which were all sampled and analysed. Out of 14 analysed individuals 11 (78.57%) showed biological relatedness with other members of the population, this group includes 5 females (2 adults and 3 subadults) and 7 adult males. Among the 3 unrelated adult individuals there are 2 females and 1 male.

Pedigrees and Relatedness

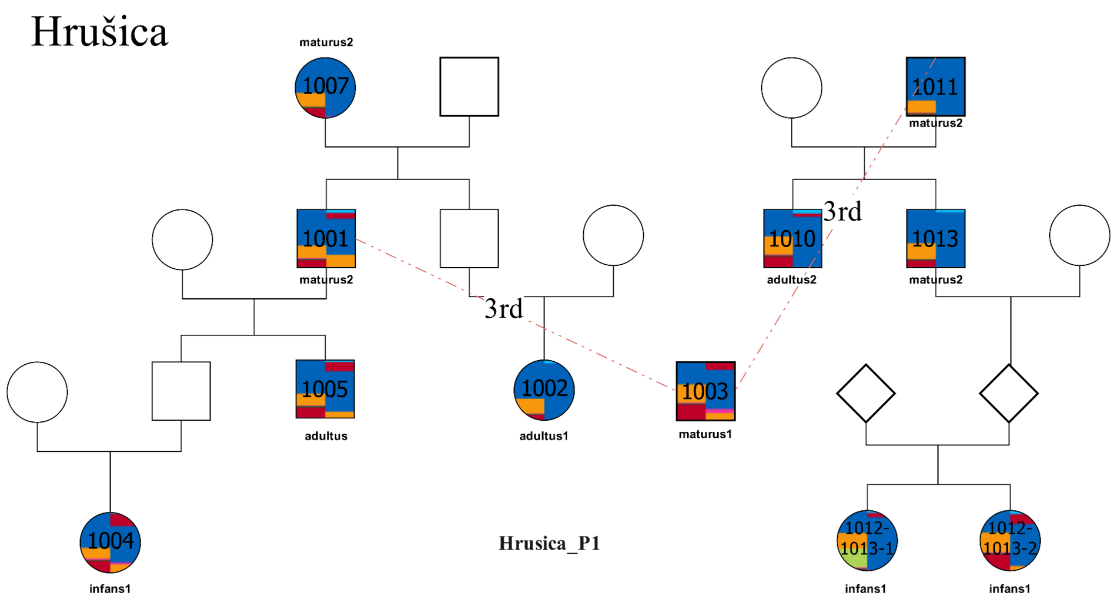


**Figure S7: Pedigrees found at Hrušica. Squares indicate males, while circles indicate females. Individuals are colored based on their fastNGSadmix results, with the modern panel results on the left and the ancient panel results on the right (Extended Data Fig. 1). The connection PO represents a parent offspring pair where the polarity is unknown. Unknown relationships are represented using dashed lines labeled with the degree.**

Genetic analysis identified one pedigree (**P1)** to which all the related individuals belong. P1 spans at least 4 (possibly 5) generations at the site. Grave 1003, an adult male, connects two subpedigrees (SP1 and SP2) of more closely related individuals as he is 3rd degree related to both.

**SP1**: is composed of an adult female 1007 (unrelated to 1003), her son 1001 (3rd degree related to 1003) and her two grandchildren, adult male 1005 (son of 1001) and an adult female 1002. The last generation is represented by a niece of 1005, subadult 1004.

Members: Hrušica_1007; Hrušica_1001; Hrušica_1005; Hrušica_1002; Hrušica_1004

**SP2**: is composed of an adult male 1011, his two adult sons 1010 and 1013 and two subadult granddaughters of 1013, 1012-1 and 1012-2. One generation, the parents of the granddaughters, is missing as are all the mothers.

Members: Hrušica_1011; Hrušica_1010; Hrušica_1013; Hrušica_1012-1; Hrušica_1012-2

Unrelated graves are all adults, females Hrušica_1006 and Hrušica_1009 and male Hrušica_1008.

**Section 3. Vipavska dolina region case study**

Vipavska dolina (valley of River Vipava) is a corridor leading from Italy and the outflow of the Soča/Isonzo River towards inland, the high Karst plateaus and central Slovenia. It gravitates towards the Friuli plain in Italy and shares most of its history. The Roman itinerary road from Italy to Pannonia runs along the valley. In contrast to the other three study areas chosen for this paper, Vipavska dolina was continually settled between the Roman period and the Middle ages. The main Roman settlement in the valley was the fort Castra (Ajdovščina) which probably represented the headquarters of the Late Roman defence system of the northeastern access to Italy. It functioned until the end of the 5th c. CE. A small cemetery of 9 graves was discovered between 2009-2013 at Miren near the modern settlement of Nova Gorica, close to the Roman road and ca. 25 km to the west from Castra. The furnished female graves and the radiocarbon dating allow for a dating to around 500 CE. Culturally the site was interpreted as belonging to the Ostrogothic kingdom. About 10 km to the north from Miren on the left bank of the Soča/Isonzo river a cemetery of 53 graves was excavated in Solkan. The graves were dated to the 4$^{th}$-5$^{th}$ c. and the late 6$^{th}$-7$^{th}$ c. CE. The later graves are ascribed to a Langobard garrison. Both Miren and Solkan cemeteries probably belonged to groups settled in the vicinity to guard the road or/and the Soča/Isonzo River crossing. About 2 km from Solkan a small group of graves was found in the ruins of late Roman buildings at Ledine in Nova Gorica. They are roughly contemporary but do not display any of the Langobard-period gravegoods like the Solkan cemetery. One of the graves is radiocarbon dated to the 7th-8th c. CE.

The main late antique settlement of Vipavska dolina developed on the hill of Sv. Pavel above the village of Vrtovin. The large rocky promontory with an excellent visual control of the valley below was protected by a defence wall, the settlement consists of stone-built houses on terraces, one or more churches and a tower protecting a water spring below the defence wall. The site was excavated to a very limited extent and only 4 graves were discovered, three belong to the 6th-7th c. CE and the fourth one to the 8th-9th c. CE. Use of the settlement is confirmed for the 6th c. CE but a longer use into the early Middle ages is possible according to individual finds from the site. Under the hill in the village of Gojače remains of an early Medieval settlement and cemetery were discovered. Most graves were destroyed but a dating to the 8th-9th c. is confirmed by small finds and radiocarbon dating of one grave.

Outside of the walls of Castra (Ajdovščina), next to the church of St George (sv. Jurij), a cemetery of 25 graves, originally dated to the 9th-11th c. CE and ascribed to the Slavic population was excavated in 2012. Radiocarbon dating revealed the graves belong to early 11th c (Table S3).

**Site: Castra**

Castra (Ajdovščina) was a fort within the Late Roman Claustra Alpium Iuliarum defence system, possibly representing its headquarters and definitely the supply base. The cemetery of the settlement continues from the 1st to the 5th c. CE. We selected a burial group of 44 graves for sampling (altogether 26 individuals could be sampled due to poor preservation), presumably the latest skeleton graves out of the western necropolis of Castra. The cemetery included a high ratio of multiple burials as well as cremations, its original size cannot be accurately reconstructed because of rare excavated areas in a densely built-over town. Only a small number of graves was furnished with beads, brooches, finger- and ear rings, oil lamps, pottery and 4th c. CE coins.

In 44 graves 55 individuals were buried. Due to poor preservation, one third of individuals remained of undetermined age and sex during the anthropological analysis in situ. Despite this there is a very small number of subadults, 4 altogether. Out of 40 adults anthropological analysis determined 12 females (2 questionable) and 10 males (1 questionable).

26 individuals were selected for sampling (46%). After genetic analysis there are 9 adult females, 13 adult males, 2 subadult females and 2 subadult males in the cemetery. In 9 cases genetic sex differs from the anthropologically assessed one, most probably due to poor preservation of the skeletal material.

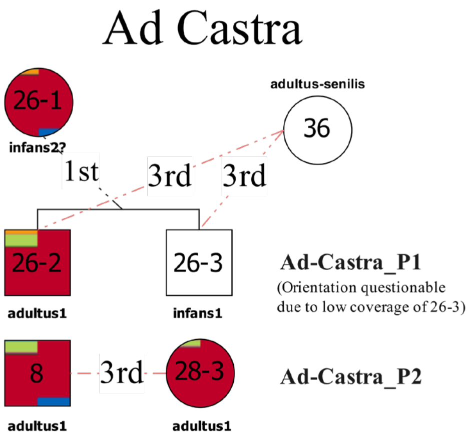


**Figure S8: Pedigrees found at Castra. Squares indicate males, while circles indicate females. Individuals are colored based on their fastNGSadmix results, with the modern panel results on the left and the ancient panel results on the right (Extended Data Fig. 1). The connection PO represents a parent offspring pair where the polarity is unknown. Unknown relationships are represented using dashed lines labeled with the degree.**

Out of 26 individuals only 6 are related. Four individuals form pedigree 1 and two form pedigree 2.

**P1**: In grave 26 a subadult female 26-1 is buried together with an adult male 26-2 and a subadult male 26-3 to whom she is first-degree related, but the relationship is unclear due to low coverage. Adult female 36 is related (3rd degree) to the brothers on the father's side. This pedigree is radiocarbon dated to (late 3rd)/late 4th-mid 6th c. CE (Table S3).

**P2**: Adult males 8 and 28-3 are 3rd degree related.

**Site: Miren**

Near the modern settlement of Nova Gorica, a group of 7 graves was discovered in 2011 at Miren, one additional grave was discovered in 2009 and another one in 2013 (grave 8 in this study). All preserved adult graves are furnished, subadult graves are not. The furnished female graves and the radiocarbon dating allow for a dating to around 500 AD (Table S3).

All the graves were singular and contained seven adults and two subadults with a balanced male (3) to female (3 plus 1?) ratio. In two female adults, signs of artificial cranial deformation were documented (graves 1 and 7) and in one adult male (grave 5) a well-healed fracture below the left elbow.

6 out of 9 graves were sampled (67%), after genetic analysis: 2 adult females, 3 adult males and one subadult male.

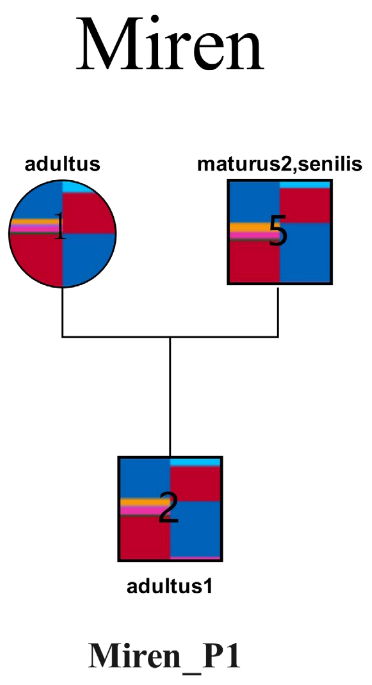


**Figure S9: Pedigrees found at Miren. Squares indicate males, while circles indicate females. Individuals are colored based on their fastNGSadmix results, with the modern panel results on the left and the ancient panel results on the right (Extended Data Fig. 1). The connection PO represents a parent offspring pair where the polarity is unknown. Unknown relationships are represented using dashed lines labeled with the degree.**

Three adults, Miren_1, Miren_2 and Miren_5, two males (father and son) and a female (mother), form a pedigree (**P1**), all other individuals are unrelated.

**Site: Solkan**

On the left bank of the Soča/Isonzo river a cemetery of 53 graves was excavated in Solkan in 1980 and 2008. Additional 19 graves were found in 2023, but they are not included in our study.

The graves were initially dated to two phases, the 4$^{th}$-5$^{th}$ c. and the late 6$^{th}$-7$^{th}$ c. It is as yet unclear whether these are the remains of two cemeteries with a temporal hiatus or a continuously used one. Some of the graves initially dated to the late Roman period form part of the pedigrees with certainly later individuals so their dating was corrected. A number of graves were equipped with weapons and belt sets, typical of Langobard Italy in the 7$^{th}$ c. while female graves are unremarkable regarding grave goods. In one grave no remains were discovered. In 53 graves, 60 individuals were buried, as seven graves were double. Subadults, predominantly males, presented one fifth of individuals. Among adults, two thirds were males. Only noted pathological changes were two healed traumas and short stature (around 120 cm - 130 cm) in the case of one individual.

Only 35 individuals could be sampled due to poor preservation. 24 of them are related and form three pedigrees: P1, P2 and P3. 11 individuals are unrelated: 2, 6, 26, 32, 36, 41, 42, 43, 45, 49, 53.

**P1**: the largest pedigree consists of three subpedigrees and four more distantly related members as well as an Vrtovin individual. This is the largest pedigree in the whole study, and the most difficult pedigree to resolve

The first sub-pedigree consists of adult female Solkan_25 and her two adult sons Solkan_44 and Solkan_51.

The second sub-pedigree are the descendants of Solkan_25's siblings. There is her unsampled brother's adult son Solkan 31, his unsampled brother's subadult sons Solkan_37 and Solkan_38 and their mother's adult sister Solkan_22. Through Solkan_31's unsampled sister this sub-pedigree is connected to the third sub-pedigree as adult male Solkan_12 and Solkan_31's unsampled sister married unsampled siblings. Solkan_31's unsampled sister and her partner had two adult sons, Solkan_30 and Solkan_23. Solkan_23's subadult son Solkan_29 is 3rd degree related to adult male Vrtovin_1.

The third sub-pedigree consists of the mentioned adult male Solkan_12, his adult son Solkan_34 and subadult grandson Solkan_13.

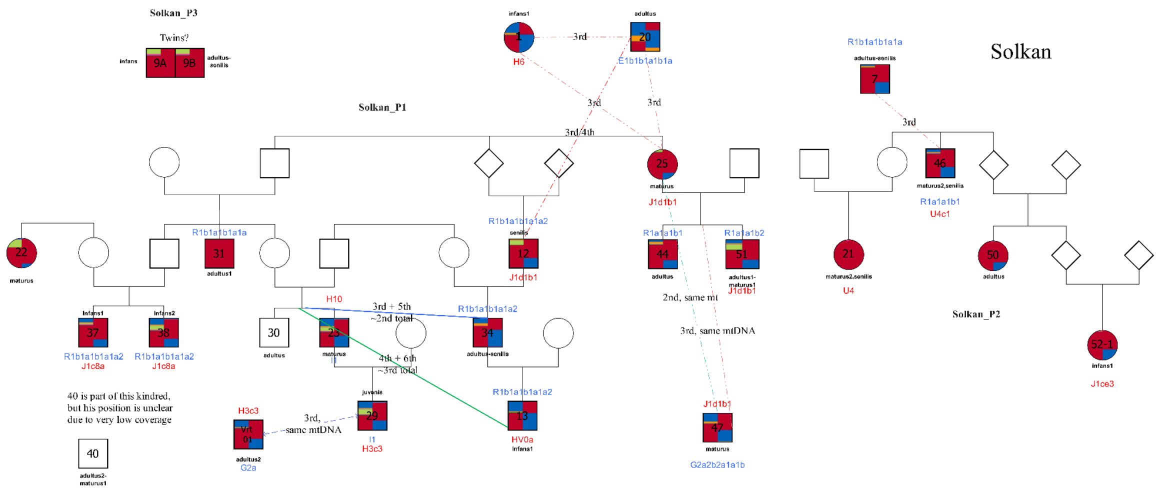

**Figure S10: Pedigrees found at Solkan. Squares indicate males, while circles indicate females. Individuals are colored based on their fastNGSadmix results, with the modern panel results on the left and the ancient panel results on the right (Extended Data Fig. 1). The connection PO represents a parent offspring pair where the polarity is unknown. Unknown relationships are represented using dashed lines labeled with the degree.**

Subadult female Solkan_1, adult male Solkan_20 and adult male Solkan_47 are 3rd degree related to pedigree members.

Adult male Solkan_40 is also part of this pedigree, but his position is unclear due to low coverage.

**P2**: consists of adult male Solkan_46, his adult niece Solkan_21 by unsampled sister, his adult niece Solkan_50 by unsampled sibling and this niece's subadult niece Solkan 52_2 by unsampled sibling. In the pedigree is also a 3rd degree relative of Solkan_46, adult male Solkan_7.

**P3**: Subadult male 9A and adult male 9B are twins, as the two graves are in superposition and stratigraphy suggests that 9B (adult) is earlier than 9A (infans), the possible mixing of bones can not be excluded. (Table S3).

The pedigrees have been radiocarbon dated and they span the time between late 6th to early 8th c. CE (Table S3).

**Site: Ledine**

In the present-day town of Nova Gorica, a group of 5 graves was dug into ruins of Roman buildings, dating to the 4th c. CE. The graves are preliminary dated to the 7th-8th c. AD. All the graves were singular, belonging to two subadults and three adults, a female and two males. All five individuals were preserved enough for sampling, one adult male (grave 3) is genetic female, subadults are a genetic male and a female.

Apart from no. 2 who is an unrelated male in an unfurnished grave, the rest of the graves form a pedigree (P1). It consists of an adult grandmother in grave 3, her adult daughter in grave 4 and her subadult daughter in grave 1. In grave 5 is a subadult male (unfurnished grave), the nephew of adult female in grave 4 on her brother's side. The three female graves are furnished, all have combs, female 3 also has a folding knife, a spindle whorl and an iron ring, female 4 also has a set of ear rings, a knife, a spindle whorl, a belt buckle and an iron ring. The subadult female in grave 1 was buried with a set of simple wire ear rings apart from the comb. Grave 1 is radiocarbon dated to 683-769 CE  (Table S3)..

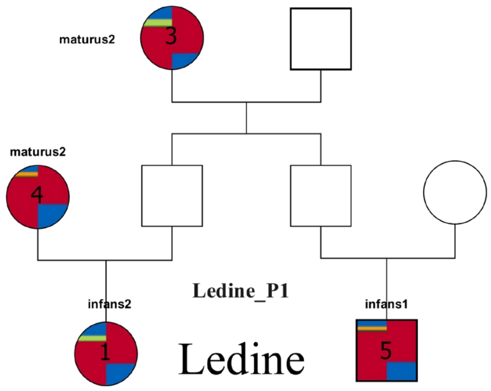


**Figure S11: Pedigrees found at Ledine. Squares indicate males, while circles indicate females. Individuals are colored based on their fastNGSadmix results, with the modern panel results on the left and the ancient panel results on the right (Extended Data Fig. 1). The connection PO represents a parent offspring pair where the polarity is unknown. Unknown relationships are represented using dashed lines labeled with the degree.**

**Site: Vrtovin**

Further along the Vipava valley towards the inland a strongly naturally defended hilltop site of St. Paul (sv. Pavel) above the village of **Vrtovin** dominates the valley. The hilltop fortification with defence walls, church(es), dwelling houses and a tower protecting a water source represents one of the largest and most impressive late antique and early Medieval hilltop sites in the region. Only a small part was excavated and within the settlement four graves were found, dated to the $6^{th}$-$7^{th}$ and $8^{th}$- 9th c. CE. Graves 1-3 were discovered in what is most probably a church. Grave 4 lay along the wall that divides the settlement in half, it is C14 dated to 764 - 894 CE  (Table S3).. Grave 4 was furnished (iron awl, iron arrowhead, lead tube) while the other three were not. All were singular graves of adults, two males and two of undetermined sex, all are radiocarbon dated. The two males were preserved enough for sampling (graves 1 and 4). Both are also genetic males and unrelated. Male in grave 1 is related ($3^{rd}$ degree) to subadult male 29 from Solkan.

**Site: Gojače**

Below the Sv. Pavel above Vrtovin hill a small settlement with wooden structures and twelve graves was discovered near the present-day village of Gojače. Six out of twelve graves were not archaeologically investigated, one has no anthropological data. In the remaining five graves, there were three subadults and two adults. Only two individuals were preserved enough for sampling: graves 11 and 12, and adult and a child, they are unrelated adult male and subadult male. Both graves were furnished, the adult with a comb, the child with a knife and two iron rings. Grave 12 is radiocarbon dated to 771-883 CE  (Table S3).

**Site: Šturje**

In present-day Ajdovščina, next to the church of St George (sv. Jurij) at Šturje (previously a separate village, first mention 1320), a cemetery of 25 graves, initially dated to the 9th-11th c. was excavated in 2012 (Table S3). It lies outside (to the east) of the walls of Castra, the Roman fortification and on the other side of the Hubelj River. The cemetery probably extends further towards the east and south. Graves were dug into a dark layer of Roman ruins which also filled the graves, therefore the grave pits were not visible. In the southern part of the cemetery the graves are highly concentrated and later graves often damaged or completely destroyed earlier ones. Some graves were lined with stones and in some remains of wooden coffins were noticed. Out of the 25 graves 13 were furnished (52%), 8 adults (2 adult males, 5 adult females, for grave 17 there is no genetic sex data, anthropological male) and 5 subadults (2 males and 3 females).

Out of 25 graves 22 (88%) were sampled and analysed. Bones in graves 2 and 11 (subadults) and 17 (adult male) were too poorly preserved to be sampled. Out of the 22 sampled individuals 11 were adults with an unbalanced ratio of 3 males to 8 females. Out of 11 subadults 4 were genetic females and 7 males.

Pedigrees and relatedness

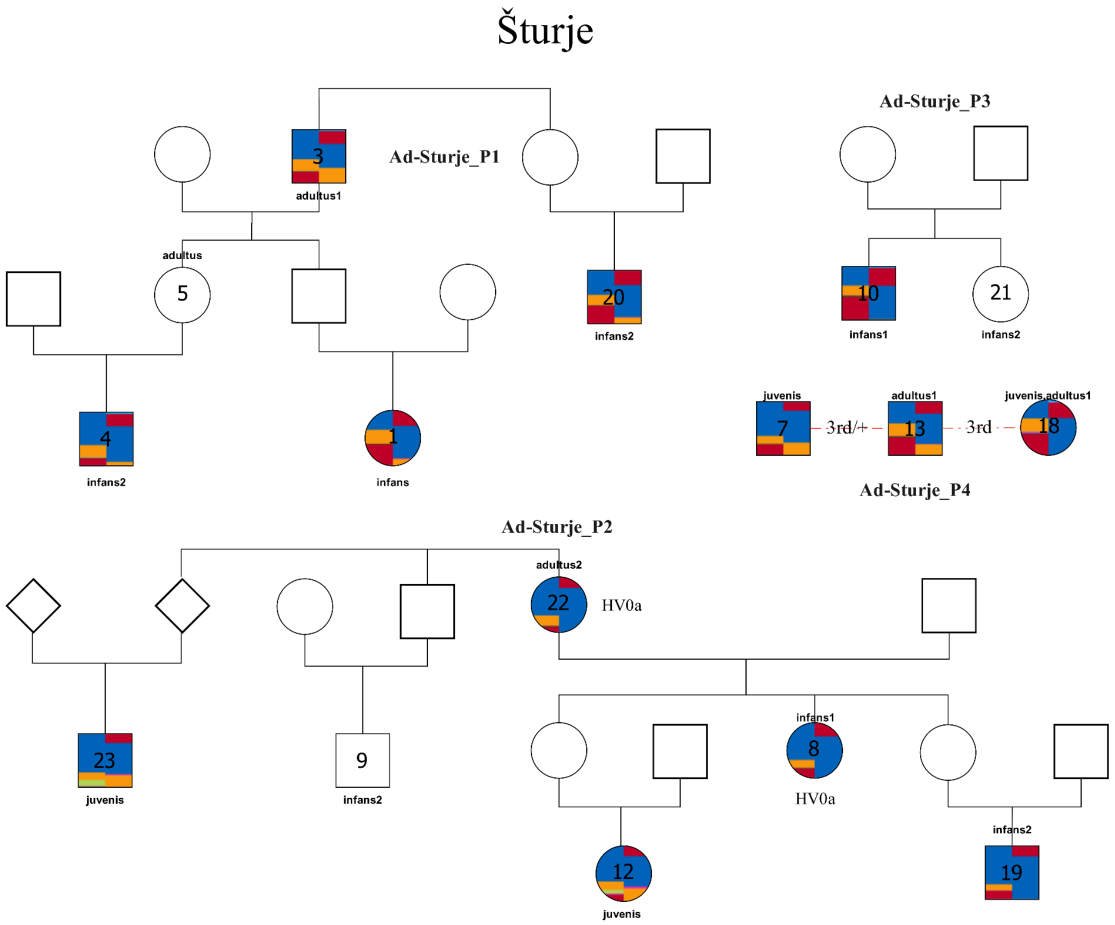

**Figure S12: Pedigrees found at Šturje. Squares indicate males, while circles indicate females. Individuals are colored based on their fastNGSadmix results, with the modern panel results on the left and the ancient panel results on the right (Extended Data Fig. 1). The connection PO represents a parent offspring pair where the polarity is unknown. Unknown relationships are represented using dashed lines labeled with the degree.**

Out of 22 sampled individuals 16 were related (72,73%): 4 adult females, 5 adult males, 3 subadult females and 4 subadult males. 6 adult individuals were unrelated; 2 adult males and 4 adult females.

Four pedigrees were detected at Šturje, P1-P4.

**P1**: consists of adult male 3, his subadult nephew 20, his (Šturje_3) adult daughter 5 and her subadult son 4 and her subadult niece 1.

**P2**: consists of adult female 22, her subadult daughter 8, her (Šturje_22) subadult grandchildren female 12 and male 19, and her (Šturje_22) subadult nephews 9 and 23.

**P3**: consists of subadult brother 10 and sister 21.

**P4**: consists of loosely related (3rd degree +-) subadult male 7, adult male 13 and adult female 18.

Individuals in graves 6, 14, 15, 16, 24, 25 are unrelated, all except grave 15 (adult male) are adult females.

Both related (8 out of 16, 50%) and unrelated (4 out of 6, 67%) individuals received furnished burials. The unrelated female in grave 16 received the richest grave goods, a necklace of 306 glass beads, 2 head rings and a finger ring. All pedigrees contained furnished and unfurnished burials. Most adult and subadult females were buried with 2 head rings, one (grave 22) with 4.

**Section 4. Dolenjska and Bela krajina region case study**

The fourth studied region lies partly along the main Roman road leading from Italy towards the eastern part of the Empire. Bela krajina with the site of Črnomelj lies along a communication route towards the Adriatic coast. This way the main land communications leading towards Italy and the sites along them from the northeast are represented in the study. The late Roman period is represented by the cemetery of the fortification at Črnomelj. In the 6th c. several hilltop sites developed in the hinterland of the major roads and along lesser routes which were easier to protect, Vrajk near Mokronog and the sites in the Gorjanci hills, Zidani gaber and Gradec above Mihovo. Less than 3 kilometres from Mihovo, the early Medieval cemetery of Camberk is dated to the 9th c. CE. It was traditionally tentatively ascribed to the autochthonous population due to the closeness of the Gorjanci settlement nucleus.

**Site: Zidani gaber**

Zidani gaber is a fortified hilltop settlement in the Gorjanci hills. It lies on the so-called Laška or Vlaška pot, a road that has connected both sides of the hills probably since prehistory. The site was used between the 3rd-9th c. CE, with the most intensive settlement phase in the 6th c. The position, artificial and natural defences and small finds speak for a military and strategic role of Zidani gaber. Remains of a church were partly excavated while several graves had been discovered in the late 19th and early 20th c., but no skeletal material or info on the location of the graves is preserved. In 1999 a sarcophagus was discovered during work in the woods. It was located in a building to the south of the late antique church and contained a skeleton of a 24-28 year-old female which was sampled for this study.

**Site: Gradec above Mihovo**

In 1987-1989 a small late antique church of a simple ground plan was discovered on an elevated plateau in the Gorjanci hills, a 20-minute walk away from Zidani gaber. It may represent the cemeterial church of Zidani gaber or a separate settlement. In it and around the apse 6 graves were found, one in a stone sarcophagus. All 6 graves were sampled for analyses.

An adult male was buried unfurnished in a stone sarcophagus inside the church, an older Roman brooch lay on the lid. The grave was radiocarbon dated to 425-565 CE. Around the apse there were three adult females Gradec *3,* Gradec_5 and Gradec_6, an adult male Gradec_2 and a subadult male Gradec_4. Graves Gradec_2 and Gradec_6 were radiocarbon dated to 382-541 CE and 257-531 CE and contained a belt buckle and an ear ring & beads respectively.

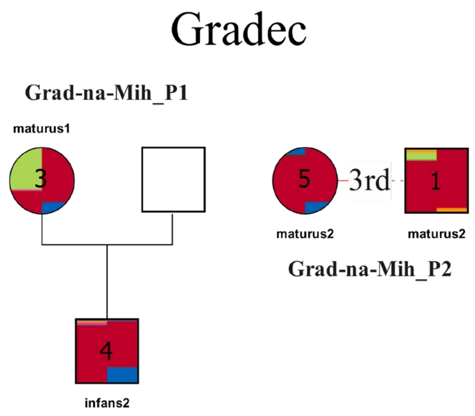


**Figure S13: Pedigrees found at Gradec. Squares indicate males, while circles indicate females. Individuals are colored based on their fastNGSadmix results, with the modern panel results on the left and the ancient panel results on the right (Extended Data Fig. 1). The connection PO represents a parent offspring pair where the polarity is unknown. Unknown relationships are represented using dashed lines labeled with the degree.**

Adult female Gradec_3 and subadult male Gradec_4 are mother and son (**P1**). Adult female Gradec_5 and adult male Gradec_1 (sarcophagus) are 3rd degree related (**P2**).

**Site: Vrajk near Mokronog**

Rescue excavations in 1996 at Vrajk in Gorenji Mokronog uncovered a partly destroyed inhumation cemetery with fifteen preserved and five destroyed or partly preserved burials. The northwestern edge of the cemetery was excavated. The cemetery probably extends further to the southeast, while the western and northern sections of the cemetery were entirely destroyed. Very modest grave goods (basket earrings, belt buckle, beads, two stemmed glass feet) led the excavator to date the cemetery to the 6th-7th c. CE, but radiocarbon dating showed mid-5th-mid-6th c. CE (Table S3). Ca. 300 m to the SE of the site lies the castle of Gorenji Mokronog with remains of a late antique settlement on the terraces. It is presumed the cemetery belonged to this settlement, which remains unpublished.

Based on anthropological analysis, 17 preserved skeletons consisted of two children, eight males and seven females. The children, who died at ages two and four, represented 11.8% of the skeleton series. The age of the majority of the adult skeletons could be assessed only in the framework of age categories. Two individuals died as juveniles, a 15 year old girl and a 19-20 year old youth, possibly also a woman that died between the ages of 15 and 30 years. Most of the individuals, 10, died as young and mature adults, while two died as an old adult.

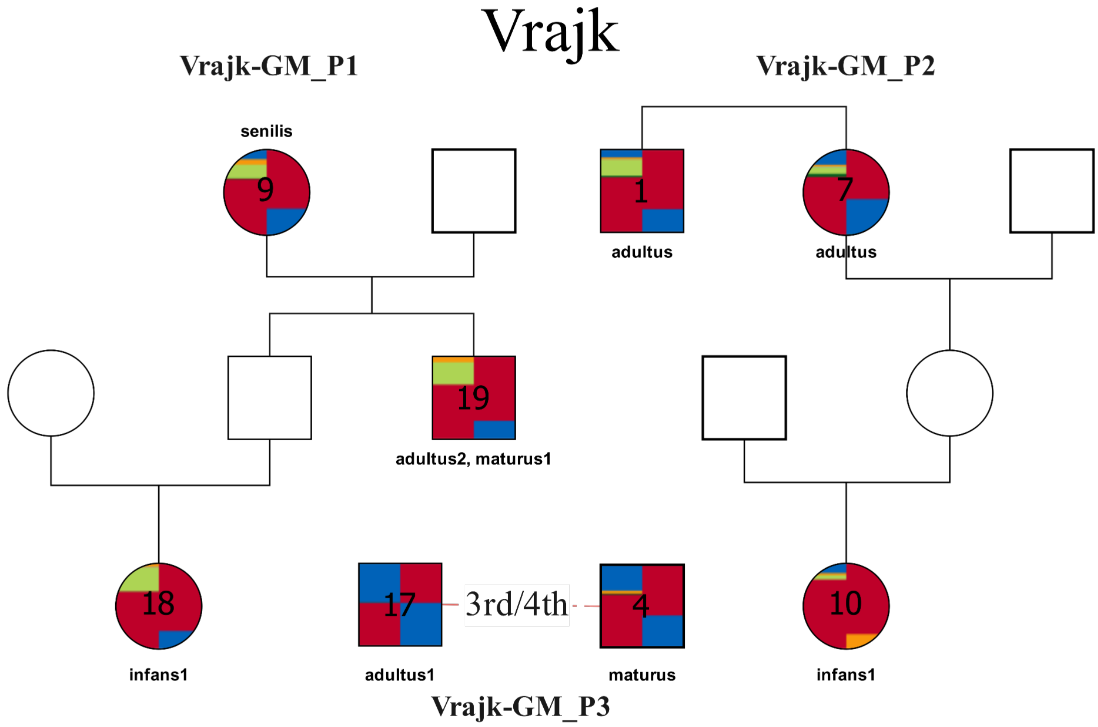


**Figure S14: Pedigrees found at Vrajk. Squares indicate males, while circles indicate females. Individuals are colored based on their fastNGSadmix results, with the modern panel results on the left and the ancient panel results on the right (Extended Data Fig. 1). The connection PO represents a parent offspring pair where the polarity is unknown. Unknown relationships are represented using dashed lines labeled with the degree.**

Eight individuals are related, forming three pedigrees: P1, P2 and P3.
**P1** consists of adult female Vrajk_9, her adult son Vrajk_19 and her subadult granddaughter Vrajk_18 (parents not sampled). **P2** consists of adult male Vrajk_1, his adult sister Vrajk_7 and her subadult granddaughter Vrajk_10. **P3** consists of 4th degree related adult males Vrajk_4 and Vrajk_17.

**Site: Camberk**

In 2002 and 2004 rescue excavations uncovered 36 graves in the quarry of Camberk above Cerov log, in Gorjanci hills. The graves were dated to the 8th-9th c. CE on the basis of relatively rare grave goods (ear rings, finger rings, belt buckle, knives, pottery), radiocarbon dating pushed the date to mostly 9th c. CE. A settlement was perhaps located further along the ridge to the east but it has not been investigated and is now destroyed by the quarry. Its proximity to the sites of Zidani gaber and Gradec may indicate the continuation of use of the same location, positioned on and near to the road across the Gorjanci hills.

Out of 36 graves, three were destroyed. In the rest, 18 individuals were children or juveniles, and 15 were adults, of which five females, five males and five of undetermined sex. Remains of 22 individuals were preserved enough for the sampling and analyses, 11 children, one juvenile and ten adults. Out of 11 children, four were genetically females and seven were males, juvenile was determined as a female. Five of the adults were females and five males.

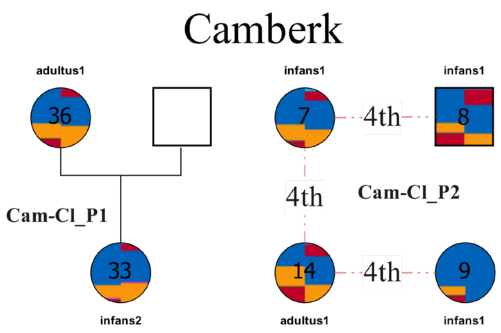


**Figure S15: Pedigrees found at Camberk. Squares indicate males, while circles indicate females. Individuals are colored based on their fastNGSadmix results, with the modern panel results on the left and the ancient panel results on the right (Extended Data Fig. 1). The connection PO represents a parent offspring pair where the polarity is unknown. Unknown relationships are represented using dashed lines labeled with the degree.**

Genetic analyses show a very low degree of relatedness, only one PO pair: adult female Camberk_36 is the mother of the subadult female Camberk_33 (**P1**). Apart from this PO pair there is a group of 4 loosely related individuals: subadult female Camberk_7 (three glass pendants), subadult male Camberk_8, subadult female Camberk_9 (earrings and a finger ring) and adult male Camberk_14 (**P2**). The three little girls with jewellery are 3 out of only 4 graves with jewellery in the whole site. (Only one unrelated adult female, Camberk_17 (18-20yrs), was buried with ear- and finger rings).

The adult male in grave Camberk_22 is a genetic outlier, he is also buried some distance away from the rest, his grave was partly lined with stones. He probably limped due to a poorly healed fracture of a femur and had problems with his teeth and neck.

**Site: Črnomelj**

Črnomelj is a small town located at the centre of Bela krajina, the extreme southeastern part of Slovenia. Between 1988-1997 rescue excavations in the historic town centre revealed traces of defence walls and towers, probably a small church and a cemetery. Črnomelj was a late Roman and antique fortified settlement, naturally defended by two rivers on three sides, Dobličica and Lahinja. On the bank of the river Lahinja a cemetery of 27 graves was excavated. The cemetery was initially dated to the 6th century CE as the graves cut and therefore postdate the cobbled surface, which can be assigned to the 5th and early 6th c. CE on the basis of the associated ceramic assemblage. The single piece of metalwork from the surface, a silver belt buckle, can be dated to the 6th century. However, the few grave goods are typologically earlier, late Roman. Radiocarbon dating of the graves confirmed they belong to the late Roman period  (Table S3)..

Out of 28 graves, 20 were available for anthropological analysis. 13 individuals were adults and seven children or juveniles. Among adults, three were assessed as females, eight as males and two as undetermined. Remains of 15 individuals were preserved enough for sampling, two children, two juveniles and 11 adults. Both children were genetically female, one juvenile was female and one male. Three of the adults were females and eight were males.

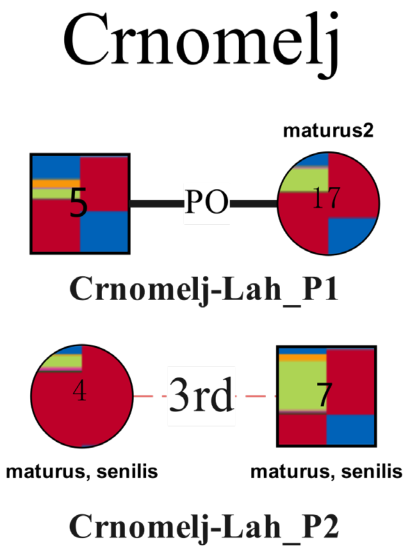


**Figure S16: Pedigrees found at Črnomelj. Squares indicate males, while circles indicate females. Individuals are colored based on their fastNGSadmix results, with the modern panel results on the left and the ancient panel results on the right (Extended Data Fig. 1). The connection PO represents a parent offspring pair where the polarity is unknown. Unknown relationships are represented using dashed lines labeled with the degree.**

Only four individuals are related. There is one P-O pair of adult males Črnomelj_5 and Črnomelj_17 (**P1**) while adult female Črnomelj_4 and adult male Črnomelj_7 are 4th degree related (**P2**).

**Comparative sites: Cividale**

**Site: Cividale Ferrovia**

Cividale Ferrovia represents one of the most illustrative, albeit partial, open-field funerary area from the Longobard period located within the periurban belt of Forum Iulii. In 2012, the excavation of the cemetery revealed a total of 79 tombs (73 with human remains and 6 empty), resulting in 81 individuals: 52 adults, 22 sub-adults, and 7 infants (ages 1–3 years). The sex ratio was fairly balanced: 25 males and 22 females; 5 individuals were undetermined due to young age, lack of grave goods, or poor skeletal preservation.

Graves were simple pit burials often lined with river cobblestones, forming bottom or side linings. This style is consistent with other Cividale cemeteries and reflects the use of locally available fluvial material. Many graves also showed signs of wooden structures, likely planks used to line the pit or cover the burial. The social structure appears highly articulated: some graves feature symbolic objects like combs, knives, pottery, amulets—especially among children; others lack any items (around 20%), suggesting status differences or functional distinctions within family groups.

The cemetery was in use for approximately one century, and three phases can be recognized:

- Phase I: ca. 590–630
- Phase II: ca. 630–670
- Phase III: ca. 670–700+

Each cluster contains burials spanning all three periods, suggesting that family groups remained in their designated areas for multiple generations.

Prestigious male burials were often armed and positioned centrally within each cluster, potentially indicating heads of family. Grave goods included: Swords, spears, shields (often decorative), silver or bronze belts, buckles, and crosses, horse-riding accessories, like spurs.

Wealthier female graves featured: fibulae (S-shaped and stirrup-shaped); gold pendants, earrings, and glass beads; comb artifacts, often ornate or encased; gold crosses with intricate decorations. A shift in fashion and funeral practices is noticeable over time, with monili (jewelry) becoming less common in later phases.

Pottery and glass vessels were found mainly in Phase I, often in children’s and female burials.

Animal-related items, like deer antlers and horse bones, suggest symbolic or ritualistic functions connected to ancestral memory or protection in the afterlife.

Gold crosses were discovered in seven graves and appear to reflect both religious beliefs and social prestige.

Combs were the most common offering across sexes and ages, possibly symbolic of personal grooming or spiritual cleansing.

The presence of similar decorative motifs across multiple crosses and belt sets suggests localized workshops or shared molds, especially in Cividale. However, some items indicate broader artisan networks, with designs found in other regions of Italy and even beyond the Alps, reflecting interregional exchange.

Borzacconi A. and Giostra C. 2018, “La necropoli presso la ferrovia a Cividale del Friuli” in Citta e campagna: culture, insediamenti, economia (secc. VI-IX), Archeologia barbarica II. SAP Società Archeologica.

Borzacconi A., Saccheri P., Travan L. 2023, "Due sepolture anomale dagli scavi della necropoli longobarda "della ferrovia" a Cividale del Friuli in Pergola P., Roascio S., Dellu' E. (eds) SIT TIBI TERRA GRAVIS, Sepolture anomale tra età medievale e moderna, Atti del Convegno internazionale di Studi, Albenga (SV) - Palazzo Oddo, 14-16 ottobre 2016, Archaeologies, histories, islands and borders in the Mediterranean 12, ISBN 978-1-80327-475-1; ISBN 978-1-80327-476-8 (e-Pdf)

**Site: Cividale San Mauro**

The necropolis of San Mauro in Cividale del Friuli was discovered in 1886. After a first excavation campaign in 1982 along the slope of San Mauro Hill, a systematic excavation campaign was initiated in 1994 by the Superintendency for Archaeological Heritage of Friuli Venezia Giulia. Altogether 22 burials with 23 individuals dated to the Langobard period came to light. The tombs were found at varying depths. Tombs are laid out in three main rows—two uphill and one downhill. All tombs were simple earth pits, with no wooden structures identified. In grave 21 stone coverings were found: one layer directly above the body and a second layer above.

Male burials typically contain weapons—such as the *spatha*, knife (or *scramasax*), spear and shield—and clothing-related items like iron or bronze belt buckles. Shears are also commonly found, possibly related to artisanal activities. Additionally, spurs, stirrup fittings, and buckles appear in the burials, serving both decorative and functional purposes. Some burials also included common ceramic vessels and coins.

Female grave goods consist of buckles, gilded bronze clasps, necklaces, earrings, rings, and brooches, along with ceramic and glass containers. Gilded buckles were found in the tombs of high-ranking women, as suggested by the presence of very ornate jewellery. The necklaces were made of polychrome glass beads, glass paste, and amber. Earrings were closed-loop and varied in size; the rings often featured decorative settings.

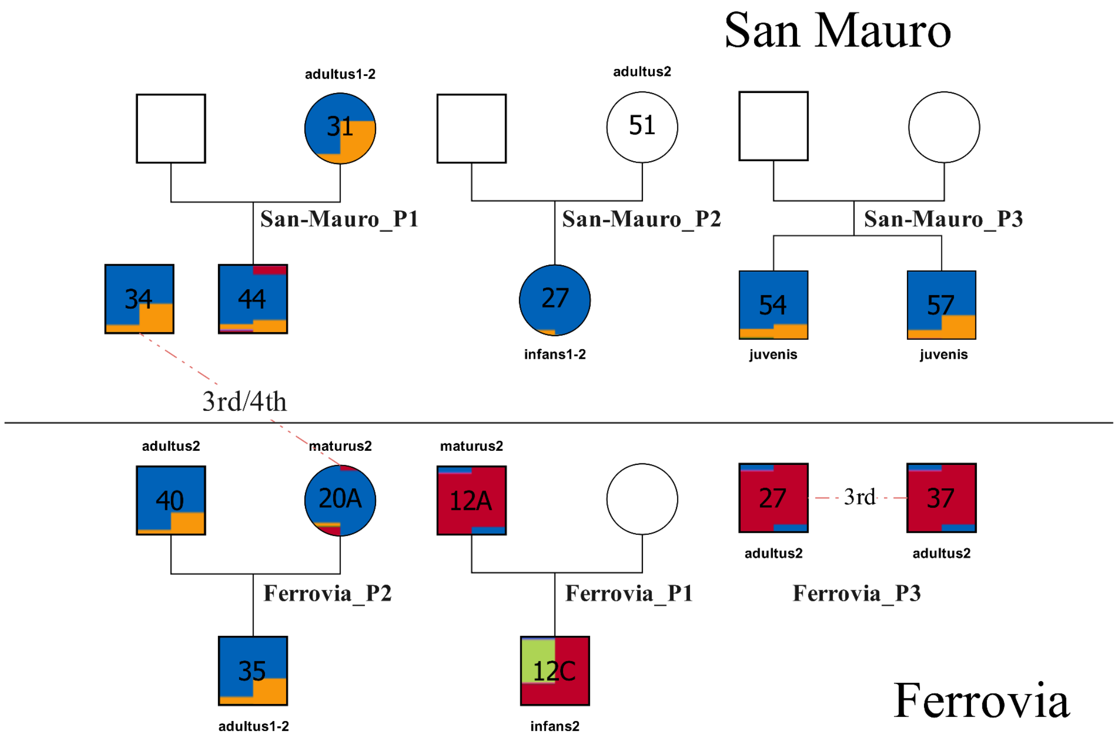


**Figure S17: Pedigrees found at San Mauro and Ferrovia. Squares indicate males, while circles indicate females. Individuals are colored based on their fastNGSadmix results, with the modern panel results on the left and the ancient panel results on the right (Extended Data Fig. 1). The connection PO represents a parent offspring pair where the polarity is unknown. Unknown relationships are represented using dashed lines labeled with the degree.**

Ahumada Silva I., 2010, “La collina di San Mauro a Cividale del Friuli. Dalla necropoli longobarda alla chiesetta bassomedievale”, All’Insegna del Giglio.

#### S2 Strontium ($^{87}Sr/^{86}Sr$) isotope analysis

As the impact of various types of mobility is one of the main topics of our research, we complemented the aDNA analysis with $^{87}Sr/^{86}Sr$ measurements. Similarly to the aDNA sampling, we aimed for comprehensive sampling of sites, but it was limited by preservation and availability of the osteological material. Altogether, we collected 329 samples from 20 sites for $^{87}Sr/^{86}Sr$ analysis.

Strontium (Sr) is a trace element with isotope ratios that vary across different geological formations. These ratios are introduced into human and animal tissues through the food chain, with Sr ratios in plants reflecting the bedrock and groundwater of the regions where they grow. While Sr is incorporated into all skeletal tissues, only dense and resistant tooth enamel is suitable for analysis in inhumed individuals. At the same time, the post-depositional environment diagenetically alters Sr in bones. Strontium ($^{87}Sr/^{86}Sr$) isotope analysis of tooth enamel is a valuable tool for understanding where individuals obtained their food during the time of enamel formation (childhood and adolescence). This method can identify individuals buried in areas with geological characteristics different from those of their childhood environment (see for example Bentley 2006 for details). Because enamel does not change once formed (Hillson 1997), the Sr values obtained represent the time of enamel formation - childhood and adolescence - depending on the analysed tooth. When possible, we have sampled 2nd or 3rd permanent molar for Sr analyses. These teeth start to form at the age between 2 and 3 years, and 8 and 9 years, respectively, with crowns being complete around the age of 9 years, and 15 years, respectively (AlQahtani et al. 2010).

##### Defining 'local' Sr ranges

"Local" community baseline was created using double standard deviation (SD) of mean Sr values obtained from the enamel of the individuals buried at each site. A more conservative (wider) and stricter (narrow) baselines were created. The conservative approach was based on the double SD values of all the individuals analysed from each site, while the strict was based only on non-adults buried at each site. In case of sites with less than two subadults, only a conservative approach was applied.

Individuals falling outside the community SD baseline were identified as outliers, who likely spent their childhood/adolescent years elsewhere (obtained food from a different geological area) and moved to the area where they were buried later.

Mean values with lower and upper bound based on 2SD for each site based on all the individuals are presented in Table S2-1.

**Table S2-1**. Mean Sr values with SD, lower and upper band for all individuals and subadults alone.

| **All individuals included** | | | | | |
|---|---|---|---|---|---|
| Site | Region | Mean | SD | Lower bound | Upper bound |
| Castra | Vipava valley | 0.7092 | 0.001 | 0.7068 | 0.7116 |
| Celje | Celje | 0.7098 | 0.001 | 0.7078 | 0.7118 |
| Dravlje | Ljubljana | 0.7106 | 0.001 | 0.7084 | 0.7128 |
| Gojače | Vipava valley | 0.7092 | 3E-04 | 0.7086 | 0.7098 |
| Gosposvetska | Ljubljana | 0.7102 | 0.001 | 0.7076 | 0.7128 |
| Hrušica | Ljubljana | 0.7117 | 0.002 | 0.7083 | 0.7151 |

| | | | | | |
|---|---|---|---|---|---|
| Ledine | Vipava valley | 0.7095 | 5E-04 | 0.7085 | 0.7105 |
| Miren | Vipava valley | 0.7092 | 5E-04 | 0.7082 | 0.7102 |
| Rifnik | Celje | 0.7098 | 8E-04 | 0.7082 | 0.7114 |
| Solkan | Vipava valley | 0.709 | 3E-04 | 0.7084 | 0.7096 |
| Sv. Pavel | Vipava valley | 0.7092 | NA | NA | NA |
| Šturje | Vipava valley | 0.7093 | 5E-04 | 0.7083 | 0.7103 |
| Camberk | Dolenjska region | 0.711 | 0.0021 | 0.7101 | 0.7118 |
| Gradec | Dolenjska region | 0.7095 | 0.0003 | 0.7092 | 0.7097 |
| Vrajk | Dolenjska region | 0.7097 | 0.0003 | 0.7095 | 0.7099 |
| Črnomelj | Bela krajina | 0.7099 | 0.0007 | 0.7095 | 0.7102 |
| **Only non-adults included** | | | | | |
| Site | Region | Mean | SD | Lower bound | Upper bound |
| Castra | Vipava valley | 0.7087 | 3E-04 | 0.7082 | 0.7092 |
| Camberk | Dolenjska region | 0.7106 | 0.0012 | 0.7082 | 0.7129 |
| Celje | Celje | 0.7101 | 0.001 | 0.7076 | 0.7125 |
| Dravlje | Ljubljana | 0.7112 | 0.002 | 0.7081 | 0.7143 |
| Gosposvetska | Ljubljana | 0.7113 | 0.002 | 0.7066 | 0.7159 |
| Rifnik | Celje | 0.7097 | 3E-04 | 0.7092 | 0.7103 |
| Solkan | Vipava valley | 0.709 | 3E-04 | 0.7085 | 0.7095 |
| Šturje | Vipava valley | 0.709 | 1E-04 | 0.7089 | 0.7092 |

**Geological background and environmental samples**

The Slovenian landscape consists of a complex combination of metamorphic, igneous and sedimentary rocks. In simple terms, the central and northern parts of Slovenia are covered by clastic sedimentary rocks from the Quaternary period. In the northern and north-eastern parts of the country, Oligocene to mid-Miocene igneous rocks predominate, and metamorphic rocks from the Palaeozoic are also rarely present. In the north-western and southern parts of the country, limestone and dolomite from the Mesozoic and Palaeocene are most common. Southwestern Slovenia consists mainly of Palaeogene flysch rocks, while central and eastern Slovenia consists of Neogene clastic sedimentary rocks (Buser 2010; Javornik et al. 1989).

Modern environmental $^{87}Sr/^{86}Sr$ data is available from truffles sampled in central and southern Slovenia and from milk from various regions of Slovenia. The determined $^{87}Sr/^{86}Sr$ values of truffles range from 0.7086 in the west (Sežana) to 0.7138 in the south (Bloke). In the central part of Slovenia, truffles from Quaternary sediments show values of 0.7121 (a sample from Meje) (Hamzić Gregorčič, Strojnik, et al., 2020). Milk ranges from 0.7085 to 0.7123, depending on the geological background. In the central and western part of Slovenia, the values are mostly between 0.7080 and 0.7090, while in the north-eastern part they are higher and range between 0.7100 and 0.7120 (Hamzić Gregoričič et al. 2021).

In addition, there is data for $^{87}Sr/^{86}Sr$ from water samples from springs in the Ljubljana basin, where the values are between 0.7081 and 0.7088 (Lojen et al. 2019).

**Results**

A general comparison of the human values obtained from the four regions (Celje, Ljubljana, Vipavska dolina, Bela krajina and Dolenjska region) shows the lowest values for $^{87}Sr/^{86}Sr$ in Vipavska dolina (mean=0.7091, SD=0.0004), followed by Celje (mean=0.7098, SD=0.0009), Bela Krajina (mean=0.7099, SD=0.0007), Dolenjska (mean=0.7100, SD=0.0006) and Ljubljana (mean= 0.7099, SD=0.0013). However, there are numerous quartile-based outliers in all the regions with Ljubljana also presenting a broader range (Figure S18).

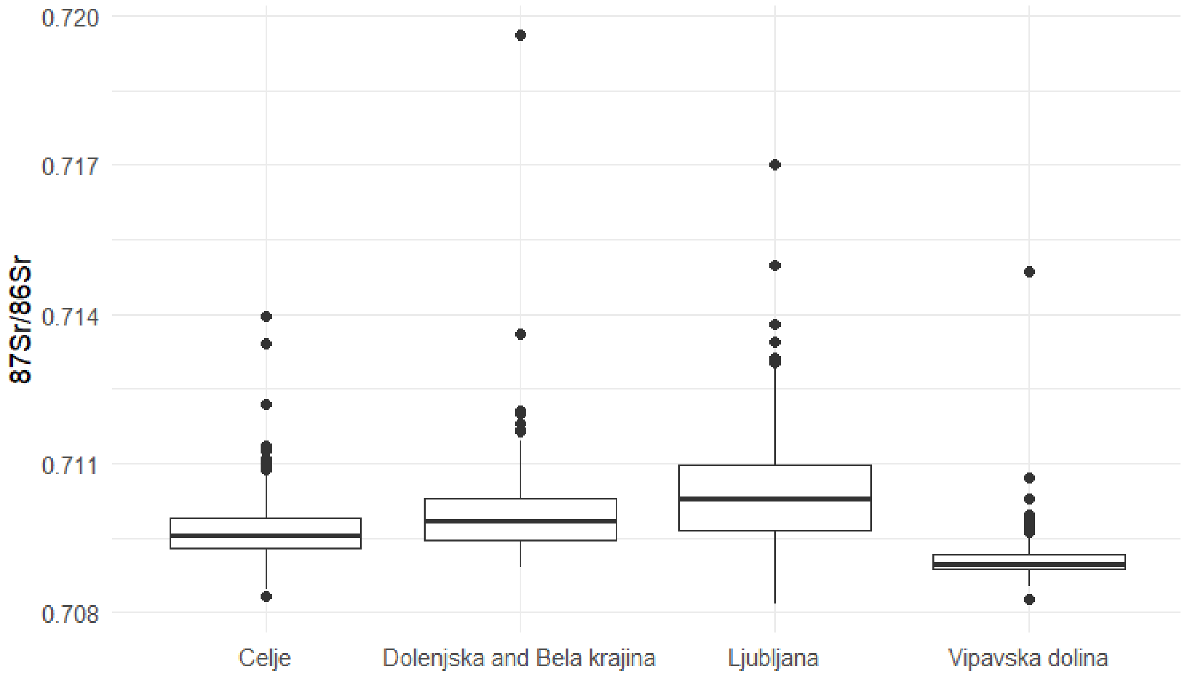


Figure S18. Comparison of $^{87}Sr/^{86}Sr$ values across Regions.

#### Regional analysis

##### Ljubljana region

Three sites from the central part of Slovenia were included: Dravlje, Gosposvetska and Hrušica, all located in present-day Ljubljana (Figure S19). The area consists of the Ljubljana Basin and the Ljubljana Marshes, which are surrounded by hills. The Ljubljana Basin consists of fluvioglacial gravels in which the Sava River has created numerous terraces, while the Ljubljana Marshes consist of Quaternary sediments, some of which were formed by waterlogging. A few low hills that can be found here consist of Mesozoic and Paleozoic sediments. The Ljubljana basin is surrounded by hills consisting mainly of Permo-Carboniferous, Permian and Triassic clay shale, sandstone, limestone and dolomite.

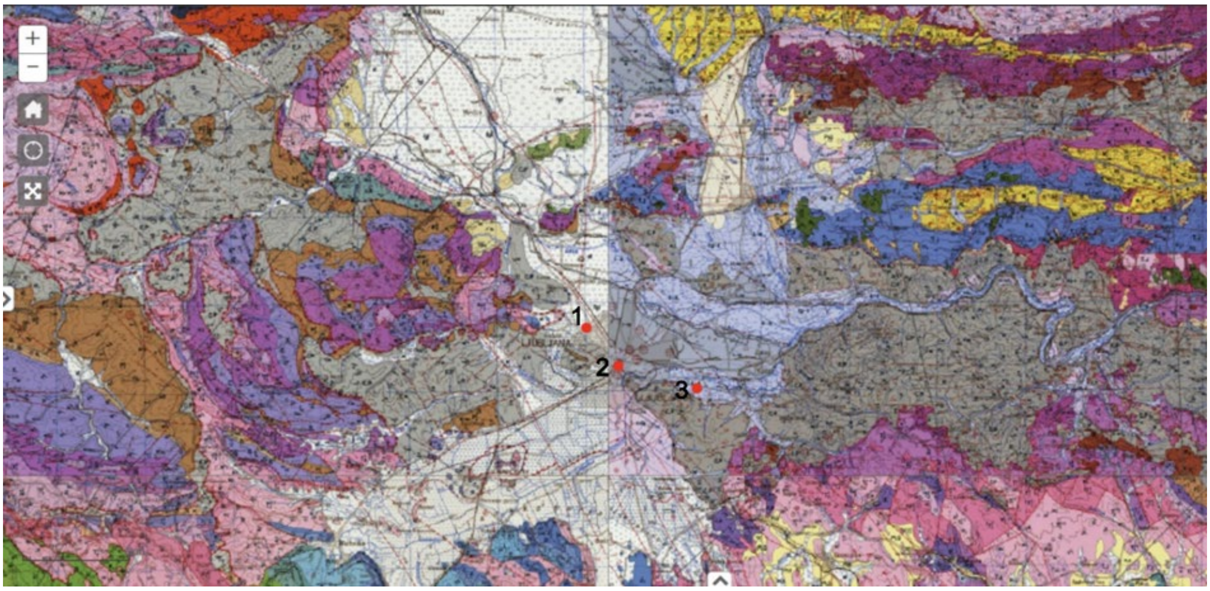


Figure S19. Geological map of Ljubljana and surroundings with location of archaeological sites Dravlje (1), Gosposvetska (2) and Hrušica (3) (Source: https://ogk100.geo-zs.si).

The only published data on $^{87}Sr/^{86}Sr$ values from an archaeological context for the area come from a Late Bronze Age/Early Iron Age cremation cemetery in Ljubljana (SAZU) (Škvor Jernejčič & Price, 2020) and part of the Gosposvetska site included here (Leskovar et al. 2025). The reported values for cremated human remains from SAZU range from 0.7093 to 0.7123, with a mean of 0.7110. The values for remains from the previously published Gosposvetska range from 0.7091 to 0.7134, with a mean of 0.7102, and from Gosposvetska as a whole, old and new data, from 0.7082 to 0.7170, with a mean of 0.7102.

The comparison of the data sets from Dravlje, Hrušica and Gosposvetska (Figure S20) shows a high but mainly overlapping variability in the Gosposvetska and Dravlje sites, with most individuals showing values between 0.7090 and 0.7110 and three obvious outliers. Hrušica presents a wide range, only partially (lower values) overlapping with Gosposvetska and Dravlje

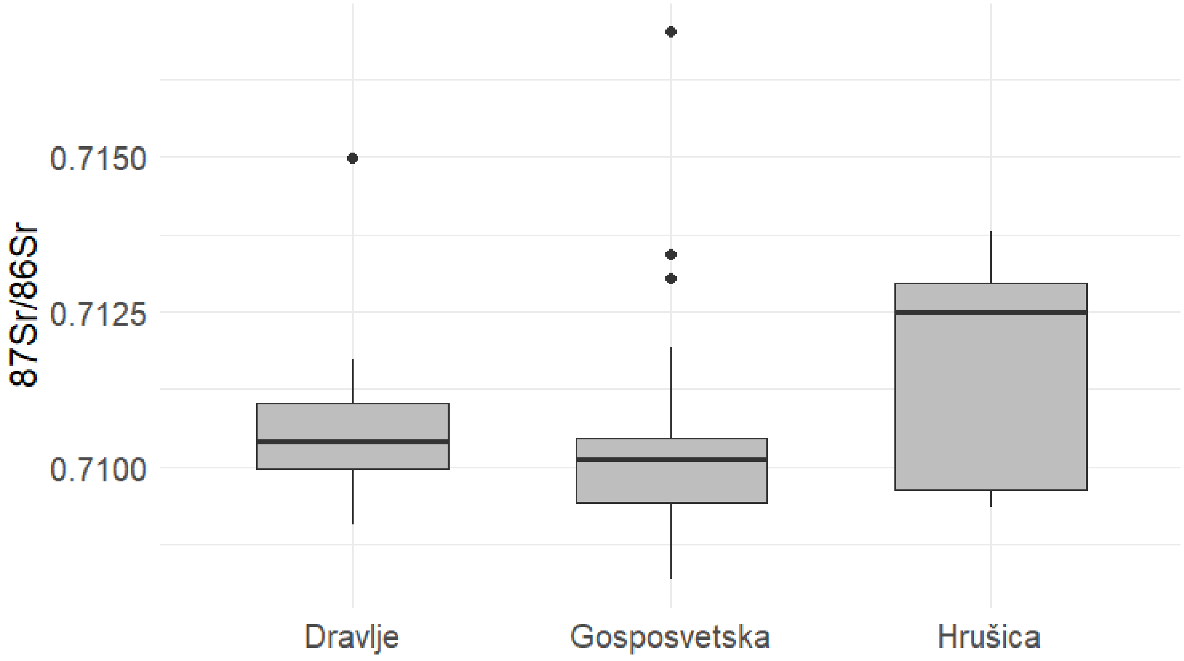

Figure S20. Boxplots of $^{87}Sr/^{86}Sr$ values of Ljubljana region sites.

In Dravlje (#29), the $^{87}Sr/^{86}Sr$ values range from 0.7091 to 0.7150 with a mean of 0.7106 ± 0.0011. Based on the Mann-Whitney U-test, there are statistically significant differences between adults and subadults (p=0.01), while no further differences can be found when comparing different age classes. There are no significant differences between adult males and females (Mann-Whitey U-test, p=0.18). However, females show a lower mean value of $^{87}Sr/^{86}Sr$, with only one female reaching the mean value of 0.7103 of the males.

The juvenile female from grave 34 with a value of 0.7150 is an outlier. Apart from outliers, such high values are not recorded at any other Slovenian site, whether modern or archaeological.

At Hrušica (#12), the $^{87}Sr/^{86}Sr$ values range from 0.7093 to 0.7138 with a mean of 0.7117 ± 0.0016. There do not appear to be any outliers at this site, although the range is quite wide. Based on the Mann-Whitney U test, there are no statistically significant differences between adults and subadults (p=0.148). However, there is only one subadult in the data set, a one-year-old child with the highest $^{87}Sr/^{86}Sr$ values (0.7138).

There are no significant differences between adult males and females (Mann-Whiteny U-test; p=0.777), but two distinct groups with two males and two females with low $^{87}SrS/^{86}Sr$ values of 0.7093 – 0.7096 and two females and five males with high values of 0.7119 – 0.7131. Compared to the rest of the Ljubljana sites, the lower group fits the “community” signal better, while the individuals in the group with high values, adults with sampled first (two individuals) or second (five individuals) molars, probably received food during their childhood and/or lived in a different area.

In Gosposvetska (#21 previously published + #42 new) the $^{87}Sr/^{86}Sr$ values range from 0.7082 to 0.7170 with a mean of 0.7102 ± 0.0012. There are three outliers in the data with higher values, one adult female with a value of 0.7134 and two juvenile females with values of 0.7131 and 0.7170. Based on the Mann-Whiteny U-test, there are statistically significant differences between adults and subadults (p=0.02). On average, non-adults, especially from age class infans II, but also from infans I, present higher $^{87}Sr/^{86}Sr$ values. There are no statistically significant differences between adult males and females (Mann-Whitney U test; p=0.94). There are three outliers at Gosposvetska, approximately 5-year-old child, juvenile (approximately 14 years old) and young adult female (from graves 3103, 9016 and 3036, respectively).

**Celje region**

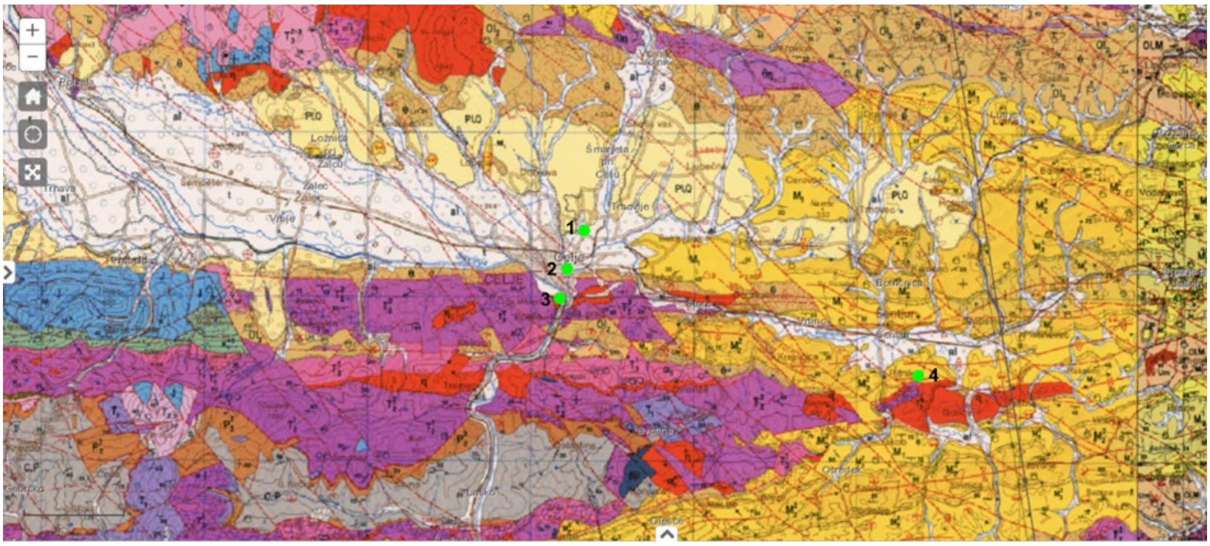


Figure S21. Geological map of Celje and surroundings with location of archaeological sites Mariborska cesta (1), CeleiaPark (2), Breg (3), Rifnik (4) (Source: https://ogk100.geo-zs.si).

Three locations from the Celje region were included: Mariborska cesta, Celeiapark and Breg in present-day Celje, and Rifnik, located 10 km outside Celje to the southeast (Figure S21). The first three sites are located in the Celje basin, surrounded by the Posavje hills to the south, the Menina and Dobrovlje karst plateaus to the west, the Vitanje-Konjice Karavanke hills to the north, the Pohorje hills to the northeast and the Voglajna hills to the east. Rifnik is a 568 m high mountain that belongs to the Posavje Hills.

Geologically, the Celje basin is a young tectonic depression filled with Quaternary carbonate deposits from the Savinja River and non-carbonate alluvial deposits from its tributaries, the Voglajna and Hudinja rivers. In the northern part there are also Plio-Pleistocene clays with intercalations of acid tuff. Andesitic tuffs from the Oligocene are exposed directly on the southern side of the Celje Basin. Similar formations are also found in the northern and eastern parts. The oldest rocks are part of the Posavje folds, where Rifnik is located, and outcrops at the southern end. These include pseudo-Zilje deposits, limestone, keratophyre and tuff from the Ladino age as well as massive limestone from the Upper Triassic (Bauser 1997).

There is no published data on $^{87}Sr/^{86}Sr$ values from archaeological contexts from the region, so the data presented here are the first. The only published data from the region around Celje and Rifnik are $^{87}Sr/^{86}Sr$ values from present-day milk (Hamzić Gregorič et al. 2021). The values from the dairy farms located 15 – 30 km west and northeast of Celje (Polzela, Ponikva and Slovenske konjice) show values of 0.7091 – 0.7110.

In Breg (#12), $^{87}Sr/^{86}Sr$ values range from 0.7087 to 0.7113 with a mean of 0.7097 ± 0.0008. Two adult females with higher values (Breg 20, 0.7111, and Breg 1U, 0.7113) are considered outliers. Based on the Mann-Whitney U-test, there are no statistically significant differences between adults and subadults (p=0.562), however, only one subadult is included in the data set. There are also no statistically significant differences between adult males and females (p=0.315), but the range for females is significantly larger compared to males, as the outliers are females. The outliers are only true outliers if only Breg is considered. Compared to the very nearby sites of CeleiaPark and Mariborska, they match the rest of the individuals from the Celje region (Figures S22 and S23).

At the CeleiaPark and Mariborska (#38) sites, the $^{87}Sr/^{86}Sr$ values range from 0.7084 to 0.7134 with a mean of 0.7099 ± 0.0011. Adult male (CP grave 10) and adult female (CP grave 12) from Celeiapark with higher values (0.7134 and 0.7114) are considered as outliers (Figures S225 and S236). According to the Mann-Whitney U-test, there are no statistically significant differences between adults and subadults (p=0.588). There are also no statistically significant differences between adult males and females (p=0.743).

In Rifnik (#37), the $^{87}Sr/^{86}Sr$ values range from 0.7092 to 0.7140 with a mean of 0.7098 ± 0.0008. Two adults, one male and one female, with higher values (0.7109 and 0.7140, respectively) are considered outliers (Figures S225 and S236). According to the Mann-Whitney U-test, there are no statistically significant differences between adults and subadults (p=0.216). There are also no statistically significant differences between adult males and females (p=0.970).

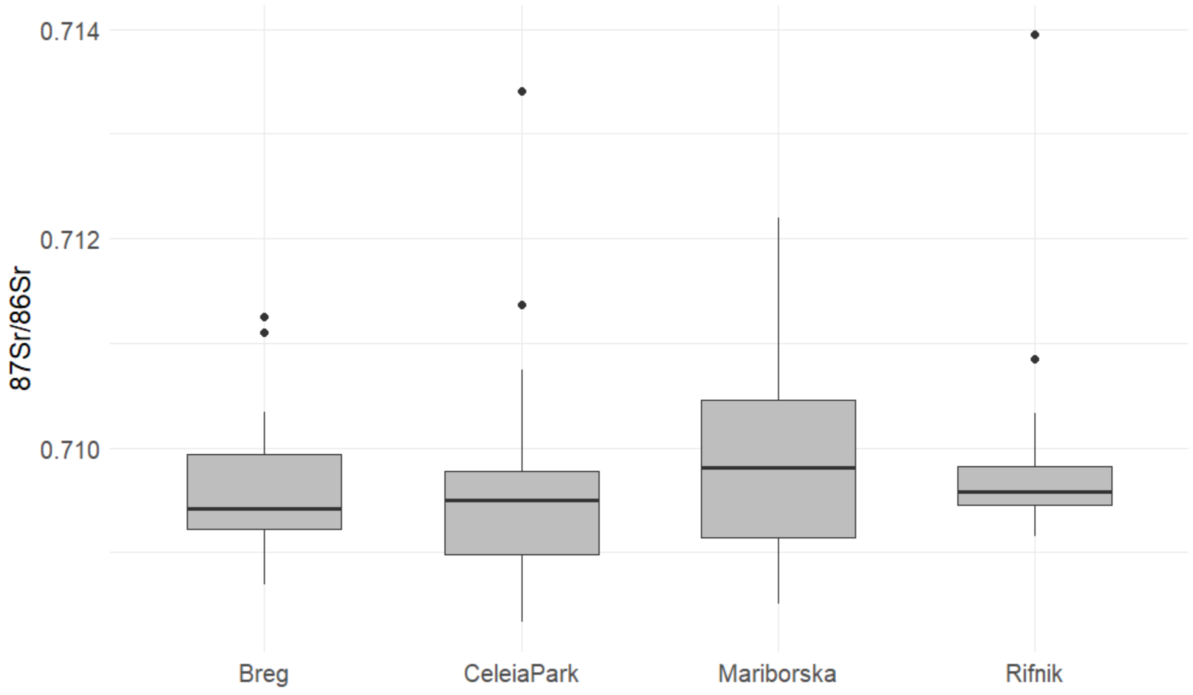


Figure S22. Boxplots of $^{87}Sr/^{86}Sr$ values of Celje region sites.

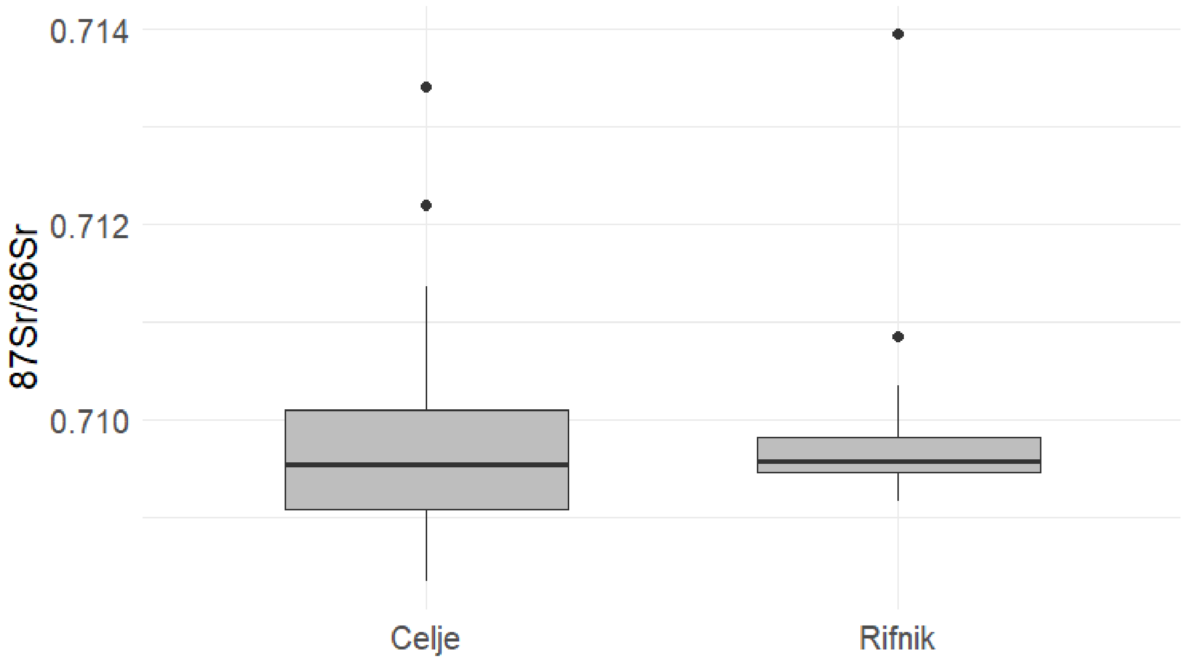


Figure S23. Boxplots of $^{87}Sr/^{86}Sr$ values of Celje region sites with merged cemeteries of Breg, Mariborska and CeleiaPark.

**Vipava valley region**

Sites from Vipava valley region are scattered across the region and include Ajdovščina Castra and Ajdovščina Šturje in the northeastern part of the valley, Vrtovin and Gojače in the central part of the valley, Solkan and Ledine in the northwestern part of the valley and Miren in the southwestern part of the valley (Figure S24). The Vipava valley is bordered by the plateaus Nanos, Hrušica, and Trnovski gozd from the east to the west, while to the south, it is separated from the Karst by the Vipava Hills. It is separated from the Adriatic coast by a karst plateau that is 300-400 meters high and 12 kilometres wide. The valley is characterized by considerable relief variation, as the altitudes range from 60 meters (the Vipava riverbed near Batuje) to 1,495 meters (Mali Golak) (Pak 1987, 29).

Vipava Valley is a flysch syncline, which was compressed by tectonic pressure between the two harder limestone blocks: Trnovski gozd with Hrušica to the north and the Karst to the south (Slapernik 1994, 13). It is composed of a Eocene flysch, covering as much as three-fifths of its surface area. On the northern side of the valley, steep slopes are covered with Cretaceous limestone. On the southern side of the valley, flysch layers rise towards the Karst region, and in some places, they even overlay it (Kladnik, Natek, 1998, 222).

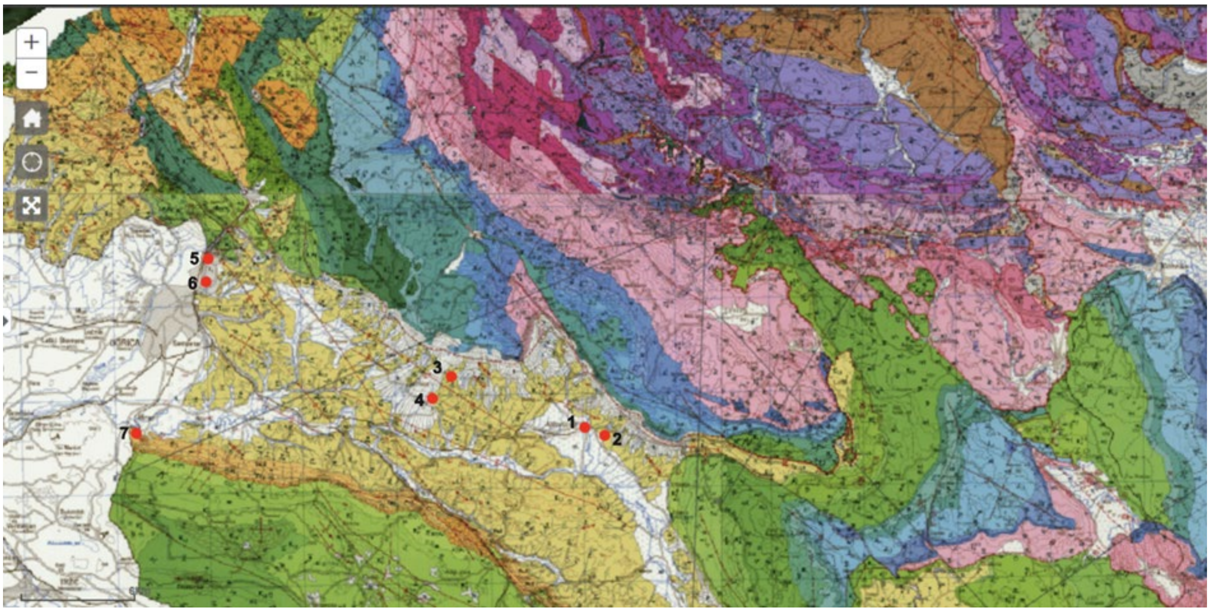


Figure S24. Geological map of Vipava valley with location of archaeological sites Ajdovščina-Castra (1), Ajdovščina-Šturje (2), Vrtovin (3), Gojače (4), Solkan (5), Ledine (6), Miren (7) (Source: https://ogk100.geo-zs.si).

There are no previously published data on $^{87}Sr/^{86}Sr$ values from archaeological or modern context from the Vipava valley. The obtained values range from 0.7083 to 0.7149, with a mean value of 0.7092 ± 0.0008. All the sites present a narrow range, with outliers identified in Castra, Šturje, Ledine and Solkan.  Except from Miren, there is a pair of sites located close by in the Vipava region. Thus, first comparisons between these pairs were made, also identifying outliers. Further adult - subadult and male - female statistical comparisons were only performed for Castra, Šturje and Solkan, as other sites have very low numbers of individuals.

Ajdovščina sites (Castra #25 and Šturje #18), located 0.5 km apart, present no significant differences (t-test; p=0.733) in mean $^{87}Sr/^{86}Sr$ values. Excluding outliers, both sites also present a narrow range of values from 0.7086 to 0.7095. There are four outliers in Castra; two individuals, young adult male (grave 18, 0.7096) and mature female (grave 14; 0.7098) present slightly higher values, which can be seen at other sites in the Vipava valley. More obvious strontium outliers are a young adult male (grave 13) with very high values (0.7149) and young adult female (grave 30) with low values (0.7083), not observed at other sites in the region. Based on the Mann-Whitney U test there are no statistically significant differences between adults and subadults (p=0.706) in Castra (#25). There are also no statistically significant differences between adult males and females (p=0.186). In Šturje, there are two outliers, young adult male (grave 3) and female (grave 18) with values (0.7103 and 0.7107, respectively) higher than observed at other sites in Vipava valley. Based on the Mann-Whitney U test there are no statistically significant differences between adults and subadults (p=0.143) in Šturje (#18). There are also no statistically significant differences between adult males and females (p=0.868).

Two small sites, Gojače – Boršt (#2) and Sv. Pavel (#1) are located close together, and the observed values overlap completely within the range of 0.7086 – 0.7098. There are no outliers.

Miren (#6) presents a wide range of $^{87}Sr/^{86}Sr$ values from 0.7082 to 0.7102. There are no outliers.

Ledine (#5) and Solkan (#25), both located in modern day Nova Gorica (1.5 km apart) present different $^{87}Sr/^{86}Sr$ values. Mean value for Ledine is 0.7095 ± 0.0005 while for Solkan 0.7090 ± 0.0003, indicating at least partially different origin of food. Solkan, located more towards the north, and nearby slopes of Trnovski gozd, is closer to a Cretaceous limestone block that borders on flysch of Vipava valley. If food was obtained from the areas of limestone, it could explain the lower values detected in Solkan. Though modern milk and truffle samples were not measured from any nearby location, the values of samples from Cretaceous Limestone present a mean value of 0.7091 ± 0.0001 (Hamzić Gregorčič et al. 2021), which agrees with lower values in Solkan. Based on the Mann-Whitey U test there are no statistically significant differences between adults and subadults (p=0.426) in Solkan. There are also no statistically significant differences between adult males and females (p=0.583). Though the range in females is wider due to the outliers being females. In Ledine, there is one outlier, an adult female (grave 4), with low values (0.7086). In Solkan, there are three outliers, two adult females (grave 6 and 53) with high $^{87}Sr/^{86}Sr$ values (0.7099 and 0.7100, respectively), and an adult female (grave 50) with low value (0.7085).

Based on the Mann-Whitney U test there are no statistically significant differences between adults and subadults (p=0.558) in Miren (#6), through the only subadult, the infant from grave 8 is on the lower end of the range. There are also no statistically significant differences between adult males and females (p=1).

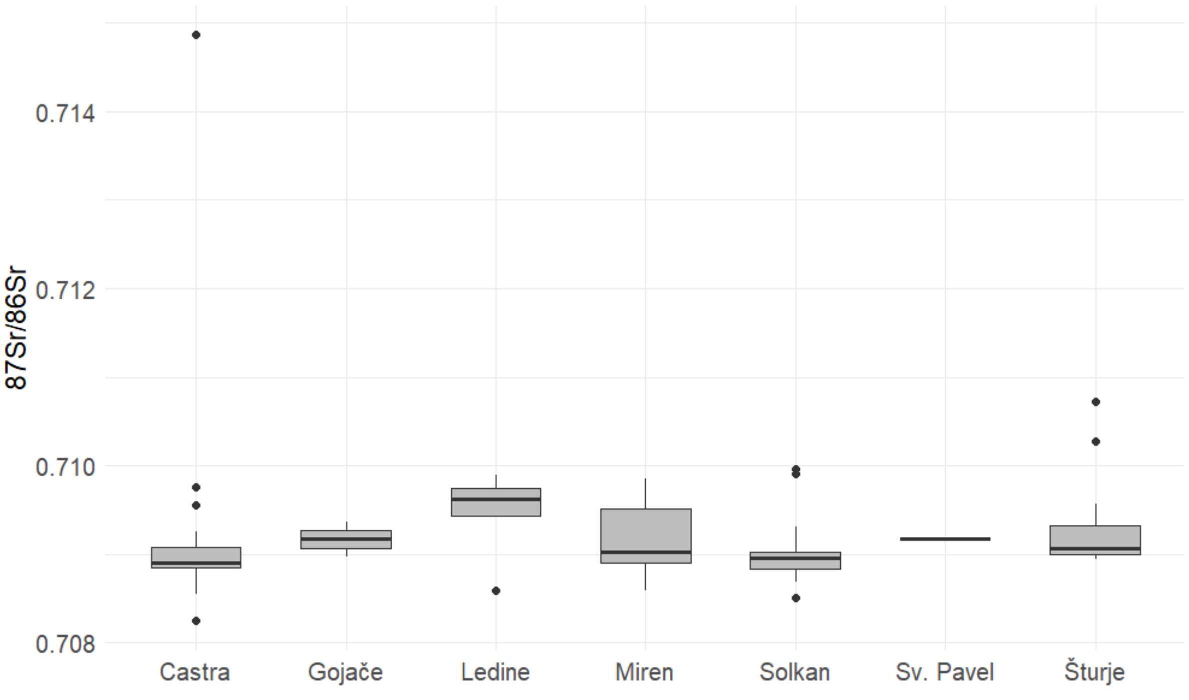


Figure S25. Boxplots of $^{87}Sr/^{86}Sr$ values of Vipava valley sites.

#### Dolenjska and Bela krajina region

The geological structure of the Novo Mesto region is composed almost entirely of sedimentary rocks, mainly from the Permo-Carboniferous period. These include quartz sandstones, conglomerates, and clay shales, some of which are found in the Žumberak area. Triassic formations are widespread, featuring layers of dolomite, limestone, marl, and anhydrite, with fossil finds such as foraminifera and bivalves. Ladinian layers (Middle Triassic) are notable for their composition of dolomite, limestone, and marls with marine fossils. The Jurassic period in this region is mostly represented by dolomites and limestones with rich fossil assemblages, including algae and stromatolites. Cretaceous formations appear mainly as marl-limestone series, rich in macro- and microfossils, particularly around the Gorjanci Hills and Žumberak. Tertiary (Miocene) deposits include marl and sandy marl with foraminifera and mollusks, while Quaternary deposits include alluvial sediments from river terraces and valleys. The area also exhibits tectonic features typical of the Dinaric system, with numerous faults and folds influencing the structure and stratigraphy (Pleničar, Premru 1970).

The Bela krajina area features a complex geological structure shaped by varied sedimentation phases and tectonic movements. It includes three main sedimentary complexes: Vrbovsko-Knežja Lipa with reduced Triassic deposits, Črnomelj-Bosiljevo with a full Mesozoic sequence, and Zvečačko-Metlika with reduced Jurassic and Lower Cretaceous layers. Sediments range from Paleozoic sandstones and limestones to richly fossiliferous Jurassic and Cretaceous limestones, often interbedded with dolomites, marls, and breccias. Tectonic activity, including nappe structures and faulting, strongly influenced deposition and preservation. The area contains economically relevant resources such as bauxite, clay, lignite, quartz sand, and building stone, with active exploitation of limestone and dolomite (Bukovac, Poljak, Šušnjar, Čakalo 1983).

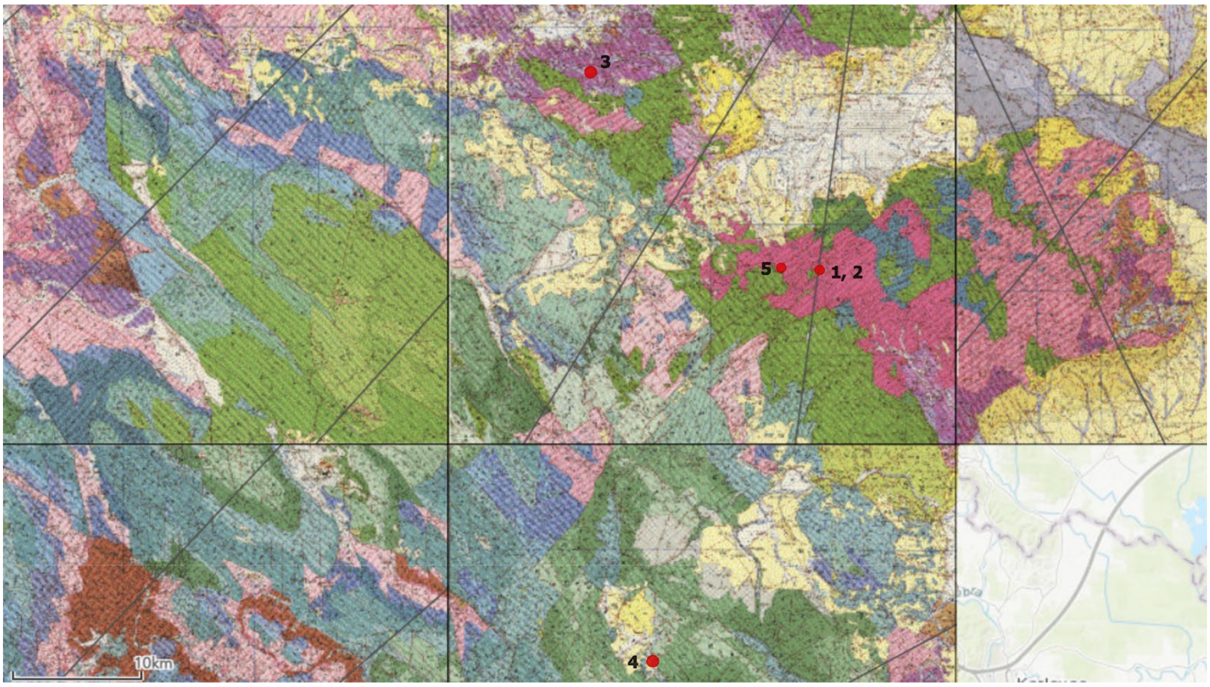


Figure S26. Geological map of Dolenjska and Bela krajina region with location of archaeological sites Gradec above Mihovo (1), Zidani Gaber (2), Vrajk (3), Črnomelj (4), Camberk (5) (Source: https://ogk100.geo-zs.si).

There is no previously published data on $^{87}Sr/^{86}Sr$ values from archaeological or context from the Dolenjska and Bela krajina. The only modern data that exist are truffles from Rajndol located 20 km East from Črnomelj with $^{87}Sr/^{86}Sr$ values of 0.7115, milk from Vinica located 15 km to the south of Črnomelj with $^{87}Sr/^{86}Sr$ values of 0.7095, milk from Ždinja vas near Šmarješke Toplice 8 km south of Vrajk and 15 km northwest from Camberk and Gradec with $^{87}Sr/^{86}Sr$ values of 0.7094. The values from here included archaeological sites range from 0.7088 to 0.7112, with a mean value of 0.7100 ± 0.0006. All the sites but Camberk present a narrow range, with outliers identified in Črnomelj and Camberk (Figure S27).

Zidani gaber (#1) and Gradec (#6), both located in the Gorjanci hills, present a narrow range of $^{87}Sr/^{86}Sr$ values from 0.7091 to 0.7097. There are no outliers. 1.5 km distant Camberk presents significantly wider range of $^{87}Sr/^{86}Sr$ values from 0.7092 to 0.7196 with mean value of 0.7110 ± 0.0021 and one outlier, adult male from grave 32, with very high value of 0.7196. Based on the Mann-Whitney U test there are no statistically significant differences between adults and subadults (p=0.69) and between adult males and females (p=0.52).

Vrajk (#15) located 20 km to the north, presents a narrow range of $^{87}Sr/^{86}Sr$ values with a mean of 0.7097 ± 0.0003. Based on the Mann-Whitney U test there are no statistically significant differences between adults and subadults (p=0.56) and between adult males and females (p=0.40).

Črnomelj (#15) in Bela krajina also present a narrow range of $^{87}Sr/^{86}Sr$ values with a mean of 0.7099 ± 0.0007. There is one outlier in Črnomelj, an adult male from grave 23, with a high value of 0.7116. Based on the Mann-Whitney U test there are no statistically significant differences between adults and subadults (p=0.076) and between adult males and females (p=0.93).

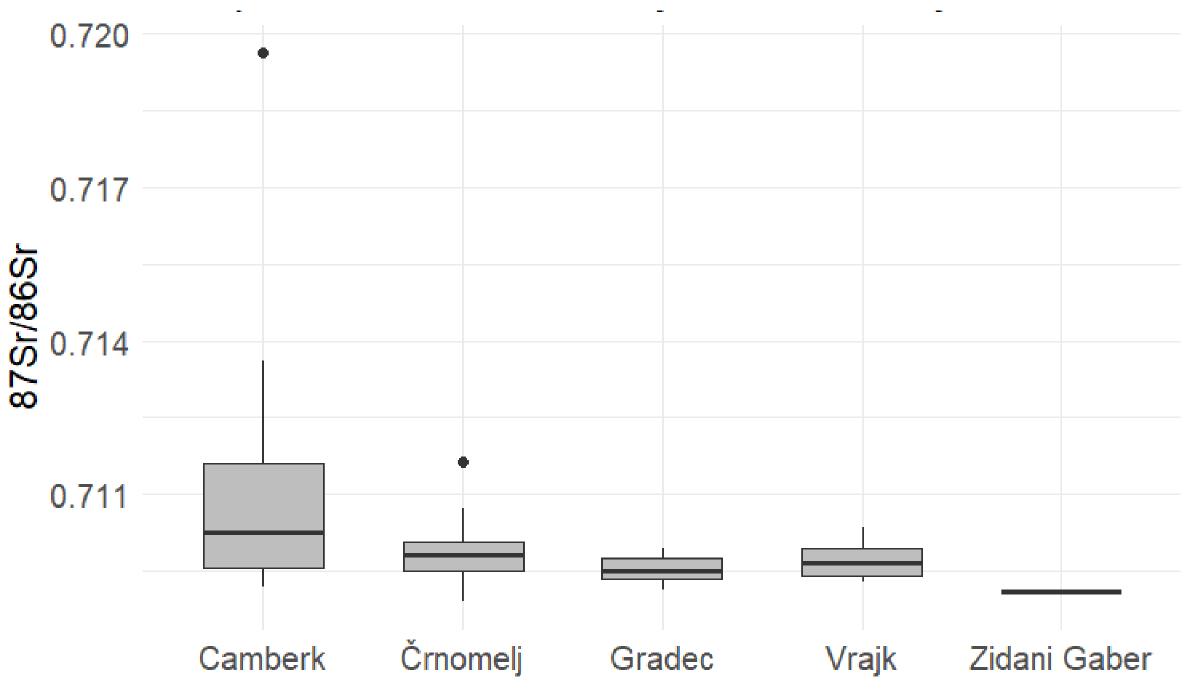


Figure S27. Boxplots of $^{87}Sr/^{86}Sr$ values of Vipava valley sites.

#### S3 Assessing community continuity using qpWave and qpAdm

In order to test if communities between later time periods could be modeled based on communities from earlier time periods, we used the programs qpWave[1,2] and qpAdm[2]. For these analysis, all individuals were called using an in-house random read genotype caller using an indent of 8 to generate pseudohaploid calls (https://github.com/kveeramah/) consistent with Vyas, Koncz, et al. (2023)[3].

These programs can test if the ancestry in a target individual (or population) is consistent with deriving from a given source population. It uses reference "right" populations as controls, alongside a target population. We used five right populations, consisting of Anatolian_Neolithic (n = 26), Steppe_Eneolithic (n = 18), Western Hunter-Gatherer (WHG) (n = 15), Iran_Neolithic (n = 9), and Morocco_Iberomaurusian (n = 4) (Table S2)[2,4–14]. For source populations, we used published European individuals (Table S2) from the time period of the target and earlier, as well as three reference Asian groups (Central Asian, East Asian, South Asian) ranging from around the 3rd to 10th centuries CE.

We used each of the individuals from our Slovenian dataset as target individuals (only excluding those with elevated contamination or less than 0.1× coverage of autosomal 1240k SNP dataset). Sites with only one or two sample individuals (Zidani gaber, Vrtovin, and Gojače) were excluded from plots or extensive analyses due to sample size.

To test for continuity with what is now Slovenia, we tested whether each LA or EM individual in our dataset could be modeled as deriving from one of the Slovenian communities from a previous time period. As our oldest data are from the LR period, we were not able to do this type of analysis of LR data. Individuals in within-site pedigrees were pruned when constructing source populations. To construct European source populations, for each time period (LR, LA, and EM), we randomly sampled 10 individuals from regions present in our dataset (Balkans, Iberia, Peninsular Italy, Northwest Italy, Austria, Bavaria, Caucasus, France, Northern Germany, Northeast Europe, Anglo-Saxons, and Scandinavia).

It must be noted that the source population reference panels vary between time periods, as we only use individuals penecontemporaneous or older than the target population for European source populations, this is to say, for example, the individuals used in the Balkan_LA panel will vary from the individuals used in the Balkan_LR and Balkan_EM panels. Due to limited sample size, for the three Asian groups (Central Asian, East Asian, South Asian), we used the same reference group for each time period

In certain cases, we also tested individuals against communities within their same broader time frame. For example, Solkan, Rifnik, Miren, and Dravlje are all treated as LA communities; however, Solkan and Rifnik post-date Miren and Dravlje and thus we conducted analyses with target individuals from Solkan/Rifnik and Miren or Dravlje as the source (Figure 3).

In other analyses, we found that at the sites of Miren and Dravlje that there was the presence of Central Asian admixture. In qpWave, we found that none of the individuals from Miren and most of the individuals from Dravlje could not be modeled by any of the LR communities (n=23). For these 23 individuals (LJ-Dravlje_01, 16, 17, 18, 19, 20, 22, 23, 24, 25, 33, 34, 38, 42, 44, 47, and 49 as well as six individuals from Miren), we conducted qpAdm analyses using the three Asian references as a possible second source along side of one of the LR communities. We find that in 11 of the 23 individuals, they were able to be modeled using at least one LR community in conjunction with the Central Asian panel, while the other 12 remained without any feasible models with tail probabilities greater than 0.05;. Additionally from the 11, 6 could be modeled by the South Asia panel and 3 by the East Asian panel, in place of the Central Asia panel (Table S6).

#### S4 Factor Analysis

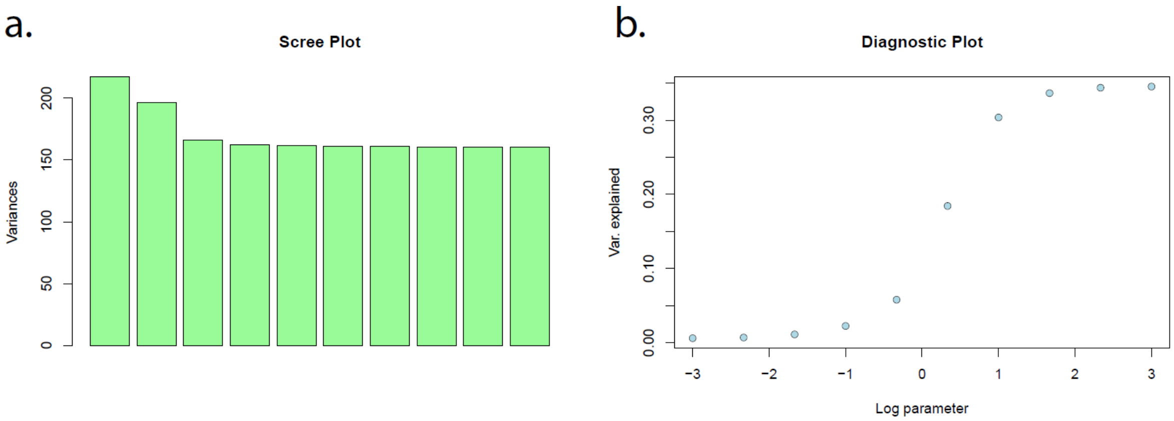


**Figure S27. a.** Scree plot showing the variance (eigenvalues) associated with each principal component for the imputed genotype dataset used in the factor analysis. **b.** Diagnostic plot from choose_lambda() showing the variance ratio as a function of log10(lamda) for k=2 when conducting factor analysis using the imputed genotype dataset.
